# Multiple routes to interspecific territoriality in sister species of North American perching birds

**DOI:** 10.1101/843516

**Authors:** Madeline C. Cowen, Jonathan P. Drury, Gregory F. Grether

## Abstract

Behavioral interference between species can influence a wide range of ecological and evolutionary processes. Here we test foundational hypotheses regarding the origins and maintenance of interspecific territoriality, and evaluate the role of interspecific territoriality and hybridization in shaping species distributions and transitions from parapatry to sympatry in sister species of North American perching birds (Passeriformes). We found that interspecific territoriality is pervasive among sympatric sister species pairs, and that interspecifically territorial species pairs have diverged more recently than sympatric non-interspecifically territorial pairs. None of the foundational hypotheses alone explain the observed patterns of interspecific territoriality, but our results support the idea that some cases of interspecific territoriality arise from misdirected intraspecific aggression while others are evolved responses to resource competition. The combination of interspecific territoriality and hybridization appears to be an unstable state associated with parapatry, while species that are interspecifically territorial and do not hybridize are able to achieve extensive fine- and coarse-scale breeding range overlap. In sum, these results suggest that interspecific territoriality has multiple origins and that interspecific territoriality and hybridization together can have striking impacts on species ranges.

## INTRODUCTION

Behavioral interference between species, such as interspecific courtship, mate guarding, or territorial defense, can have considerable impacts on the ecology and evolution of co-occurring species (Robinson and Terborgh 1995; Amarasekare 2002; Gröning and Hochkirch 2008; Grether et al. 2009, 2013; Kishi and Nakazawa 2013; Drury et al. 2015). Understanding the causes of different types of behavioral interference, their impacts on species coexistence, and the timescale over which they operate are thus active areas of research (Laiolo 2013; Martin and Ghalambor 2014; Losin et al. 2016; Grether et al. 2017; Kyogoku and Sota 2017; Sottas et al. 2018). Recent empirical and theoretical work has documented influences of interspecific territoriality on species coexistence and evolution in diverse taxonomic systems (reviewed in Grether et al. 2017). For instance, interspecific territoriality can facilitate species replacements (e.g., Duckworth and Badyaev 2007), accelerate competitive exclusion (e.g., Pasch et al. 2013), and foster coexistence between resource competitors that otherwise might not be expected to coexist (e.g., Ovadia and Dohna 2003; Ziv and Kotler 2003). While these findings highlight an important role for interspecific territoriality in fundamental ecological and evolutionary processes, general explanations for the occurrence, stability, and impacts of interspecific territoriality remain elusive.

Four sets of hypotheses provide possible explanations for interspecific territoriality. The *resource competition hypothesis* posits that interspecific territoriality persists due to resource competition and acts as a mechanism of spatial partitioning. In some cases, interspecific territoriality persists among resource competitors through adaptive convergence in territorial signals and/or competitor recognition (Cody 1969, 1973; Grether et al. 2009). Another hypothesis that assumes interspecific territoriality is adaptive when there is resource competition is that one species gains access to more resources through this behavior (MacArthur 1972). One pattern predicted by this *asymmetric competition hypothesis* is that interspecific territoriality is more likely to occur when one species is dominant in aggressive interactions. Third, local mate competition arising from reproductive interference (e.g., indiscriminate male mate recognition) could also make interspecific territorial defense adaptive and persist through time (Payne 1980; Drury et al. 2015). This *reproductive interference hypothesis* predicts a positive association between interspecific territoriality and indices of reproductive interference (e.g., rate of cross-species mating attempts, occurrence or frequency of hybridization). Fourth, if interspecific territoriality arises from misdirected intraspecific aggression, it should be transient and disappear over time as species evolve mechanisms to discriminate between heterospecifics and conspecifics (Murray 1971). However, it could persist if the species encounter each other too infrequently to evolve discriminatory mechanisms, or if hybridization prevents divergence (Murray 1971). We refer to this explanation for the persistence of interspecific territoriality as the *misdirected aggression hypothesis*.

Although interspecific territoriality has been documented in diverse two-species systems (e.g., Kral et al. 1988; Drury et al. 2015; Reif et al. 2015), to our knowledge, only one study has tested for a general explanation for interspecific territoriality across numerous taxa above the genus level (Losin et al. 2016). In North American representatives of the wood-warbler family (Passeriformes: Parulidae), Losin et al. (2016) found that interspecific territoriality is common, suggesting that this behavior is a more stable phenomenon than commonly assumed. They found that interspecific territoriality was positively associated with fine-scale habitat overlap (syntopy), supporting the resource competition hypothesis over the misdirected aggression hypothesis. Yet, wood-warblers are broadly ecologically similar (Lovette and Hochachka 2006), so to further evaluate the role of resource competition and other ecological circumstances in generating or maintaining interspecific territoriality, assessing these hypotheses in a dataset with greater ecological and phylogenetic diversity is key. Moreover, the diverse observed effects of interspecific territoriality on species coexistence (Ovadia and Dohna 2003; Ziv and Kotler 2003; Duckworth and Badyaev 2007; Pasch et al. 2013) raise the question of whether interspecific territoriality is adaptive for some species and maladaptive for others, or whether this behavior predominantly emerges and persists under one set of circumstances.

Characterizing the origins and persistence of interspecific territoriality is important for understanding not only how it manifests between interacting species, but also how it impacts their population dynamics. Research on species ranges suggests that competition or interference between species may impact range limits (Case et al. 2005; Price and Kirkpatrick 2009; Jankowski et al. 2010). In fact, evidence from sister taxa studies across vertebrate groups supports the hypothesis that becoming sympatric after allopatric speciation is constrained by ecological similarity or incomplete reproductive isolation (Price 2010; Weir and Price 2011; Pigot and Tobias 2013; Laiolo et al. 2017). While interspecific territoriality in some systems has led to competitive exclusion, it might also serve to increase alpha-diversity by enabling competing species to coexist (Robinson and Terborgh 1995; Grether et al. 2013; Grether et al. 2017); thus, the impact of interspecific territoriality on coexistence across breeding ranges remains unknown. If interspecific territoriality does affect the likelihood of two species coexisting, it might reduce the rate at which parapatric species transition into sympatry. Alternatively, interspecific territoriality might enable closely related species, strong resource competitors, and/or hybridizing species to transition more rapidly into sympatry than if they were not interspecifically territorial.

To address these knowledge gaps, here we examine interspecific territoriality in sister species of perching birds (order Passeriformes) that breed in North America, a group with a larger breadth of ecological and life history strategies than in any previous study of interspecific territoriality. First, we document the prevalence of interspecific territoriality across a large taxonomic group, spanning diverse ecologies and evolutionary histories. Second, we evaluate foundational hypotheses about the emergence and maintenance of interspecific territoriality, taking a step further than previous work by testing whether multiple hypotheses explain the observed pattern of interspecific territoriality. Third, we determine whether interspecific territoriality, alone and in combination with hybridization, contributes to regional coexistence and range expansion over evolutionary time.

Among the most recently diverged passerine birds in North America, we find support for the misdirected aggression and asymmetric competition hypotheses, suggesting that interspecific territoriality has multiple origins and evolutionary trajectories. Our work also identifies the potential for interspecific territoriality and reproductive interference to determine breeding range overlap between closely related species.

## METHODS

### Species pairs identification and classification

Our dataset consists of sister species of passerine birds that breed in North America and that overlap in breeding range. We identified sister species by sampling 10^4^ trees from the posterior distribution of a North American passerine phylogeny (Jetz et al. 2012) and selecting those that appeared as sister species in 90% or more of the phylogenies. Since allopatric sister species do not have the opportunity to be interspecifically territorial, we excluded species pairs that are allopatric in the breeding season according to 2016 and 2017 species distribution shapefiles from BirdLife International (www.birdlife.org). For each allopatric sister species pair, we selected the next most closely related species in the phylogeny that is sympatric with only one of the allopatric species to form a pair of closely related sympatric species. We only did this for one species from each allopatric pair to avoid sampling from non-independent nodes. We then created a maximum clade credibility tree from this posterior distribution in TreeAnnotator v1.8.4 (Suchard et al. 2018). Next, we calculated patristic distance between species from this phylogeny using the cophenetic.phylo function in the R package ape (Paradis et al. 2004). Due to recent taxonomic splits, we could not calculate patristic distance for all species pairs using this method. We obtained the patristic distance for one such pair, *Troglodytes pacificus* and *T. hiemalis*, from the literature (Toews and Irwin 2008). The other two species pairs that lacked patristic distances were omitted from our analyses.

We determined whether each species pair is interspecifically territorial with comprehensive literature searches using Web of Science, Birds of North America Online (Rodewald 2015), ProQuest Theses and Dissertations, and Google Scholar. We also contacted Birds of North America Online authors for additional behavioral observations. As in Losin et al. (2016), we considered a study sufficient evidence for interspecific territoriality if it contained at least two accounts of interspecific territorial aggression between unique individuals. Behaviors that qualified as interspecific territorial aggression include aggressive displays or countersinging, fighting, or chasing a heterospecific from a territory. We did not consider aggression over a food source or defense of a nest from a predator to be evidence of interspecific territoriality. Aggressive response to playbacks of territorial song and expansion of territory in response to removal of heterospecifics supported the classification of interspecific territoriality but were not required, since not all species pairs had been studied with these methods. If the behavior of both species in a pair had been studied together and no interspecific territoriality was reported, we classified that pair as non-interspecifically territorial. We omitted from our dataset any species pairs whose behavior had not been studied in sympatry (25 pairs), with two exceptions: the *Empidonax* species *E. difficilis* and *E. occidentalis* and the *Troglodytes* species *T. pacificus* and *T. hiemalis* have only recently been recognized as separate species (Johnson 1980; Toews and Irwin 2008), and have been reported to have non-overlapping territories in sympatry, so we classified them as interspecifically territorial. We also excluded species pairs for which neither species in the pair was intraspecifically territorial (2 species pairs), or for which we lacked data on fine-scale breeding habitat overlap (1 species pair). A full list of species pairs can be found in Table S1.

We classified species as hybridizing in the wild or not based on McCarthy (2006) and literature searches for newer reports of hybridization published in the years 2000 to 2018.

To assess whether greater study effort increased the likelihood of species pairs being reported as interspecifically territorial, we used the number of records of each species pair in the Zoological Records database (Thomson Reuters, New York, NY) as a proxy for past research and used Mann-Whitney tests to compare interspecifically versus non-interspecifically territorial species.

### Breeding range and habitat overlap quantification

We used two metrics to represent breeding season range overlap and habitat overlap of species pairs. First, we calculated the proportion of breeding range sympatry by dividing the area of overlap between BirdLife shapefiles by the breeding range area of the species with the smallest breeding range in each pair. However, BirdLife shapefiles were missing for two species pairs. We therefore also estimated sympatry using the Breeding Bird Survey (BBS; Sauer et al. 2017), a dataset of transects run across North America during the breeding season since the 1960s to survey the number of birds observed. Each BBS route is run annually, with 50 stops along each route. We measured sympatry by dividing the number of routes shared by both species by the total number of routes where the species with the fewest routes was observed. To replace the missing Birdlife sympatry values with rescaled BBS sympatry estimates, we used predicted values from a zero-intercept linear regression of the available Birdlife sympatry estimates on the BBS sympatry estimates (*R*^*2*^ = 0.69, df = 85, *P* < 0.0001).

Our second measure of overlap was syntopy (Rivas 1964), a fine-scale measure of breeding habitat overlap within the region of sympatry, such that species with higher syntopy are more likely to occur in the same habitat at the same time within their breeding range. We measured syntopy by identifying BBS routes where both species in a breeding season were found and dividing the number of “shared” stops (where both species were observed) by the number of stops where either species was observed. For two sympatric species pairs without BBS data (*Plectrophenax hyperboreus* and *Plectrophenax nivalis*; *Ammodramus caudacutus* and *Ammodramus nelsoni*), we used rescaled measures of syntopy from eBird records (Sullivan et al. 2009) (Supplement 1).

### Ecological trait quantification

To determine whether interspecific territoriality can be predicted by species-level traits, we collected ecomorphological data for each species and calculated the difference between these traits for each species pair. We focused on male traits since males perform territorial displays and defense for all territorial species in our dataset. We collected mass and bill length (exposed culmen length) values from the Birds of North America Online or additional references (e.g., Oberholser 1974, Dunning 2008). To account for possible geographic variation in the traits, when possible we used measurements collected close to the location where interspecific territoriality was studied. If the bill length measurement we found for a species was a measurement from the nostril to the tip of the bill instead of the exposed culmen length, we used a linear regression equation based on species for which both types of measurements were available (*R*^*2*^ = 0.985, df = 23, *P* < 0.0001) to predict exposed culmen length from the nostril-to-tip measurement.

We categorized foraging guild overlap between species in a pair by calculating the number of foraging guild axes on which the species overlap based on de Graaf et al. (1985). Specifically, species were categorized by the food types, foraging techniques, and foraging substrates used during the breeding season, and each species pair was assigned a score based on the number of overlapping axes (0 to 3).

### Quantification of territorial signal similarity

To determine whether interspecific territoriality could be predicted by overlap in common territorial signals, we quantified species similarity in territorial song and plumage coloration. To assess similarity in song, we downloaded high quality sound files from xeno-canto (https://www.xeno-canto.org/) and the Cornell Macaulay Library (Table S2) that matched the description in the Birds of North America of the vocalization used by each species for territorial advertisement and interactions. We categorized the size of the territorial repertoire for each species with descriptions in the Birds of North America, and determined the number of song files needed to capture repertoires of different sizes with a sensitivity analysis (Supplement 2, Figure S1). For species with relatively small repertoires (fewer than 4 song types), we collected 2 representative song files, and for species with relatively large repertoires (4 or more song types), we collected 4 song files. We performed noise reduction on sound files with background noise in Audacity version 2.1.3 (http://web.audacityteam.org/), using starting values of noise reduction = 12, sensitivity = 6, frequency smoothing = 0. We then normalized all sound files together.

To assess similarity in song between the species in a pair, we used two approaches. First, we calculated a measure of song dissimilarity based on numerous song parameters. We used the R package warbleR (Araya-Salas and Smith-Vidaurre 2016) to extract acoustic parameters (Table S3) and then additionally calculated the number of notes, length of the longest note, total note duration, average note duration, longest pause between notes, and average pause length per song. We averaged parameters for the sound files for each species and performed phylogenetic principal component analysis (pPCA; Revell 2009; Figure S2) on these averaged parameters (since pPCA requires exactly one data point per species in the phylogeny). We then calculated the Euclidean distance between all phylogenetic principal component scores for each species pair as a measure of song dissimilarity.

Second, we used spectral cross-correlation analysis (Clark et al. 1987) to quantify similarity in the frequency-time structure of song files. Spectral cross-correlation incrementally time-shifts spectrograms and calculates the cross-correlation between the frequency-time matrices of the spectrograms at each increment. We used the xcor function in warbleR to perform spectral cross-correlation analysis between all song files in a species pair, and averaged the maximum cross-correlation value from those comparisons as a second metric of song similarity. These two song measures are significantly correlated but not strongly enough to be considered redundant measures (*r* = -0.37, *N* = 45, *P* = 0.011).

To quantify similarity in plumage coloration and pattern, we recruited volunteers to score images of birds based on how similar they appeared. We obtained digital images of each species from two field guides (Sibley 2000; Dunn and Alderfer 2006) and asked participants to rank the plumage similarity of each species pair on a 0-4 scale using those images. We partitioned the images into seven surveys that we distributed with Survey Gizmo (https://www.surveygizmo.com) through social media and birding groups. Each survey contained approximately 30 pairs of images, with images repeated across surveys and within surveys, and a test for colorblindness. We filtered out incomplete responses and responses from participants who failed the color vision test. After obtaining at least 10 complete responses per survey, we calculated the mean similarity score for each species pair. Plumage similarity scores were strongly correlated between field guides (*ρ* = 0.79, *N* = 14), within surveys (*ρ* = 0.92, *N* = 14), and across surveys (*ρ* = 0.85, *N* = 14).

### Assessing ecological predictors of interspecific territoriality

We first used univariate tests to determine whether the trait differences (such as song similarity or bill length difference) within interspecifically territorial species pairs differed from non-interspecifically territorial species pairs. Because the potential to detect such differences depends on the level of variability among sister species, we calculated coefficients of variation for traits measured on a ratio scale and coefficients of nominal variation for binary traits (Kvålseth 1995).

To assess whether a single hypothesis explained the observed pattern of interspecific territoriality, we ran a generalized linear model with interspecific territoriality as a binomial response variable and the ecological, phenotypic, and behavioral traits in Table 1 as the predictor variables: hybridization (presence or absence), syntopy, ecomorphological differences, the number of overlapping foraging niche axes (0-3), song similarity (pPCA distance and maximum spectral cross-correlation), and plumage similarity. We also examined whether habitat complexity and species symmetries in dominance and aggression help explain the observed patterns (Supplement 3).

**Table 1.** Direction of association^†^ between predictor variables and interspecific territoriality, as predicted by four hypotheses.

|  | Misdirected aggression | Adaptive for resource competition | Adaptive for reproductive interference | Adaptive for asymmetric competition |
| --- | --- | --- | --- | --- |
| Patristic distance | – | – | – | – |
| Plumage similarity | + | + | + |  |
| Song similarity | + | + | + |  |
| Foraging guild overlap |  | + |  | + |
| Bill length difference |  | – |  | – |
| Mass difference |  | – |  | + |
| Hybridization | + |  | + |  |
| Syntopy | – | + | + | + |

<sup>†</sup>+, positive association; –, negative association

To evaluate whether interspecific territoriality has multiple origins, we included interactions between syntopy and other relevant predictor variables in the generalized linear model. Maladaptive interspecific territoriality, arising from misdirected aggression, should not persist between highly synoptic species that overlap extensively in breeding habitat and encounter each other frequently, whereas interspecific territoriality that is adaptive could persist between such species (Losin et al. 2016). To evaluate whether the misdirected aggression hypothesis and the reproductive interference hypothesis each explain a subset of the cases of interspecific territoriality, we included an interaction term between syntopy and hybridization. Under these two hypotheses, interspecific territoriality should primarily occur between non-hybridizing species with infrequent encounters or between hybridizing species that encounter each other frequently (Figure 1A). To test whether the misdirected aggression hypothesis and the resource competition hypothesis each explain a subset of the cases of interspecific territoriality, we included an interaction term between syntopy and the number of overlapping foraging guild axes. Under these two hypotheses, interspecific territoriality should primarily occur between species that encounter each other infrequently or between species with very similar ecological niches and breeding habitats (Figure 1B). Size asymmetry could be a proxy for exploitative resource competition (Losin et al. 2016), but also for whether one species is likely to dominate the other in aggressive interactions (Martin and Ghalambor 2014; Martin et al. 2017; Chock et al. 2018). Since sister species are on average very phenotypically similar, mass difference may not be a strong proxy for species differences in niche overlap, but even a small difference in size could impact aggressive interactions. Thus, we assume that size asymmetry is a better proxy for asymmetry in aggressive dominance than for resource competition in our dataset, and include an interaction term between syntopy and mass difference to test whether the misdirected aggression and asymmetric competition hypotheses each explain a subset of the cases of interspecific territoriality. Under these two hypotheses, interspecific territoriality should primarily occur between species that encounter each other infrequently or that occupy the same breeding habitats and are asymmetric in size (Figure 1C). For each of these linear models, we ran a second generalized linear model that included patristic distance as a predictor variable to control for phylogenetic non-independence.

**Figure 1.**
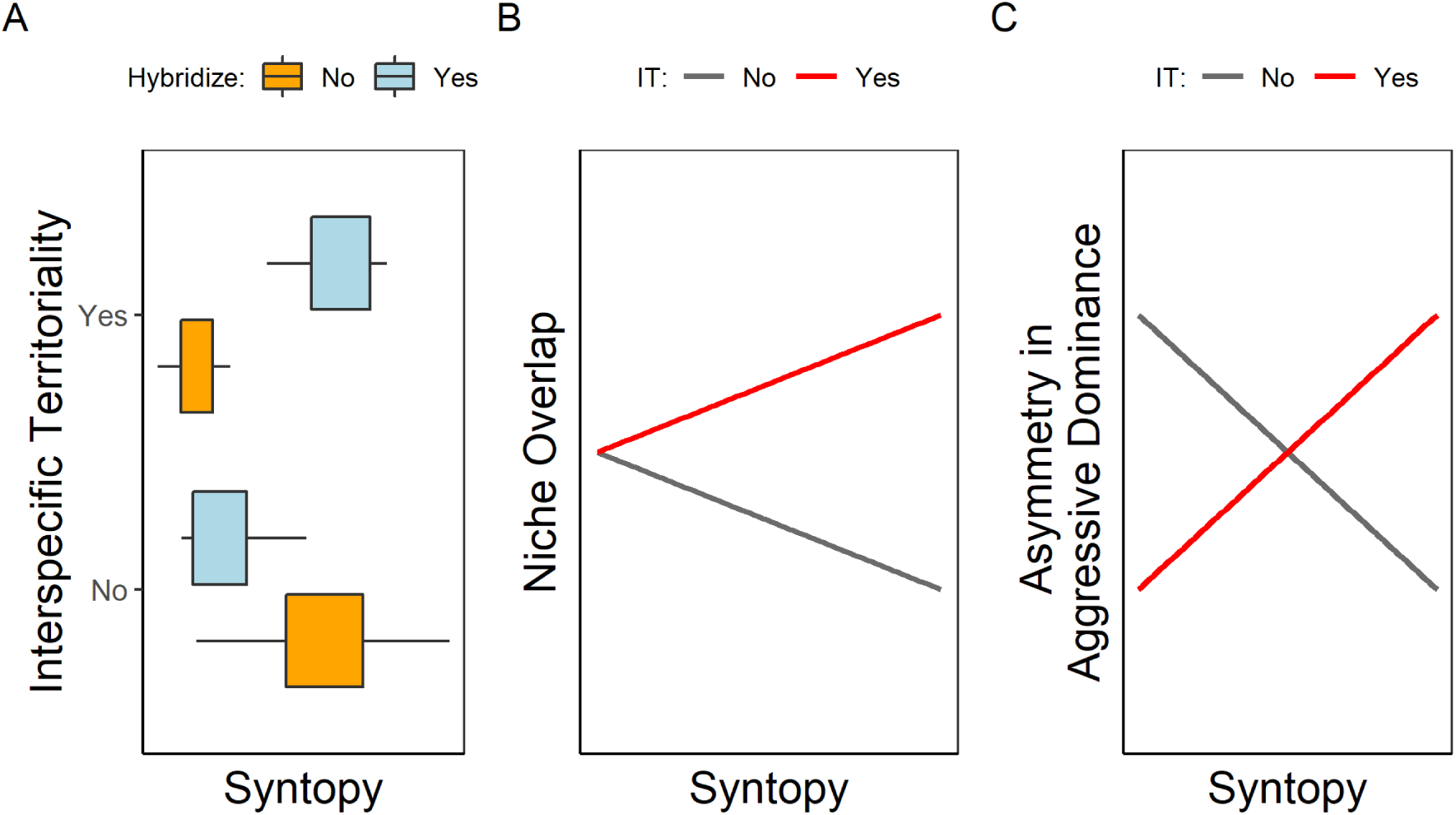
Predicted results if more than one hypothesis explains patterns of interspecific territoriality among closely related species. If the misdirected aggression and reproductive interference hypotheses each account for a subset of cases of interspecific territoriality (A), interspecific territoriality should primarily be found between hybridizing species that encounter each other frequently (high syntopy) or between species that rarely encounter each other (low syntopy). Under the misdirected aggression and the resource competition hypotheses (B), interspecific territoriality should primarily be found between species that encounter each other infrequently (low syntopy) or between species with very similar ecological niches and breeding habitats (high syntopy). The resource competition hypothesis further predicts that highly syntopic non-interspecifically territorial species occupy different ecological niches. Under the misdirected aggression and asymmetric competition hypotheses (C), interspecific territoriality occurs when species are low in syntopy or high in syntopy and one species dominates aggressive interactions.

While the syntopy metric captures variation among species pairs in fine-scale breeding habitat overlap in sympatry, the degree to which species are sympatric across their respective ranges might also affect whether interspecific territoriality persists in the zone of overlap. For example, gene flow from allopatry might swamp local adaptation in sympatry if the species are only sympatric in a small portion of their ranges. Thus, we examined whether controlling for breeding range sympatry impacted the results of each pair of phylogenetically controlled and non-phylogenetically controlled linear models examining ecological predictors of interspecific territoriality.

### Modeling transitions to sympatry

To test the hypothesis that behavioral interference shapes coarse-scale distributional patterns, we ran five generalized linear models with percent breeding range overlap as the response variable (using the R package betareg; Cribari-Neto and Zeileis 2010). In the first model, we used only patristic distance as a predictor to test whether breeding range overlap is related to divergence time. In subsequent models, we examined whether interspecific territoriality, hybridization, the combination of those two variables, or the interaction of those two variables predicted the percent breeding range overlap (Table S11). We compared these models with AICc.

Finally, to evaluate the effects of behavioral interference on regional coexistence with a more explicit evolutionary framework, we used two recent sister taxa approaches for modeling factors that impact the probability of species occurring in sympatry. These approaches assume allopatric speciation, which is thought to be the predominant mode of speciation in birds (Mayr 1942; Coyne and Orr 2004; Phillimore et al. 2008), and that following speciation, species transition from an allopatric phase to a parapatric phase before coming into broadly overlapping secondary sympatry (Cooney et al. 2017). First, we used a maximum likelihood approach to compare three types of models modified from Shi et al. (2018), in which the probability of occurring in sympatry depends on several parameters that describe how divergence time or other covariates relate to the probability of sympatry. The first model tests a null hypothesis that the probability of sympatry is based on the percent of species in sympatry and is unrelated to divergence time, while the two remaining models use different functions to associate divergence time, covariates, and the probability of sympatry (Supplement 4). Second, we implemented a multi-state Markov modeling approach (Pigot and Tobias 2013; Cooney et al. 2017) to assess whether interspecific territoriality impacts the rate at which species pairs transition from parapatry to sympatry. This approach assumes that the waiting time before transitioning to sympatry is associated with divergence time, but that there is a lag before sympatry is attained, which can represent species needing to diverge enough to be able to coexist in sympatry. We conducted simulations to determine whether the results we found were likely to occur by chance (Supplement 5). For both the multi-state Markov and the maximum likelihood approaches, we tested a range of values of continuous breeding range overlap (in 5% increments between 20% and 65%) as a cutoff between parapatric and sympatric distributions, as in Cooney et al. (2017). We did not consider the effect of interspecific territoriality or hybridization on transitions from allopatry to sympatry since it is not possible for allopatric species pairs to exhibit behavioral interference. For each approach, we compared models for which the rate or likelihood of transitioning between geographic states was determined only by phylogenetic distance to models that included interspecific territoriality, hybridization, or both as a covariate.

Finally, since the range of divergence times in a dataset can impact the generalization of how divergence time relates to sympatry from that dataset to other systems, we examined the range of phylogenetic distances in our dataset relative to other studies of sympatry in avian sister species (Supplement 6). To determine whether the species pairs in our dataset are older than average passerine sister species, we compared the phylogenetic distances between species pairs in our dataset to those of randomly sampled passerine sister species pairs (Supplement 6, Figure S4).

All data processing and statistical analyses were performed in R version 3.5.0.

## RESULTS

### Data Summary

In our dataset of true North American passerine sister species (*n* = 75), 63 (84%) pairs overlap in breeding range, and 35 (56%) of those are sympatric, defined as having at least 20% breeding range overlap. Only 12 sister species pairs are allopatric, and the remaining 28 are parapatric (< 20% breeding range overlap). After replacing allopatric sister species with the most closely related sympatric or parapatric species pairs, we were left with 71 phylogenetically independent pairs of closely related species. We were able to classify 48 of the 71 species pairs as interspecifically territorial or not. Excluding species that lacked information on patristic distance or breeding range overlap, our final dataset consisted of 45 sympatric or parapatric species pairs. Of those, approximately 21 pairs (47%) are interspecifically territorial.

In general, the species pairs in our dataset have similar plumage and song and overlap greatly in foraging guild, and also have low coefficients of variation for these variables (Table 2). The paired species vary most in morphological trait differences, syntopy, and sympatry (Table 2), and are relatively evenly divided across the categories of interspecifically territorial/non-interspecifically territorial and hybridizing/non-hybridizing (coefficient of nominal variation = 0.93 and 0.8, respectively). The average divergence time between species pairs is 4.7 Myr (range = 0.4 Myr – 34 Myr; Figure 2).

**Table 2.** Univariate comparisons between interspecifically territorial (I.T.) species pairs (N = 20) and non-interspecifically territorial (non-I.T.) species pairs (N = 25), and coefficients of variation.

| Variable | Transformation | Non-I.T. pairs |  | I.T. pairs |  | t | P<sub>t-test</sub> | CV |
| --- | --- | --- | --- | --- | --- | --- | --- | --- |
|  |  | Mean | SE | Mean | SE |  |  |  |
| Patristic distance | log | 2.10 | 0.04 | 1.44 | 0.04 | 2.35 | **0.012** | 117.86 |
| Song similarity (SPCC) |  | 0.34 | 0.01 | 0.40 | 0.01 | -1.61 | 0.058 | 35.68 |
| Song dissimilarity (pPCA) |  | 14.52 | 0.19 | 12.44 | 0.27 | 1.37 | 0.089 | 37.9 |
| Mass difference | log(x + 0.01) | 1.16 | 0.07 | 0.69 | 0.08 | 0.89 | 0.189 | 332.87 |
| Plumage dissimilarity |  | 1.76 | 0.04 | 1.78 | 0.04 | -0.09 | 0.535 | 49.85 |
| Syntopy | log(x + 0.01) | -3.59 | 0.02 | -3.57 | 0.04 | -0.09 | 0.536 | 95.49 |
|  |  | Median | Range | Median | Range |  | P<sub>Mann-Whitney</sub> | CV |
| Bill difference | log(x + 0.01) | 0.19 | -4.61 – 2.94 | -0.06 | -4.61 – 1.51 |  | 0.14 | 159.06 |
| Sympatry | sqrt | 0.79 | 0.14 – 0.98 | 0.52 | 0.05 – 1 |  | 0.17 | 61.93 |
| Foraging guild overlap |  | 3 | 1 – 3 | 3 | 0 – 3 |  | 0.58 | 28.88 |

**Figure 2.**
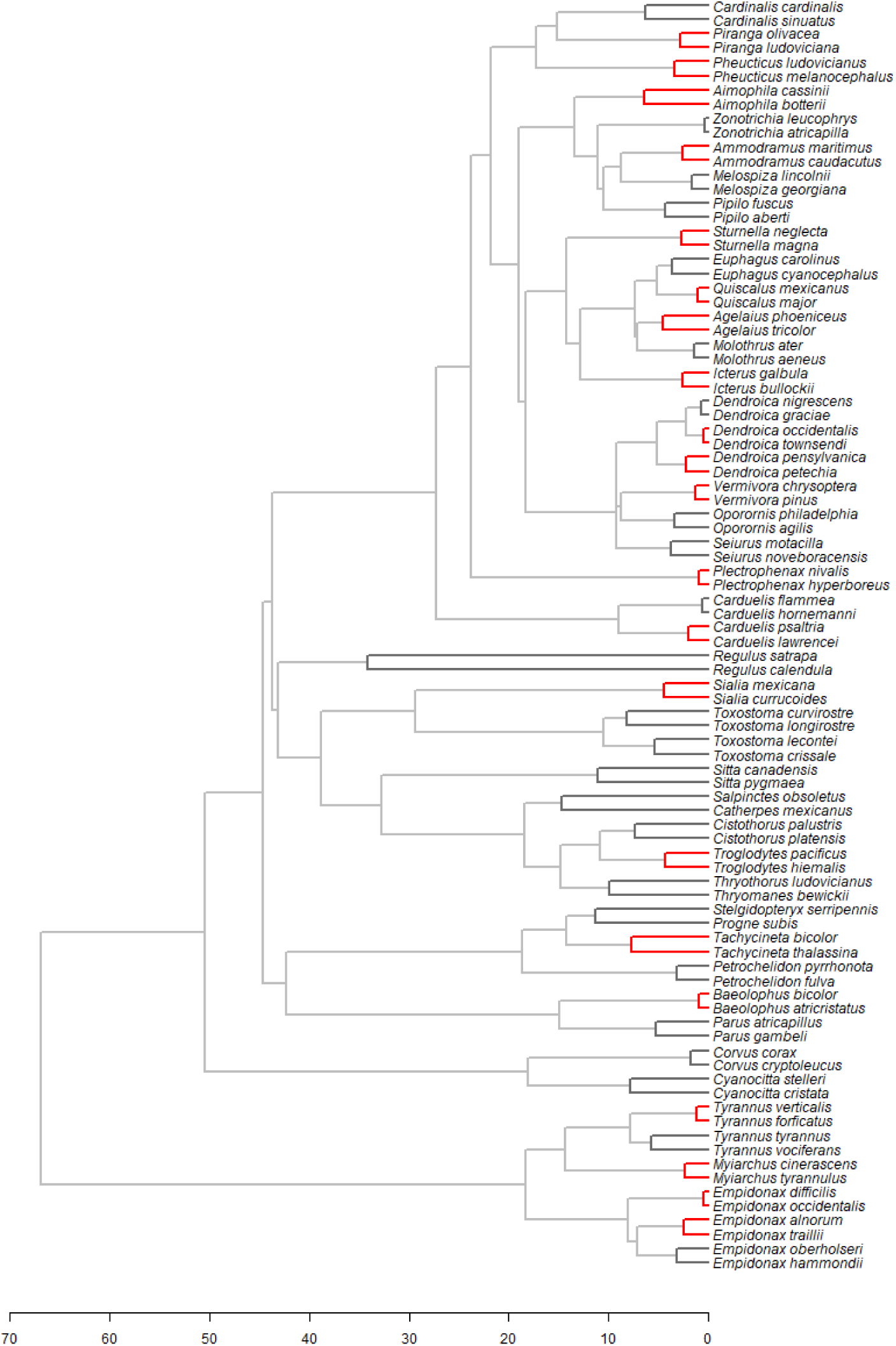
Interspecifically territorial sister species (red) are separated by shorter patristic distances (shaded branches; Myr), on average, than non-interspecifically territorial sister species (dark gray).

There were more records in the Zoological Records database for species pairs classified as interspecifically territorial than for species pairs classified as non-interspecifically territorial, suggesting that there could be unreported cases of interspecific territoriality (range_1_ = 0 – 53; range_2_ = 3 – 105; median_1_ = 7; median_2_ = 15; Mann-Whitney test, *n*_*1*_ = 24, *n*_*2*_ = 21, *P* = 0.015).

### Ecological predictors of interspecific territoriality

Interspecifically territorial species pairs are more closely related than non-interspecifically territorial species pairs (Table 2; Figure 2) but species pairs in these two categories do not differ significantly in other measured traits and behaviors (Table 2; 15 of 21 interspecifically territorial species pairs vs. 12 of 24 non-interspecifically territorial species pairs hybridize; Fisher’s exact test, *P* = 0.22).

The generalized linear models without interaction terms that we used to assess support for the four hypotheses separately (Table 1) yielded no significant predictors of interspecific territoriality (Tables S4, S5). However, in models with an interaction between hybridization and syntopy, the interaction term was significant: among hybridizing species, interspecifically territorial species are less syntopic than non-interspecifically territorial species, whereas among non-hybridizing species, interspecifically territorial species are more syntopic than non-interspecifically territorial species (Figure 3A, Table 3, S6). The results for hybridizing species are consistent with the misdirected aggression hypothesis but not with the reproductive interference hypothesis, while the results for the non-hybridizing species are consistent with the resource competition or the asymmetric competition hypotheses (Figure 1).

**Table 3.** Generalized linear model predicting interspecific territoriality with interaction between syntopy and hybridization.

| Variable | Estimate | SE | *z* | *P* |
| --- | --- | --- | --- | --- |
| (Intercept) | 1.19 | 3.04 | 0.39 | 0.696 |
| Syntopy | 5.88 | 2.64 | 2.23 | **0.026** |
| Hybridization | 3.63 | 1.78 | 2.04 | **0.042** |
| Plumage dissimilarity | 0.20 | 0.44 | 0.46 | 0.647 |
| Song dissimilarity (pPCA) | -0.70 | 0.58 | -1.20 | 0.230 |
| Song similarity (SPCC) | -0.88 | 0.66 | -1.33 | 0.184 |
| Mass difference | -0.16 | 0.40 | -0.40 | 0.691 |
| Bill length difference | -0.31 | 0.47 | -0.65 | 0.514 |
| Guild overlap | -1.69 | 1.05 | -1.61 | 0.108 |
| Syntopy x hybridization | -6.69 | 2.74 | -2.44 | **0.015** |

**Figure 3.**
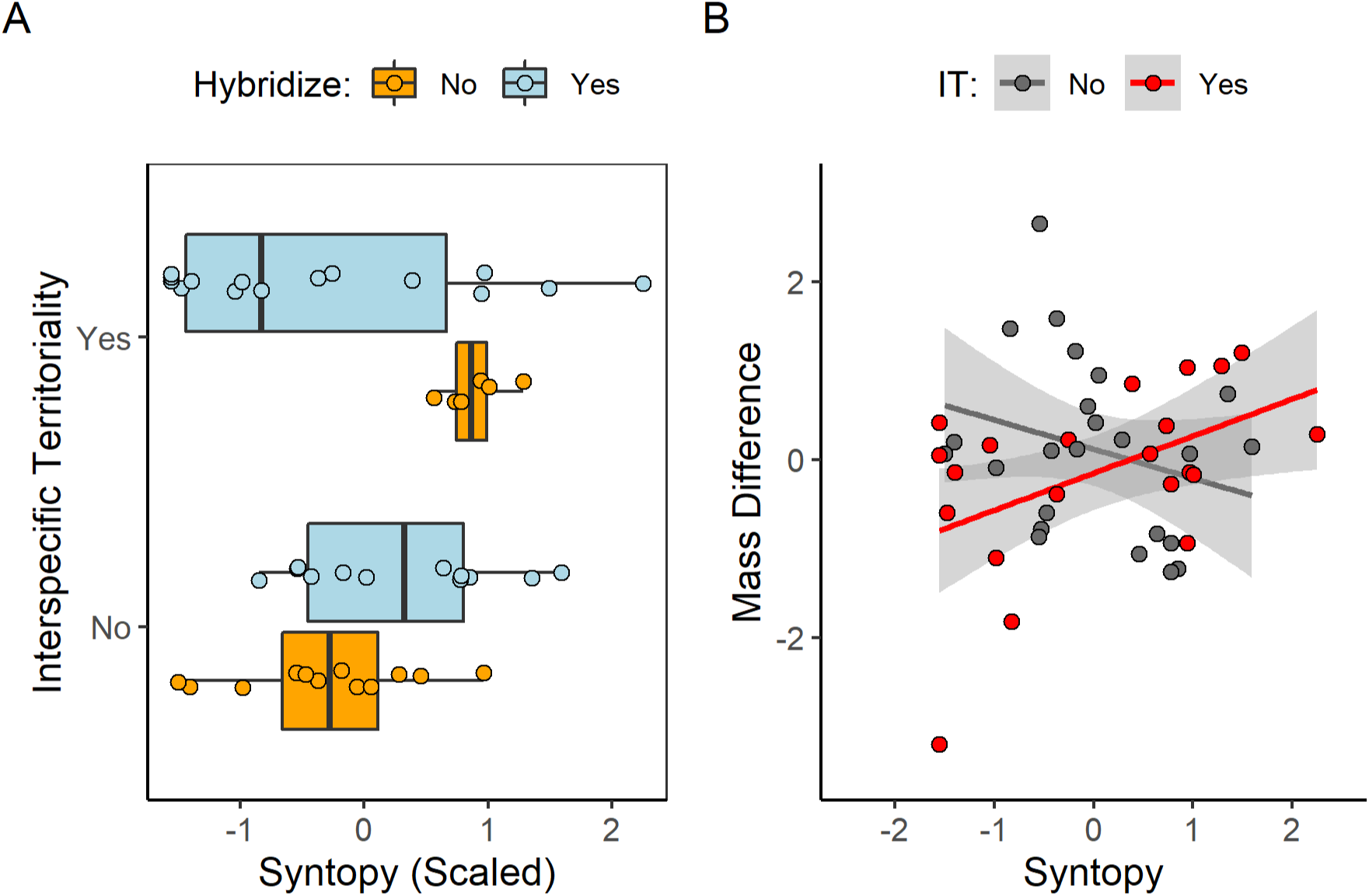
Interaction plots showing that (A) interspecifically territorial species that hybridize are less syntopic than non-interspecifically territorial species that hybridize, while interspecifically territorial species that do not hybridize are more syntopic than non-interspecifically territorial species that do not hybridize; (B) interspecifically territorial species (red) are more similar in size when low in syntopy than when high in syntopy, while the reverse is true for non-interspecifically territorial species (gray). Shading represents 95% confidence intervals. Mass difference and syntopy are both scaled to have a mean of zero and standard deviation of 1.

The models with an interaction between foraging guild overlap and syntopy yielded no significant terms (Tables S7, S8). In the models with an interaction between mass difference and syntopy, however, the interaction term emerged as positively associated with interspecific territoriality, regardless of phylogenetic correction, suggesting support for the misdirected aggression and the asymmetric competition hypotheses (Figure 3B, Tables 4, S9).

**Table 4.** Generalized linear model predicting interspecific territoriality with interaction between syntopy and size difference PC.

| Variable | Estimate | SE | *z* | *P* |
| --- | --- | --- | --- | --- |
| (Intercept) | 0.58 | 1.66 | 0.35 | 0.728 |
| Syntopy | -0.049 | 0.38 | -0.13 | 0.899 |
| Mass difference | -0.13 | 0.57 | -0.23 | 0.821 |
| Guild overlap | -0.51 | 0.62 | -0.82 | 0.414 |
| Hybridization | 0.79 | 0.86 | 0.93 | 0.353 |
| Plumage dissimilarity | 0.43 | 0.42 | 1.03 | 0.306 |
| Song dissimilarity (pPCA) | -0.45 | 0.43 | -1.03 | 0.302 |
| Song similarity (SPCC) | 0.13 | 0.49 | 0.26 | 0.791 |
| Bill length difference | 0.042 | 0.50 | 0.084 | 0.933 |
| Syntopy x mass difference | 1.78 | 0.75 | 2.37 | **0.018** |

Controlling for sympatry did not affect which terms were significant in any of the models, but in several cases the AICc score decreased (Table S10), i.e., sympatry improved the model fit.

### Transitions to sympatry

Regression models built to examine factors associated with breeding range sympatry suggest that the interaction of interspecific territoriality and hybridization may predict the degree of breeding range overlap, whereas the amount of time since divergence does not. Although the model with only patristic distance as an independent variable had the best AICc value, the effect size of patristic distance was small and its association with sympatry was non-significant (Table S11). The next best model (ΔAICc = 0.35) for predicting percent breeding range overlap included the interaction between interspecific territoriality and hybridization and did not include patristic distance (Table S11). In this model, the interaction between both forms of behavioral interference had a large effect size, although this was not statistically significant (*P* = 0.07; Table S11). Species that are both interspecifically territorial and hybridized appear to have narrower breeding range overlap relative to other species in the dataset (Figure 4).

**Figure 4.**
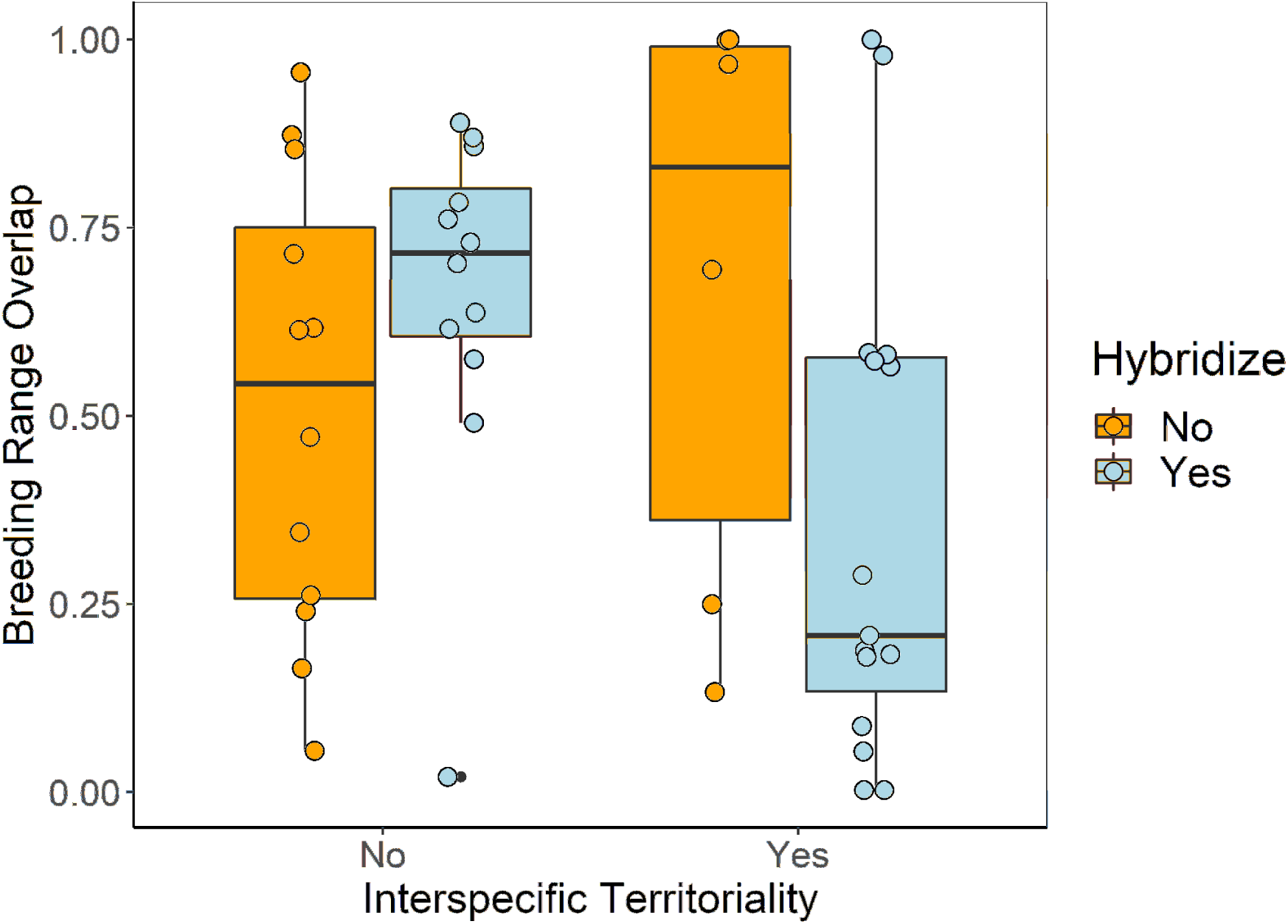
The best regression model for predicting percent breeding range overlap included the interaction between interspecific territoriality and hybridization (also see Table S11).

Further modeling of a categorical index of sympatry yielded similar results: the best model in the maximum likelihood approach for predicting sympatry includes the interaction between interspecific territoriality and hybridization and does not include patristic distance, regardless of the threshold of parapatry-sympatry considered (Tables S12-S18).

When explicitly modeling the transition rates in sympatry using the multi-state Markov models, results depended on the breeding range cutoff (Table S19). However, the confidence intervals around these waiting time estimates overlapped, indicating that none of the covariates significantly predicts the time it takes species to transition from parapatry to sympatry (Figures S3 and S4), and simulations on randomly shuffled data yielded similar results, suggesting that the observed results are likely to occur by chance (Supplement 5).

The species pairs in our true sister species dataset are not significantly older than random samples of passerine sister species pairs worldwide (Figure S5; Supplement 6).

## DISCUSSION

In the most phylogenetically diverse survey of interspecific territoriality completed so far, we found that interspecific territoriality occurs in almost half of all sympatric sister species of North American passerine birds. This finding alone suggests that interspecific interference competition ought to be an important consideration for researchers studying distributional patterns and diversification in birds. Whether interspecific territoriality is a maladaptive byproduct of intraspecific territoriality that reduces the prospects of species coexisting (Murray 1971) or instead is an evolved mechanism of spatial resource partitioning that stabilizes coexistence (Grether et al. 2013) is of obvious relevance for predicting its ecological and evolutionary effects.

Consistent with all four hypotheses (Table 1), we found that interspecifically territorial sister species are more closely related than non-interspecifically territorial sister species, despite the shallow timescale involved. Beyond that, however, none of the hypotheses’ specific predictions held up across the entire clade. As a whole, interspecifically territorial sister species are not less syntopic (i.e., do not overlap less in breeding habitat) than non-interspecifically territorial species, as the misdirected aggression hypothesis predicts, nor are they more syntopic, as the resource competition and reproductive interference hypotheses predict. Likewise, neither foraging guild overlap, morphological divergence, nor hybridization predict interspecific territoriality across the clade. In short, none of the foundational hypotheses alone accounts for the distribution of interspecific territoriality among sister species of North American perching birds.

To evaluate whether multiple hypotheses together could explain the distribution of interspecific territoriality, we included interactions between syntopy and other key predictor variables in the models. The logic behind this approach is that maladaptive interspecific territoriality should be eliminated quickly by selection if the species overlap broadly in breeding habitat, but it might persist indefinitely if the species rarely encounter each other (Losin et al. 2016). By contrast, adaptive forms of interspecific territoriality are more likely to evolve, and be maintained by selection, if the species are highly syntopic (Losin et al. 2016). Therefore, if both maladaptive and adaptive cases of interspecific territoriality occur in our dataset, we would expect to find significant interactions between syntopy and proxies for adaptive processes operating in these systems (Figure 1). We did indeed find such interactions (Figure 3).

Our results are consistent with the misdirected aggression and asymmetric competition hypotheses each explaining a subset of cases: we found that interspecifically territorial species that are low in syntopy are more similar in size, on average, than interspecifically territorial species that are high in syntopy (Figure 3B). Our findings from examining the interaction between syntopy and hybridization are also consistent with the misdirected aggression hypothesis and the asymmetric competition or resource competition hypotheses: the presence of hybridizing interspecifically territorial species that do not often encounter each other in breeding habitat may indicate that these species pairs engage in high levels of behavioral interference that might eventually be eliminated by agonistic character displacement (Grether et al. 2017), and the presence of non-hybridizing interspecifically territorial species that frequently co-occur in time and habitat suggests that interspecific territoriality may also arise as an adaptive response to resource competition among species that overlap broadly in breeding habitat. The finding that hybridizing species are more likely to be interspecifically territorial only when they are narrowly syntopic (Figure 3A) suggests that interspecific territoriality is not generally an adaptive response to reproductive interference among sister taxa. Instead, the combination of hybridization and interspecific territoriality in closely related species appears to be an unstable state that only persists when species have low encounter rates, but in the absence of hybridization, interspecific territoriality can mediate resource partitioning among highly syntopic species.

In combination, the misdirected aggression hypothesis and the resource competition hypothesis predict an interaction between foraging guild overlap and syntopy because the former hypothesis predicts that interspecific territoriality is associated with low syntopy while the latter predicts that interspecific territoriality is associated with high syntopy and high foraging guild overlap. We did not find such an association, but this might be due to low variation in the foraging guild metric; most species pairs in our dataset overlapped in all three foraging guild axes. While not all of the highly syntopic, interspecifically territorial species overlap in all three foraging axes, in theory even moderate levels of niche overlap can be sufficient to maintain interspecific territoriality (Grether et al. 2009).

Being larger in body size can provide an advantage in aggressive interactions between closely related species (Martin and Ghalambor 2014; Martin et al. 2017; Chock et al. 2018; Freeman 2019). Indeed, we found that, among highly syntopic species pairs in our dataset, those that are interspecifically territorial differ more in size than species that are not interspecifically territorial. Whether asymmetries in aggression explain this finding remains unresolved, however, because in many cases we were unable to determine whether one species was consistently the aggressor or victor (Supplement 3). Such asymmetries could be important for predicting evolutionary and ecological outcomes of interspecific interactions, just as asymmetries in exploitative competition or reproductive interference are recognized as critical for predicting outcomes of species coexistence (Tilman 1980; Amarasekare 2002; Kishi and Nakazawa 2013).

Even if size difference is unrelated to asymmetries in interspecific aggression among closely related North American passerines, size could still play an important role in the emergence of interspecific territoriality as an adaptive response to resource competition that permits coexistence between closely related species. For example, large differences in size could indicate asymmetric efficiency at exploiting a common limiting resource (Persson 1985), and interspecific territoriality could provide enough of an advantage to the less efficient resource exploiter for the two species to coexist (Grether et al. 2013). Alternatively, the increase in size difference between interspecifically territorial species across increasing levels of syntopy could represent divergence in morphology driven by ecological character displacement.

Interspecific territoriality can occur between species that identify heterospecific competitors via the same characters used to identify conspecific competitors, but may also occur between species that have evolved in competitor recognition and identify heterospecifics using a different character (Cody 1969, 1973; Grether et al. 2009). Although we could not directly measure competitor recognition for the species in our dataset, we tested whether characters commonly used by birds to identify conspecifics are associated with interspecific territoriality. Indeed, we found that song similarity likely plays a role in competitor recognition, since species that are interspecifically territorial are more similar in song than non-interspecifically territorial species, although this finding was marginally non-significant (Table 2).

Our study is similar in approach to a recent study of wood-warblers (Losin et al. 2016), but has distinct findings. Losin et al. (2016) inferred that interspecific territoriality is likely an adaptive response to competition in wood-warblers, but they were unable to determine whether hybridization or resource competition drives interspecific territoriality. In our study of closely related passerines, we found some evidence in support of the asymmetric competition hypothesis, but we also found that a subset of species pairs is best explained by the mistaken identity hypothesis. The most likely explanation for these differences is the average divergence time between species in the two datasets. Because wood-warbler species pairs on average have diverged less recently than the sister species in our dataset, interspecific territoriality in wood-warblers that may have at one point been the result of misdirected intraspecific aggression could have disappeared as species evolved mechanisms to discriminate between heterospecifics and conspecifics. Secondary contact between distantly related species is also unlikely to lead to mistaken species identity since plumage and song characteristics are more likely to be different with increased divergence time, so interspecific territoriality may never have developed as a maladaptive phenomenon for many of the wood-warbler species pairs.

Our work on sister species of North American perching birds also uncovered several noteworthy distributional patterns. Although several studies find that co-occurrence in secondary sympatry is associated with greater phylogenetic distance (Price 2010; Pigot and Tobias 2013), approximately 84% (71/85) of sister species in our dataset are sympatric, with an average breeding range overlap of 44.2% of the range of the species with the smaller range. We found that time since divergence does not predict whether species are in sympatry, which contrasts with patterns found in other avian groups (e.g., ovenbirds, Tobias et al. 2014; Old World warblers, Price 2010), but might be consistent with evidence that waiting times to sympatry are relatively short in temperate North America (Weir and Price 2011; Weir and Price 2019). Our results instead suggest that the combination of territoriality and hybridization between closely related species may limit their ability to coexist in extensive sympatry. The difference between our results and the findings of other studies is not because the species pairs in our dataset are significantly older (i.e., sharing a more distant common ancestor) than avian sister taxa tend to be; the species we included in these analyses are not significantly older than passerine sister species around the world and are similar in divergence time to species in several other studies (Supplement 6).

Taken together, our findings lend insight into the important role of behavioral interference in the early stages of secondary contact following allopatric speciation. Our results point to a possible stage in the speciation process of secondary contact between closely related species that treat each other as competitors and mates, thus remaining in parapatry until they diverge sufficiently in competitor and mate recognition. Other closely related species, however, have achieved breeding range sympatry and extensive fine-scale breeding range overlap along with, and perhaps in part because of, interspecific territorial aggression. We found that interspecific territoriality is common among closely related species of passerine birds, but that even at the tips of the songbird phylogeny, the ecological circumstances associated with interspecific territoriality are diverse. Our work suggests that the evolutionary stability of interspecific territoriality may also vary across taxa, and calls for additional empirical research to further improve our understanding of how interspecific territoriality arises and contributes to the ecologies and coexistence of animal species.

## Acknowledgments

We thank Alexa Sheldon, David Blake, Colette Troughton, Katherine Zhou, Prottasha Khan, Sierra Hovey, and Tarran Walter for assistance with data collection. For helpful comments on early drafts of this manuscript, we thank Robert Cooper, Shawn McEachin, Gaurav Kandlikar, Thomas Smith, Michael Alfaro, and Noa Pinter-Wollman. This research was funded by a grant to GFG from the National Science Foundation (DEB-1457844).

## Data Accessibility Statement

All data and code will be made available on Dryad upon initial acceptance.

