## Supporting Materials for "Multiple routes to interspecific territoriality in sister species of North American perching birds"

### Supplement 1: Syntopy calculations

Similarly to syntopy calculated from the BBS, syntopy from eBird was calculated within years, and then averaged across years. We excluded redundant records that occurred within 70 meters of each other and any records greater than 39.4 km from the record of the other species in the species pair, since 39.4 km is the length of a BBS route. We defined the number of “shared” locations as the number of records within 0.4 km of a record of the other species, corresponding to the 0.4 km search radius at BBS stops. We used predicted values from a quadratic zero-intercept linear regression of eBird syntopy on BBS syntopy ($R^{2}$ = 0.67, F_2,66_ = 68.99, p < 0.0001) to replace the missing BBS syntopy values. To evaluate whether the paired species are more (or less) syntopic than expected by chance, we calculated a second syntopy metric based just on the BBS data: we divided the number of stops where both species were found by the expected number of stops where both species would be found if their distributions were independent (as in Losin et al. (2016)). This metric equals zero if the species were never found at the same stop, one if the species were found together as often as expected by chance and greater than one if the species were found together more often than expected by chance.

### Supplement 2: Determining sample size for bird vocalizations based on repertoire size

Since songbird species vary in the size of their vocal repertoires, the number of song samples needed to capture the vocal array of a species could also vary depending on repertoire size. While there are many recordings for some species on online databases, other species have not been frequently recorded, and it can be difficult to find high quality song samples without background noise. Thus, we performed a sensitivity analysis to determine the number of song samples needed to represent species with small or large vocal repertoires. We randomly selected six species from each category of “small”, “medium”, and “large” repertoire size (1-4, 5-14, and 15 or more songs, respectively). For each species, we downloaded and edited six sound files from xeno-canto.org, following the methods in the main text. We chose six because it was difficult to find more than six high quality files without other birds or noises in the background for most species.

After collecting these songs, we calculated the “true” phylogenetic principal component scores for each species by averaging the acoustic parameters across the six song samples, and then performing phylogenetic principal component analysis (pPCA) with all 15 species. We then repeated this process of averaging acoustic parameters by species and calculating the pPCA score, but averaged parameters across different numbers of song samples depending on the size of the species’ repertoires. First, we used two song samples for each species regardless of repertoire size. Next, we used two samples for species with small repertoires or with medium-sized repertoires, and four samples for species with large repertoires. Next, we used two samples for species with small repertoires and four samples for species with medium or large repertoires. Finally, we averaged across two samples for species with small repertoires, four for species with medium repertoires, and six for species with large repertoires. To compare the pPCA scores calculated from these sampling schemes, we repeated the process of randomly sampling songs from the six files we downloaded, averaging acoustic parameters for those songs, and calculating the pPCA scores for each species one hundred times, and took the correlation between these pPCA scores with the “true” scores. The sampling scheme that yielded high correlations with the fewest samples needed was sampling two songs for species with small repertoires and four samples for species with medium or large repertoires (Figure S1). Thus, for species with a repertoire size of one to four songs, we collected two songs, and for species with a repertoire of five or more songs, we collected four songs.

### Supplement 3: Testing hypotheses about the effects of habitat complexity and asymmetries in dominance and aggression

Orians and Willson (1964) proposed that interspecific territoriality is maladaptive but could persist between two species in the presence of ecological constraints, such as habitat that is to “simple” to support multiple niches, or adjacent niches are already occupied by other species, because these constraints prevent the species from diverging. Although MacArthur (1972) did not describe a mechanism for how a species could dominate territorial interactions, asymmetries in body size often lead to asymmetries in dominance (Martin and Ghalambor 2014; Martin et al. 2017; Chock et al. 2018). Here, we test predictions from these two hypotheses.

We categorized the habitat complexity of each species using descriptions from the Birds of North America Online and categories from Losin et al. (2016), with 1 representing a simple habitat such as a grassland, marsh, or tundra, 2 representing intermediate habitat, such as chaparral or forest edge, and 3 representing complex habitat such as deciduous forest. If a study in our literature search indicated whether one species initiated the interspecific territorial defense or successfully displaced the other species, we recorded that species as the “aggressor” or the “dominant” species, respectively.

To test the hypothesis that interspecific territoriality persists among ecological competitors living in simple habitats, we performed a Fisher’s exact test to evaluate whether interspecific territoriality is associated with either species occurring in a simple habitat. To test the hypothesis that interspecific territoriality occurs because it benefits the dominant species, and that the larger species are dominant, we performed a binomial test of whether size difference corresponds to being the “aggressor” or “dominant” species in the interspecific territorial encounters observed, and compared our findings to the regression model that included an interaction between syntopy and the size principal component.

*Results*

In contrast to the Orians and Willson (1964) prediction that interspecific territoriality could persist in simple habitats, habitat complexity did not differ between interspecifically territorial and non-interspecifically territorial species pairs (Fisher’s exact test, *P* = 0.15), with most interspecifically territorial species pairs (32 out of 42 pairs we could score) occurring in complex or intermediate habitats. For the 15 species pairs we were able to categorize (of 21 interspecifically territorial species pairs) according to aggression and dominance in interspecific territorial interactions, aggression was asymmetric in 8 pairs and dominance was asymmetric in 10 pairs. For pairs in which there is asymmetry in territorial interactions, the dominant species is not significantly larger (binomial test, *N* = 10, *P* = 0.95), nor is the aggressive species consistently larger (binomial test, *N* = 8, *P* = 0.94). In conclusion, we do not find support for the Orians and Willson (1964) hypothesis (neither did Losin et al (2016)), and we do not find evidence that asymmetries in aggression and dominance are related to asymmetries in size. However, we caution that many of our categorizations of aggression and dominance came from only two observations of species interactions, whereas other studies of asymmetrical aggression have required at least six observations (Martin and Ghalambor 2014).

### Supplement 4: Explanation of the models in the maximum likelihood approach

We adapted a maximum likelihood approach from Shi et al. (2018) to modelling the likelihood of species pairs transitioning from parapatry to sympatry. This approach fits three models in which the probability of occurring in sympatry varies according to a single parameter. In the first model, this parameter is a constant. In the second model, this parameter varies as an exponential decay function such that sympatry varies with the time since speciation, either by itself or in combination with a behavioral interference covariate. In the third model, this parameter varies logistically, again with phylogenetic distance alone or in combination with a behavioral interference covariate. For covariates, we considered interspecific territoriality alone, hybridization alone, or the combination of interspecific territoriality and hybridization.

### Supplement 5: Assessing false positive rate in multistate Markov modeling approach

To assess whether the multistate Markov modeling approach we used is likely to yield false positives with a dataset of the size and structure as ours, we performed simulations as follows. First, we picked two results from our actual data to test: (1) we calculated the ΔAICc value between a model with hybridization as a covariate and a parapatry-sympatry cutoff = 35% compared to a model with the same cutoff but with no covariates (since the former model had the lowest AICc value at a cutoff of 35% range overlap). (2) We calculated the ΔAICc for a model with interspecific territoriality x hybridization (I.T.*hybrid) as a covariate and a parapatry-sympatry cutoff value of 65%, compared to a model with the same cutoff and no covariates (again, since the former model had the lowest AICc value of all models tested at a cutoff of 65% range overlap).

(so in this case, a smaller delta AIC value means that the raw AIC score of the covariate model is smaller than the AIC score of the no-covariate model)

Second, we simulated 500 datasets by shuffling the hybridization assignments 500 times, and ran a Markov model for each of these with hybridization as the covariate and a cutoff value of 35% range overlap. We then simulated another 500 datasets by shuffling I.T.*hybrid 500 times, and ran a Markov model for each of these with I.T.*hybrid as the covariate and a cutoff value of 65% range overlap. We then calculated ΔAICc values for these simulated datasets, compared to the models we previously ran with no covariates. We then compared the ΔAICc value calculated from the actual data to this simulated distribution of ΔAICc values. Since we subtracted AICc_covariate model_ – AICc_no covariate_, the smaller the ΔAICc means the smaller the AICc_covariate model_ relative to the AICc_no covariate_. Thus, we looked to see whether the actual ΔAICc was in the lower 5% of the simulated distribution of ΔAICc values.

Third, we repeated this simulation approach, but instead of using randomly shuffled data, we simulated hybridization and interspecific territoriality to determine whether random factors that are associated with phylogenetic distance would come out as important in the model (in other words, does the signal of phylogenetic relatedness drive the result?). To simulate hybridization, we simulated two traits with EvoRAG and found the difference between the traits for each species pair, and assigned hybridization to the most similar species pairs (so that the number of hybrids matches the number in the actual dataset). For I.T.*hybrid, we simulated two additional traits (to represent interspecific territoriality), calculated the distance between all four simulated traits, and then assigned I.T.*hybrid = 1 to the most similar pairs, such that this number matched the number in the actual dataset.

*Results*

For each of these four comparisons (two models (hybrid and IT*hybrid) x two types of simulations), I found that the observed ΔAICc value was not smaller than the lower 5% of the distribution. This finding indicates that the results I found in the Markov models of our dataset are not exceptional relative to either randomly shuffled data or data where the covariates are associated with phylogenetic distance.

### Supplement 6: Comparison of average age of North American sister passerines to samples of worldwide sister passerines

Our dataset consists of species pairs that are “sister” species or the next most closely related species according to a North American passerine phylogeny, and in some cases, the “true” sister species occurs outside of North America. To evaluate whether the average divergence time between the species pairs in our study is significantly greater than in other studies of sister species that do not include continental boundaries, we conducted a simulation. Using a phylogeny of all bird species and a posterior distribution with 10,000 trees from birdtree.org (Jetz et al. 2012), we extracted all passerine species and identified all sister species pairs that had 90% node support across the posterior distribution. From that list, we randomly sampled 55 species pairs, corresponding to the number of species in the dataset we used to compare patterns of patristic distance and range overlap, averaged the patristic distances in that sample, and repeated this process 1000 times.

The average phylogenetic divergence time in our dataset of 55 species is 4.42 Myr, which is less than the upper 95% quartile from the simulations, although close to the upper quartile. Thus, we conclude that our dataset contains species pairs that are comparable in age to worldwide passerine sister species, although on the older end of the distribution (Figure S5).

Several studies of sister species of birds have found that sympatric species pairs are greater in phylogenetic distance than allopatric species. Since this contrasts with our findings (see Results), we examined the divergence times of two of these studies to consider the age of the species in those datasets relative to the species in our dataset. The sympatric sister species we studied diverged on average 4.42 Myr, with divergence times ranging from 0.4 Myr to 34 Myr; excluding the ruby-crowned and golden-crowned kinglets, the divergence times range from 0.4 Myr to 14.7 Myr. A study of sympatric sister lineages of ovenbirds (Furnariidae; Pigot and Tobias 2013) found that secondary sympatry is associated with greater phylogenetic and morphological distance. The sister species included in their dataset are have diverged on average 3.56 Myr ago (range: 0.12 – 11.68 Myr), which is younger than but still comparable to our dataset. A study of species of two genera of Eurasian Old World leaf warblers (Price 2010) considered species ranging from about 1 Myr old to about 12 Myr ago.

**Figures (S1-S4) and Tables (S1-S19)**

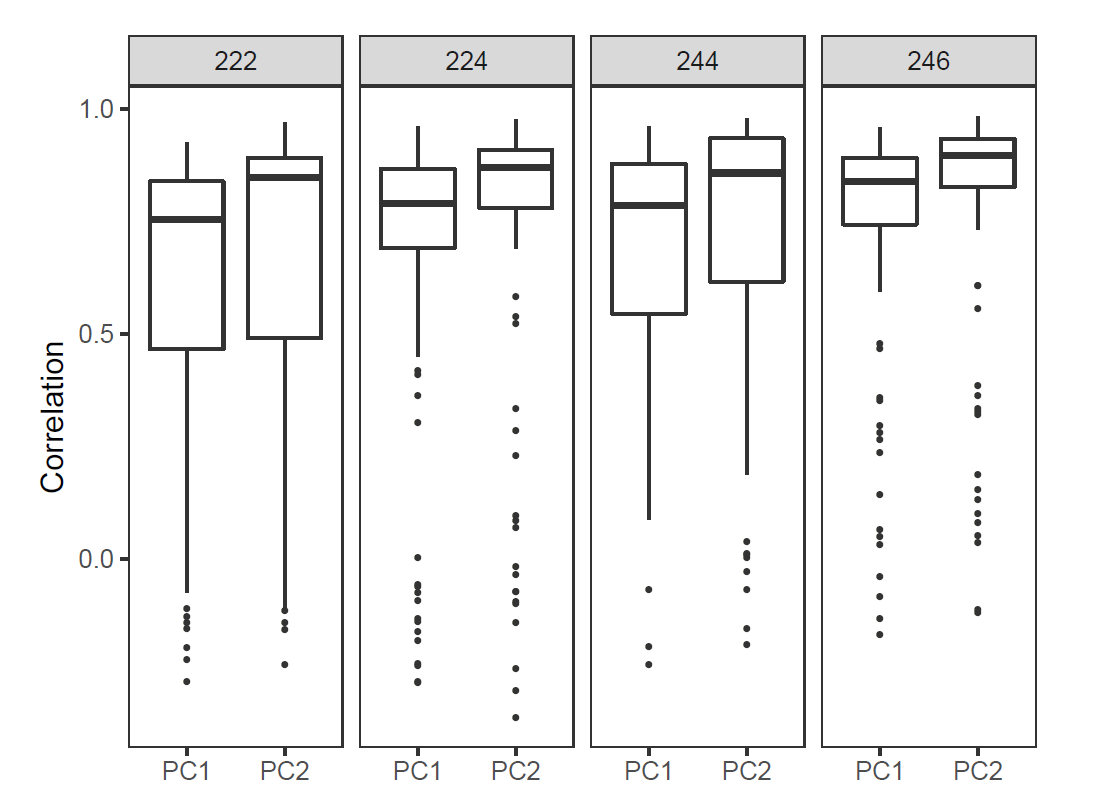

### Figure S1. Song simulation results depicting the correlation between the scores calculated from each sampling scheme and the scores calculated across six song files for each species. Each panel represents a different sampling scheme based on the size of species repertoires. For example, 224 means two songs sampled from species with small repertoires, two songs sampled from species with medium repertoires, and four songs sampled from species with large repertoires.

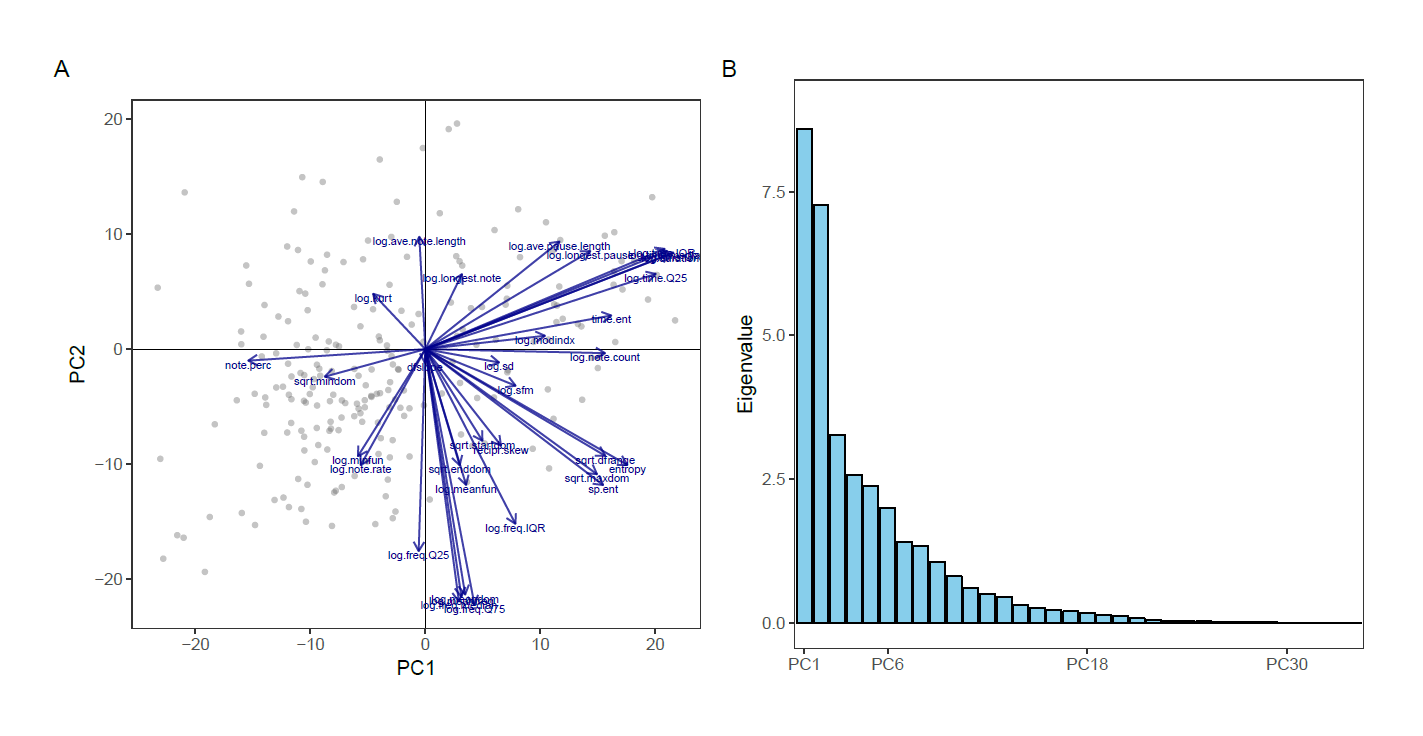

### Figure S2. Phylogenetic principal component analysis of acoustic parameters with (A) variable loadings and (B) variation captured by each principal component axis.

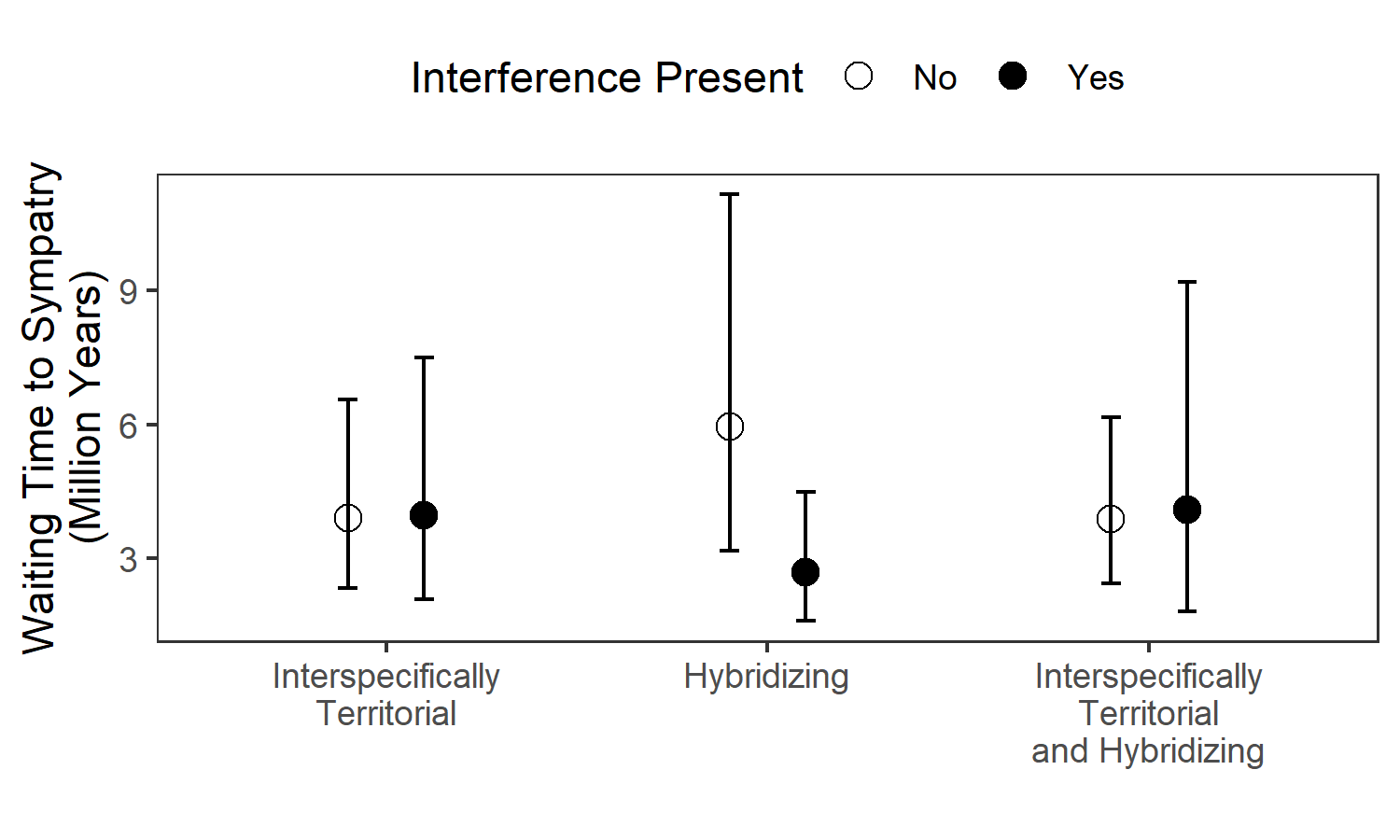

### Figure S3. Rates of transitioning into sympatry based on multi-state Markov modeling approach, with sympatry defined as more than 35% breeding range overlap (these results are identical to results using a cutoff of 40% and 45%, and similar to results using a cutoff of 30%, 50%, and 55%). Each multi-state Markov model includes all species pairs in the dataset, and varies in the kind of behavioral interference included as a covariate (interspecific territoriality, hybridization, or the interaction of the two). Estimated waiting times for species that do not engage in the behavioral interference (empty circles) or that do engage in the behavioral interference are calculated from maximum likelihood parameters, with error bars represent 95% confidence intervals around the estimates.

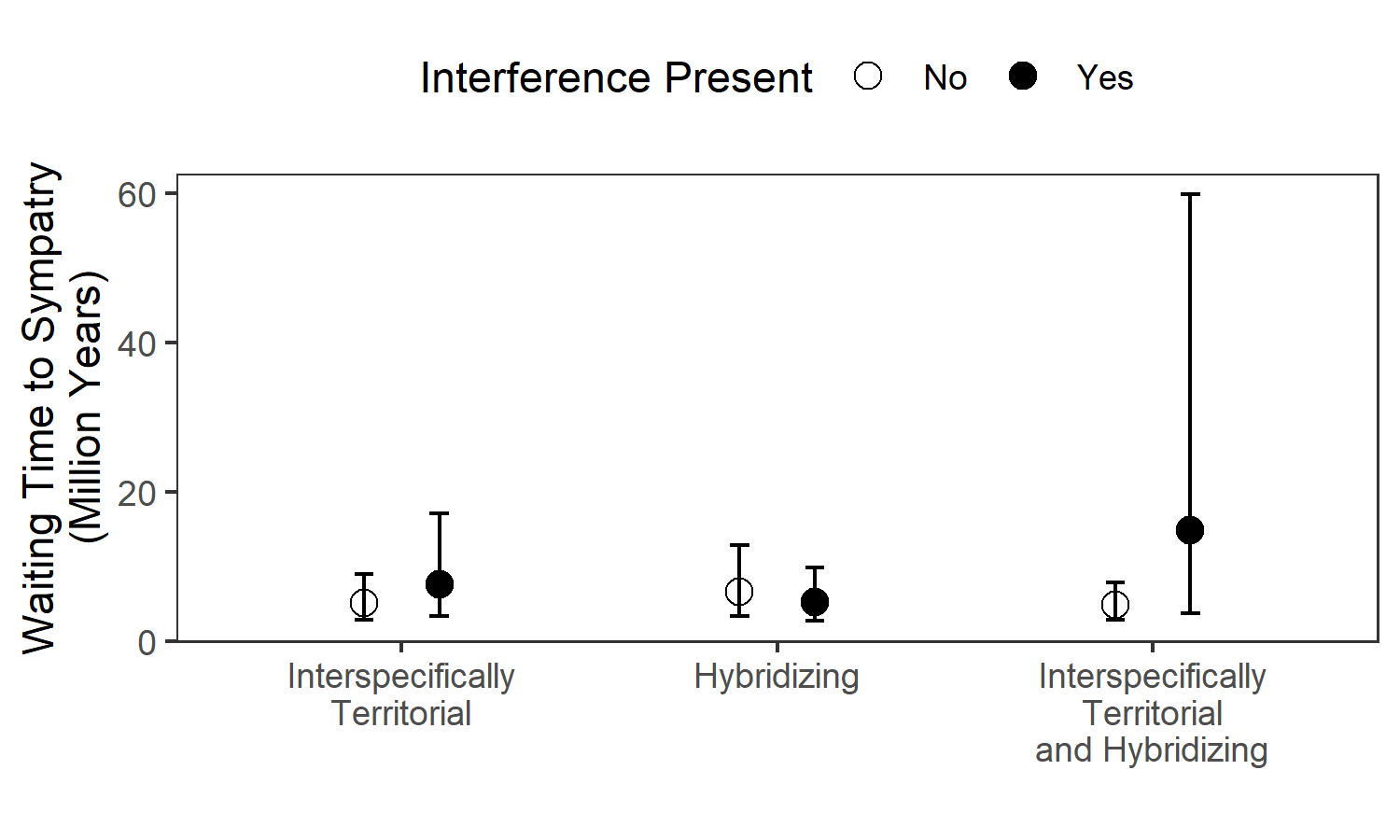

### Figure S4. Rates of transitioning into sympatry based on multi-state Markov modeling approach, with sympatry defined as more than 60% breeding range overlap. Each multi-state Markov model includes all species pairs in the dataset, and varies in the kind of behavioral interference included as a covariate (interspecific territoriality, hybridization, or the interaction of the two). Estimated waiting times for species that do not engage in the behavioral interference (empty circles) or that do engage in the behavioral interference are calculated from maximum likelihood parameters, with error bars represent 95% confidence intervals around the estimates.

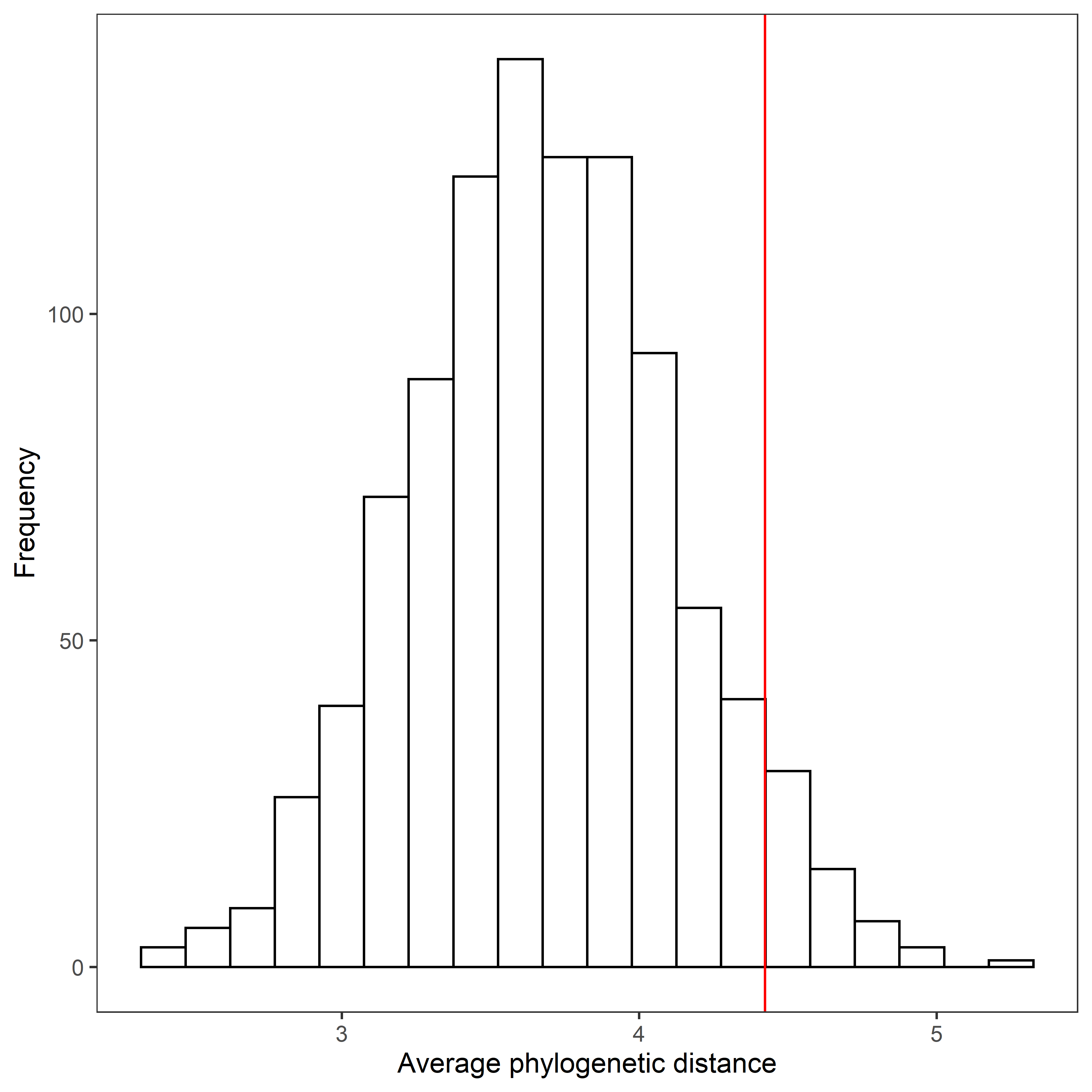

### Figure S5. Comparison of average divergence time (million years ago) in North American species pair dataset (red line, N = 55) to average divergence times of passerine sister species worldwide (from 1000 samples of N = 55 species pairs).

### Table S1. Interspecific territoriality (I.T.) classifications based on literature search

| Species 1 | Species 2 | I.T.? | References |
| --- | --- | --- | --- |
| *Aphelocoma insularis* | *Aphelocoma californica* | Allopatric |  |
| *Lanius excubitor* | *Lanius ludovicianus* | Allopatric |  |
| *Baeolophus inornatus* | *Baeolophus ridgwayi* | Allopatric |  |
| *Sitta pygmaea* | *Sitta pusilla* | Allopatric |  |
| *Toxostoma rufum* | *Toxostoma longirostre* | Allopatric |  |
| *Calcarius ornatus* | *Calcarius pictus* | Allopatric |  |
| *Setophaga virens* | *Setophaga chrysoparia* | Allopatric |  |
| *Cardellina pusilla* | *Cardellina rubrifrons* | Allopatric |  |
| *Melozone aberti* | *Melozone crissalis* | Allopatric |  |
| *Ammodramus henslowii* | *Ammodramus bairdii* | Allopatric |  |
| *Junco phaeonotus* | *Junco hyemalis* | Allopatric |  |
| *Peucaea aestivalis* | *Peucaea cassinii* | Allopatric |  |
| *Empidonax traillii* | *Empidonax alnorum* | Yes | Prescott 1987;Hechtenthal 2007; Gorski 1969 |
| *Tyrannus forficatus* | *Tyrannus verticalis* | Yes | Sutton 1967 |
| *Baeolophus atricristatus* | *Baeolophus bicolor* | Yes | Curry & Patten 2016 |
| *Tachycineta thalassina* | *Tachycineta bicolor* | Yes | Winkler 2011 |
| *Polioptila californica* | *Polioptila melanura* | Yes | Atwood 1988 |
| *Sialia currucoides* | *Sialia mexicana* | Yes | Herlugson 1980; Duckworth 2013; Pinkowski 1979; Duckworth and Badyaev 2007 |
| *Spinus lawrencei* | *Spinus psaltria* | Yes | Coutlee 1968; Linsdale 1957 |
| *Plectrophenax hyperboreus* | *Plectrophenax nivalis* | Yes | Winker et al 2002; Maley & Winker 2010; Johnson et al. 2013; Sealy 1969 |
| *Vermivora cyanoptera* | *Vermivora chrysoptera* | Yes | Ficken and Ficken 1968; Confer and Larkin 1998; Confer and Knapp 1977; Will 1986; Confer et al. 2003 |
| *Setophaga petechia* | *Setophaga pensylvanica* | Yes | Morse 1966 |
| *Setophaga townsendi* | *Setophaga occidentalis* | Yes | Pearson 2000, Pearson and Manuwal 2000, Pearson and Rohwer 2000 |
| *Icterus bullockii* | *Icterus galbula* | Yes | Edinger 1985 |
| *Agelaius tricolor* | *Agelaius phoeniceus* | Yes | Payne 1969 |
| *Quiscalus major* | *Quiscalus mexicanus* | Yes | Selander & Giller 1961; Pratt 1973 |
| *Sturnella magna* | *Sturnella neglecta* | Yes | Lanyon 1957; Wiens 1969; Lanyon 1956b; Elbert 2010; Rohwer 1973 |
| *Pheucticus melanocephalus* | *Pheucticus ludovicianus* | Yes | Kroodsma 1974 |
| *Piranga ludoviciana* | *Piranga olivacea* | Yes | Peterson 1995 |
| *Aphelocoma woodhouseii* | *Aphelocoma californica* | Yes | Correspondance with John McCormack |
| *Peucaea botterii* | *Peucaea cassinii* | Yes | Tramontano 1971 |
| *Myiarchus tyrannulus* | *Myiarchus cinerascens* | Yes | Brush 1983 |
| *Ammodramus caudacutus* | *Ammodramus maritimus* | Yes | Woolfenden 1956; BNA |
| *Empidonax occidentalis* | *Empidonax difficilis* | Yes (recent split) | Johnson 1980; Correspondance with Peter Lowther |
| *Troglodytes pacificus* | *Troglodytes hiemalis* | Yes (recent split) | Toews and Irwin 2008 |
| *Empidonax hammondii* | *Empidonax oberholseri* | No |  |
| *Tyrannus vociferans* | *Tyrannus tyrannus* | No |  |
| *Cyanocitta cristata* | *Cyanocitta stelleri* | No |  |
| *Corvus cryptoleucus* | *Corvus corax* | No |  |
| *Poecile gambeli* | *Poecile atricapillus* | No |  |
| *Petrochelidon fulva* | *Petrochelidon pyrrhonota* | No |  |
| *Progne subis* | *Stelgidopteryx serripennis* | No |  |
| *Thryomanes bewickii* | *Thryothorus ludovicianus* | No |  |
| *Cistothorus platensis* | *Cistothorus palustris* | No |  |
| *Catherpes mexicanus* | *Salpinctes obsoletus* | No |  |
| *Toxostoma crissale* | *Toxostoma lecontei* | No |  |
| *Bombycilla cedrorum* | *Bombycilla garrulus* | No |  |
| *Regulus calendula* | *Regulus satrapa* | No |  |
| *Acanthis hornemanni* | *Acanthis flammea* | No |  |
| *Parkesia noveboracensis* | *Parkesia motacilla* | No |  |
| *Setophaga graciae* | *Setophaga nigrescens* | No |  |
| *Euphagus cyanocephalus* | *Euphagus carolinus* | No |  |
| *Melospiza georgiana* | *Melospiza lincolnii* | No |  |
| *Ammodramus caudacutus* | *Ammodramus nelsoni* | No |  |
| *Zonotrichia atricapilla* | *Zonotrichia leucophrys* | No |  |
| *Cardinalis sinuatus* | *Cardinalis cardinalis* | No |  |
| *Sitta pygmaea* | *Sitta canadensis* | No |  |
| *Toxostoma curvirostre* | *Toxostoma longirostre* | No |  |
| *Geothlypis philadelphia* | *Oporornis agilis* | No |  |
| *Molothrus aeneus* | *Molothrus ater* | No |  |
| *Melozone aberti* | *Melozone fusca* | No |  |
| *Piranga ludoviciana* | *Piranga flava* | No |  |
| *Sayornis phoebe* | *Sayornis nigricans* | No data |  |
| *Contopus virens* | *Contopus sordidulus* | No data |  |
| *Myiarchus crinitus* | *Myiarchus cinerascens* | No data |  |
| *Pica nuttalli* | *Pica hudsonia* | No data |  |
| *Corvus caurinus* | *Corvus brachyrhynchos* | No data |  |
| *Vireo plumbeus* | *Vireo cassinii* | No data |  |
| *Vireo gilvus* | *Vireo philadelphicus* | No data |  |
| *Oreoscoptes montanus* | *Mimus polyglottos* | No data |  |
| *Catharus fuscescens* | *Catharus minimus* | No data |  |
| *Haemorhous purpureus* | *Haemorhous cassinii* | No data |  |
| *Loxia leucoptera* | *Loxia curvirostra* | No data |  |
| *Geothlypis philadelphia* | *Geothlypis tolmiei* | No data |  |
| *Oreothlypis luciae* | *Oreothlypis virginiae* | No data |  |
| *Setophaga striata* | *Setophaga castanea* | No data |  |
| *Icterus parisorum* | *Icterus graduacauda* | No data |  |
| *Icterus spurius* | *Icterus cucullatus* | No data |  |
| *Dolichonyx oryzivorus* | *Xanthocephalus xanthocephalus* | No data |  |
| *Chondestes grammacus* | *Calamospiza melanocorys* | No data |  |
| *Pipilo erythrophthalmus* | *Pipilo maculatus* | No data |  |
| *Spizella breweri* | *Spizella pusilla* | No data |  |
| *Spizella pallida* | *Spizella passerina* | No data |  |
| *Ammodramus savannarum* | *Arremonops rufivirgatus* | No data |  |
| *Passerina caerulea* | *Passerina amoena* | No data |  |
| *Passerina ciris* | *Passerina versicolor* | No data |  |
| *Artemisiospiza nevadensis* | *Artemisiospiza belli* | No data |  |
| *Catharus fuscescens* | *Catharus bicknelli* | No data |  |
| *Cardellina pusilla* | *Cardellina canadensis* | No data |  |

### Table S2. Sound files downloaded from xeno-canto to characterize song similarity between species pairs.

| Recording ID | Species | Recordist | URL |
| --- | --- | --- | --- |
| 316236 | *Sayornis phoebe* | Antonio Xeira | <https://www.xeno-canto.org/316236> |
| 132866 | *Sayornis phoebe* | Jonathon Jongsma | <https://www.xeno-canto.org/132866> |
| 318152 | *Sayornis nigricans* | Bobby Wilcox | <https://www.xeno-canto.org/318152> |
| 192057 | *Sayornis nigricans* | Paul Marvin | <https://www.xeno-canto.org/192057> |
| 110655 | *Sayornis nigricans* | Lauren Harter | <https://www.xeno-canto.org/110655> |
| 237806 | *Sayornis nigricans* | Jarrod Swackhamer | <https://www.xeno-canto.org/237806> |
| 72079 | *Sayornis nigricans* | Richard E Webster | <https://www.xeno-canto.org/72079> |
| 313072 | *Sayornis nigricans* | Ted Floyd | <https://www.xeno-canto.org/313072> |
| 18694 | *Contopus virens* | David Bradley | <https://www.xeno-canto.org/18694> |
| 158909 | *Contopus virens* | Ian Cruickshank | <https://www.xeno-canto.org/158909> |
| 141880 | *Contopus sordidulus* | Patrick Turgeon | <https://www.xeno-canto.org/141880> |
| 190093 | *Contopus sordidulus* | Richard E. Webster | <https://www.xeno-canto.org/190093> |
| 161658 | *Empidonax hammondii* | Ian Cruickshank | <https://www.xeno-canto.org/161658> |
| 196460 | *Empidonax hammondii* | Richard E. Webster | <https://www.xeno-canto.org/196460> |
| 14293 | *Empidonax oberholseri* | Andrew Spencer | <https://www.xeno-canto.org/14293> |
| 161661 | *Empidonax oberholseri* | Ian Cruickshank | <https://www.xeno-canto.org/161661> |
| 196032 | *Empidonax traillii* | Richard E. Webster | <https://www.xeno-canto.org/196032> |
| 6757 | *Empidonax traillii* | Darrell L. Peterson | <https://www.xeno-canto.org/6757> |
| 302253 | *Empidonax alnorum* | James Bradley | <https://www.xeno-canto.org/302253> |
| 376468 | *Empidonax alnorum* | Michael Harrison | <https://www.xeno-canto.org/376468> |
| 254619 | *Empidonax occidentalis* | Jonathon Jongsma | <https://www.xeno-canto.org/254619> |
| 253552 | *Empidonax occidentalis* | Eric DeFonso | <https://www.xeno-canto.org/253552> |
| 330648 | *Empidonax difficilis* | Ken Chamberlain | <https://www.xeno-canto.org/330648> |
| 181813 | *Empidonax difficilis* | Skip Russell | <https://www.xeno-canto.org/181813> |
| 334630 | *Empidonax difficilis* | Scott Gravette | <https://www.xeno-canto.org/334630> |
| 225903 | *Myiarchus crinitus* | Peter Boesman | <https://www.xeno-canto.org/225903> |
| 312459 | *Myiarchus crinitus* | Paul Marvin | <https://www.xeno-canto.org/312459> |
| 132678 | *Myiarchus cinerascens* | Dan Lane | <https://www.xeno-canto.org/132678> |
| 348126 | *Myiarchus cinerascens* | Richard E. Webster | <https://www.xeno-canto.org/348126> |
| 376756 | *Tyrannus vociferans* | Manuel Grosselet | <https://www.xeno-canto.org/376756> |
| 172942 | *Tyrannus vociferans* | Eric DeFonso | <https://www.xeno-canto.org/172942> |
| 140649 | *Tyrannus tyrannus* | Paul Driver | <https://www.xeno-canto.org/140649> |
| 132630 | *Tyrannus tyrannus* | Dan Lane | <https://www.xeno-canto.org/132630> |
| 82743 | *Tyrannus tyrannus* | Ryan P. O’Donnell | <https://www.xeno-canto.org/82743> |
| 134875 | *Tyrannus tyrannus* | Jonathon Jongsma | <https://www.xeno-canto.org/134875> |
| 132677 | *Tyrannus forficatus* | Dan Lane | <https://www.xeno-canto.org/132677> |
| 147499 | *Tyrannus forficatus* | Paul Marvin | <https://www.xeno-canto.org/147499> |
| 316205 | *Tyrannus verticalis* | Bobby Wilcox | <https://www.xeno-canto.org/316205> |
| 184924 | *Tyrannus verticalis* | Richard E. Webster | <https://www.xeno-canto.org/184924> |
| 154313 | *Psilorhinus morio* | Bernard Fort | <https://www.xeno-canto.org/154313> |
| 351151 | *Psilorhinus morio* | Frank Lambert | <https://www.xeno-canto.org/351151> |
| 368411 | *Cyanocorax luxuosus* | Dan Lane | <https://www.xeno-canto.org/368411> |
| 278387 | *Cyanocorax luxuosus* | Paul Marvin | <https://www.xeno-canto.org/278387> |
| 116374 | *Cyanocitta cristata* | Ryan O’Donnell | <https://www.xeno-canto.org/116374> |
| 234669 | *Cyanocitta cristata* | Danny Zapata-Henao | <https://www.xeno-canto.org/234669> |
| 361830 | *Cyanocitta stelleri* | Frank Lambert | <https://www.xeno-canto.org/361830> |
| 14433 | *Cyanocitta stelleri* | Andrew Spencer | <https://www.xeno-canto.org/14433> |
| 22007 | *Cyanocitta stelleri* | Chris Parrish | <https://www.xeno-canto.org/22007> |
| 355087 | *Cyanocitta stelleri* | Thomas Magarian | <https://www.xeno-canto.org/355087> |
| 362511 | *Cyanocitta stelleri* | Frank Lambert | <https://www.xeno-canto.org/362511> |
| 41245 | *Cyanocitta stelleri* | Scott Olmstead | <https://www.xeno-canto.org/41245> |
| 360668 | *Pica nuttalli* | Paul Marvin | <https://www.xeno-canto.org/360668> |
| 297461 | *Pica nuttalli* | Ross Gallardy | <https://www.xeno-canto.org/297461> |
| 254491 | *Pica hudsonia* | Jonathon Jongsma | <https://www.xeno-canto.org/254491> |
| 321397 | *Pica hudsonia* | Nick Komar | <https://www.xeno-canto.org/321397> |
| 109766 | *Corvus caurinus* | Andrew Spencer | <https://www.xeno-canto.org/109766> |
| 160058 | *Corvus caurinus* | Ian Cruickshank | <https://www.xeno-canto.org/160058> |
| 124432 | *Corvus brachyrhynchos* | Gabriel Leite | <https://www.xeno-canto.org/124432> |
| 316302 | *Corvus brachyrhynchos* | Sue Riffe | <https://www.xeno-canto.org/316302> |
| 322466 | *Corvus cryptoleucus* | Richard E. Webster | <https://www.xeno-canto.org/322466> |
| 136206 | *Corvus cryptoleucus* | Dan Lane | <https://www.xeno-canto.org/136206> |
| 325124 | *Corvus corax* | Sue Riffe | <https://www.xeno-canto.org/325124> |
| 213233 | *Corvus corax* | Richard Hoyer | <https://www.xeno-canto.org/213233> |
| 213233 | *Corvus corax* | Richard Hoyer | <https://www.xeno-canto.org/213233> |
| 91967 | *Lanius excubitor* | Andrew Spencer | <https://www.xeno-canto.org/91967> |
| 44982 | *Lanius excubitor* | Todd Wilson | <https://www.xeno-canto.org/44982> |
| 319815 | *Lanius excubitor* | Bruce Lagerquist | <https://www.xeno-canto.org/319815> |
| 91968 | *Lanius excubitor* | Andrew Spencer | <https://www.xeno-canto.org/91968> |
| 44981 | *Lanius excubitor* | Todd Wilson | <https://www.xeno-canto.org/44981> |
| 44982 | *Lanius excubitor* | Todd Wilson | <https://www.xeno-canto.org/44982> |
| 307003 | *Lanius ludovicianus* | Ted Floyd | <https://www.xeno-canto.org/307003> |
| 76966 | *Lanius ludovicianus* | Andrew Spencer | <https://www.xeno-canto.org/76966> |
| 76963 | *Lanius ludovicianus* | Andrew Spencer | <https://www.xeno-canto.org/76963> |
| 76964 | *Lanius ludovicianus* | Andrew Spencer | <https://www.xeno-canto.org/76964> |
| 318011 | *Lanius ludovicianus* | Nick Komar | <https://www.xeno-canto.org/318011> |
| 367268 | *Lanius ludovicianus* | Vicki Dern | <https://www.xeno-canto.org/367268> |
| 374109 | *Vireo plumbeus* | Matt Wistrand | <https://www.xeno-canto.org/374109> |
| 170091 | *Vireo plumbeus* | Micah Riegner | <https://www.xeno-canto.org/170091> |
| 323415 | *Vireo plumbeus* | Richard E. Webster | <https://www.xeno-canto.org/323415> |
| 323431 | *Vireo plumbeus* | Richard E. Webster | <https://www.xeno-canto.org/323431> |
| 323424 | *Vireo plumbeus* | Richard E. Webster | <https://www.xeno-canto.org/323424> |
| 14274 | *Vireo plumbeus* | Chris Parrish | <https://www.xeno-canto.org/14274> |
| 76908 | *Vireo cassinii* | Stuart Fisher | <https://www.xeno-canto.org/76908> |
| 136030 | *Vireo cassinii* | Brian Sullivan | <https://www.xeno-canto.org/136030> |
| 378430 | *Vireo cassinii* | Jeremy Minns | <https://www.xeno-canto.org/378430> |
| 135027 | *Vireo cassinii* | Tom Forwood Jr. | <https://www.xeno-canto.org/135027> |
| 102950 | *Vireo cassinii* | Eric DeFonso | <https://www.xeno-canto.org/102950> |
| 379329 | *Vireo cassinii* | Lance A. M. Benner | <https://www.xeno-canto.org/379329> |
| 376564 | *Vireo gilvus* | Michael Harrison | <https://www.xeno-canto.org/376564> |
| 14276 | *Vireo gilvus* | Chris Parrish | <https://www.xeno-canto.org/14276> |
| 297175 | *Vireo gilvus* | Paul Marvin | <https://www.xeno-canto.org/297175> |
| 297175 | *Vireo gilvus* | Paul Marvin | <https://www.xeno-canto.org/297175> |
| 102968 | *Vireo gilvus* | Jonathon Jongsma | <https://www.xeno-canto.org/102968> |
| 325352 | *Setophaga aestiva* | Sue Riffe | <https://www.xeno-canto.org/325352> |
| 137608 | *Vireo philadelphicus* | Randy Dzenkiw | <https://www.xeno-canto.org/137608> |
| 317596 | *Vireo philadelphicus* | Matt Wistrand | <https://www.xeno-canto.org/317596> |
| 180710 | *Vireo philadelphicus* | Ian Cruickshank | <https://www.xeno-canto.org/180710> |
| 189440 | *Vireo philadelphicus* | Andrew Spencer | <https://www.xeno-canto.org/189440> |
| 298975 | *Vireo philadelphicus* | Martin St-Michel | <https://www.xeno-canto.org/298975> |
| 317596 | *Vireo philadelphicus* | Matt Wistrand | <https://www.xeno-canto.org/317596> |
| 160849 | *Poecile gambeli* | Ian Cruickshank | <https://www.xeno-canto.org/160849> |
| 240242 | *Poecile gambeli* | Carrie Branch | <https://www.xeno-canto.org/240242> |
| 352827 | *Poecile atricapillus* | Ted Floyd | <https://www.xeno-canto.org/352827> |
| 335759 | *Poecile atricapillus* | Thomas Magarian | <https://www.xeno-canto.org/335759> |
| 34320 | *Baeolophus atricristatus* | Andrew Spencer | <https://www.xeno-canto.org/34320> |
| 147279 | *Baeolophus atricristatus* | Paul Marvin | <https://www.xeno-canto.org/147279> |
| 21965 | *Baeolophus atricristatus* | Chris Parrish | <https://www.xeno-canto.org/21965> |
| 77331 | *Baeolophus atricristatus* | Chris Harrison | <https://www.xeno-canto.org/77331> |
| 33596 | *Baeolophus bicolor* | Andrew Spencer | <https://www.xeno-canto.org/33596> |
| 233656 | *Baeolophus bicolor* | David Sarkozi | <https://www.xeno-canto.org/233656> |
| 370491 | *Baeolophus bicolor* | Eric DeFonso | <https://www.xeno-canto.org/370491> |
| 33585 | *Baeolophus bicolor* | Andrew Spencer | <https://www.xeno-canto.org/33585> |
| 360933 | *Baeolophus bicolor* | Matt Brady | <https://www.xeno-canto.org/360933> |
| 97374 | *Baeolophus bicolor* | Daniel Lane | <https://www.xeno-canto.org/97374> |
| 350982 | *Baeolophus inornatus* | Richard E. Webster | <https://www.xeno-canto.org/350982> |
| 350922 | *Baeolophus inornatus* | Richard E. Webster | <https://www.xeno-canto.org/350922> |
| 350941 | *Baeolophus inornatus* | Richard E. Webster | <https://www.xeno-canto.org/350941> |
| 350943 | *Baeolophus inornatus* | Richard E. Webster | <https://www.xeno-canto.org/350943> |
| 350899 | *Baeolophus ridgwayi* | Richard E. Webster | <https://www.xeno-canto.org/350899> |
| 350901 | *Baeolophus ridgwayi* | Richard E. Webster | <https://www.xeno-canto.org/350901> |
| 350899 | *Baeolophus ridgwayi* | Richard E. Webster | <https://www.xeno-canto.org/350899> |
| 350902 | *Baeolophus ridgwayi* | Richard E. Webster | <https://www.xeno-canto.org/350902> |
| 361256 | *Alauda arvensis* | Elias A. Ryberg | <https://www.xeno-canto.org/361256> |
| 302578 | *Alauda arvensis* | James Bradley | <https://www.xeno-canto.org/302578> |
| 361649 | *Alauda arvensis* | Krzysztof Deoniziak | <https://www.xeno-canto.org/361649> |
| 285581 | *Alauda arvensis* | David M. | <https://www.xeno-canto.org/285581> |
| 34861 | *Eremophila alpestris* | Tayler Brooks | <https://www.xeno-canto.org/34861> |
| 345553 | *Eremophila alpestris* | Thomas Magarian | <https://www.xeno-canto.org/345553> |
| 277952 | *Eremophila alpestris* | Paul Marvin | <https://www.xeno-canto.org/277952> |
| 378584 | *Eremophila alpestris* | Rachel Hudson | <https://www.xeno-canto.org/378584> |
| 113685 | *Phylloscopus examinandus* | Adria Sole | <https://www.xeno-canto.org/113685> |
| 340441 | *Phylloscopus borealis* | Tom Wulf | <https://www.xeno-canto.org/340441> |
| 40211 | *Phylloscopus examinandus* | Klaas Felix Jachmann | <https://www.xeno-canto.org/40211> |
| 140994 | *Phylloscopus borealis* | Andrew Spencer | <https://www.xeno-canto.org/140994> |
| 215758 | *Psaltriparus minimus* | Ted Floyd | <https://www.xeno-canto.org/215758> |
| 358701 | *Psaltriparus minimus* | Ted Floyd | <https://www.xeno-canto.org/358701> |
| 34112 | *Petrochelidon fulva* | Andrew Spencer | <https://www.xeno-canto.org/34112> |
| 34112 | *Petrochelidon fulva* | Andrew Spencer | <https://www.xeno-canto.org/34112> |
| 203606 | *Petrochelidon pyrrhonota* | Andrew Spencer | <https://www.xeno-canto.org/203606> |
| 317115 | *Petrochelidon pyrrhonota* | Hal Mitchell | <https://www.xeno-canto.org/317115> |
| 140266 | *Tachycineta thalassina* | Paul Driver | <https://www.xeno-canto.org/140266> |
| 105332 | *Tachycineta thalassina* | Andrew Spencer | <https://www.xeno-canto.org/105332> |
| 22009 | *Tachycineta thalassina* | Chris Parrish | <https://www.xeno-canto.org/22009> |
| 105332 | *Tachycineta thalassina* | Andrew Spencer | <https://www.xeno-canto.org/105332> |
| 163176 | *Tachycineta bicolor* | Paul Marvin | <https://www.xeno-canto.org/163176> |
| 313481 | *Tachycineta bicolor* | Matt Wistrand | <https://www.xeno-canto.org/313481> |
| 299635 | *Tachycineta bicolor* | Paul Marvin | <https://www.xeno-canto.org/299635> |
| 322861 | *Tachycineta bicolor* | Peter Boesman | <https://www.xeno-canto.org/322861> |
| 33563 | *Progne subis* | Andrew Spencer | <https://www.xeno-canto.org/33563> |
| 278201 | *Progne subis* | Patrick Turgeon | <https://www.xeno-canto.org/278201> |
| 302415 | *Stelgidopteryx serripennis* | James Bradley | <https://www.xeno-canto.org/302415> |
| 123367 | *Stelgidopteryx serripennis* | Chris Harrison | <https://www.xeno-canto.org/123367> |
| 122453 | *Thryomanes bewickii* | Chris Harrison | <https://www.xeno-canto.org/122453> |
| 141351 | *Thryomanes bewickii* | Mike Nelson | <https://www.xeno-canto.org/141351> |
| 217813 | *Thryomanes bewickii* | Paul Marvin | <https://www.xeno-canto.org/217813> |
| 70970 | *Thryomanes bewickii* | Mary Beth Stowe | <https://www.xeno-canto.org/70970> |
| 124068 | *Thryothorus ludovicianus* | Chris Harrison | <https://www.xeno-canto.org/124068> |
| 122452 | *Thryothorus ludovicianus* | Chris Harrison | <https://www.xeno-canto.org/122452> |
| 70982 | *Thryothorus ludovicianus* | Mary Beth Stowe | <https://www.xeno-canto.org/70982> |
| 236014 | *Thryothorus ludovicianus* | Danny Zapata-Henao | <https://www.xeno-canto.org/236014> |
| 137656 | *Cistothorus stellaris* | Patrick Turgeon | <https://www.xeno-canto.org/137656> |
| 151605 | *Cistothorus stellaris* | Randy Dzenkiw | <https://www.xeno-canto.org/151605> |
| 137656 | *Cistothorus stellaris* | Patrick Turgeon | <https://www.xeno-canto.org/137656> |
| 137656 | *Cistothorus stellaris* | Patrick Turgeon | <https://www.xeno-canto.org/137656> |
| 230753 | *Cistothorus stellaris* | Paul Marvin | <https://www.xeno-canto.org/230753> |
| 230753 | *Cistothorus stellaris* | Paul Marvin | <https://www.xeno-canto.org/230753> |
| 153248 | *Cistothorus palustris* | Bird Studies Canada - Prairie and Parkland Marsh Monitoring Program (PPMMP) | <https://www.xeno-canto.org/153248> |
| 110106 | *Cistothorus palustris* | Andrew Spencer | <https://www.xeno-canto.org/110106> |
| 153248 | *Cistothorus palustris* | Bird Studies Canada - Prairie and Parkland Marsh Monitoring Program (PPMMP) | <https://www.xeno-canto.org/153248> |
| 153248 | *Cistothorus palustris* | Bird Studies Canada - Prairie and Parkland Marsh Monitoring Program (PPMMP) | <https://www.xeno-canto.org/153248> |
| 110106 | *Cistothorus palustris* | Andrew Spencer | <https://www.xeno-canto.org/110106> |
| 14863 | *Cistothorus palustris* | Andrew Spencer | <https://www.xeno-canto.org/14863> |
| 279681 | *Catherpes mexicanus* | Paul Marvin | <https://www.xeno-canto.org/279681> |
| 297532 | *Catherpes mexicanus* | Ross Gallardy | <https://www.xeno-canto.org/297532> |
| 373820 | *Salpinctes obsoletus* | Jeff Dyck | <https://www.xeno-canto.org/373820> |
| 325011 | *Salpinctes obsoletus* | Jim Holmes | <https://www.xeno-canto.org/325011> |
| 325011 | *Salpinctes obsoletus* | Jim Holmes | <https://www.xeno-canto.org/325011> |
| 229185 | *Salpinctes obsoletus* | Peter Boesman | <https://www.xeno-canto.org/229185> |
| 147295 | *Polioptila californica* | Paul Marvin | <https://www.xeno-canto.org/147295> |
| 203741 | *Polioptila californica* | Andrew Spencer | <https://www.xeno-canto.org/203741> |
| 71907 | *Polioptila melanura* | Richard E Webster | <https://www.xeno-canto.org/71907> |
| 361932 | *Polioptila melanura* | Paul Marvin | <https://www.xeno-canto.org/361932> |
| 348158 | *Polioptila melanura* | Richard E. Webster | <https://www.xeno-canto.org/348158> |
| 361929 | *Polioptila melanura* | Paul Marvin | <https://www.xeno-canto.org/361929> |
| 303354 | *Sitta pygmaea* | Ted Floyd | <https://www.xeno-canto.org/303354> |
| 343422 | *Sitta pygmaea* | Bruce Lagerquist | <https://www.xeno-canto.org/343422> |
| 161393 | *Sitta pygmaea* | Paul Marvin | <https://www.xeno-canto.org/161393> |
| 363167 | *Sitta pygmaea* | Frank Lambert | <https://www.xeno-canto.org/363167> |
| 192108 | *Sitta pusilla* | Paul Marvin | <https://www.xeno-canto.org/192108> |
| 138619 | *Sitta pusilla* | Paul Marvin | <https://www.xeno-canto.org/138619> |
| 33523 | *Sitta pusilla* | Andrew Spencer | <https://www.xeno-canto.org/33523> |
| 152170 | *Sitta pusilla* | Dan Lane | <https://www.xeno-canto.org/152170> |
| 75118 | *Oreoscoptes montanus* | Ryan P. O’Donnell | <https://www.xeno-canto.org/75118> |
| 12118 | *Oreoscoptes montanus* | Nathan Pieplow | <https://www.xeno-canto.org/12118> |
| 320563 | *Oreoscoptes montanus* | Ted Floyd | <https://www.xeno-canto.org/320563> |
| 13841 | *Oreoscoptes montanus* | Andrew Spencer | <https://www.xeno-canto.org/13841> |
| 321942 | *Mimus polyglottos* | Richard E. Webster | <https://www.xeno-canto.org/321942> |
| 321923 | *Mimus polyglottos* | Richard E. Webster | <https://www.xeno-canto.org/321923> |
| 172494 | *Mimus polyglottos* | Eric DeFonso | <https://www.xeno-canto.org/172494> |
| 21733 | *Mimus polyglottos* | Chris Parrish | <https://www.xeno-canto.org/21733> |
| 255629 | *Toxostoma crissale* | Richard E. Webster | <https://www.xeno-canto.org/255629> |
| 231467 | *Toxostoma crissale* | Peter Boesman | <https://www.xeno-canto.org/231467> |
| 255759 | *Toxostoma crissale* | Richard E. Webster | <https://www.xeno-canto.org/255759> |
| 301084 | *Toxostoma crissale* | Ed Pandolfino | <https://www.xeno-canto.org/301084> |
| 17826 | *Toxostoma lecontei* | Nathan Pieplow | <https://www.xeno-canto.org/17826> |
| 28253 | *Toxostoma lecontei* | Andrew Spencer | <https://www.xeno-canto.org/28253> |
| 141432 | *Toxostoma lecontei* | Paul Marvin | <https://www.xeno-canto.org/141432> |
| 141433 | *Toxostoma lecontei* | Paul Marvin | <https://www.xeno-canto.org/141433> |
| 269473 | *Toxostoma rufum* | Paul Marvin | <https://www.xeno-canto.org/269473> |
| 231517 | *Toxostoma rufum* | Peter Boesman | <https://www.xeno-canto.org/231517> |
| 104665 | *Toxostoma rufum* | Andrew Spencer | <https://www.xeno-canto.org/104665> |
| 13184 | *Toxostoma rufum* | Andrew Spencer | <https://www.xeno-canto.org/13184> |
| 21656 | *Toxostoma longirostre* | Chris Parrish | <https://www.xeno-canto.org/21656> |
| 5077 | *Toxostoma longirostre* | Nathan Pieplow | <https://www.xeno-canto.org/5077> |
| 368455 | *Toxostoma longirostre* | Dan Lane | <https://www.xeno-canto.org/368455> |
| 176252 | *Toxostoma longirostre* | Ted Floyd | <https://www.xeno-canto.org/176252> |
| 205426 | *Sialia currucoides* | Eric DeFonso | <https://www.xeno-canto.org/205426> |
| 100904 | *Sialia currucoides* | Andrew Spencer | <https://www.xeno-canto.org/100904> |
| 71747 | *Sialia mexicana* | Richard E Webster | <https://www.xeno-canto.org/71747> |
| 33308 | *Sialia mexicana* | Andrew Spencer | <https://www.xeno-canto.org/33308> |
| 184465 | *Catharus fuscescens* | Martin St-Michel | <https://www.xeno-canto.org/184465> |
| 318447 | *Catharus fuscescens* | Jim Berry | <https://www.xeno-canto.org/318447> |
| 356192 | *Catharus minimus* | Frank Lambert | <https://www.xeno-canto.org/356192> |
| 65822 | *Catharus minimus* | Ian Cruickshank | <https://www.xeno-canto.org/65822> |
| 370820 | *Calliope calliope* | Vladimir Shkurov | <https://www.xeno-canto.org/370820> |
| 329588 | *Calliope calliope* | Masato Nagai (Japan) | <https://www.xeno-canto.org/329588> |
| 326518 | *Calliope calliope* | Alex Thomas | <https://www.xeno-canto.org/326518> |
| 329588 | *Calliope calliope* | Masato Nagai (Japan) | <https://www.xeno-canto.org/329588> |
| 376525 | *Calliope calliope* | Lars Edenius | <https://www.xeno-canto.org/376525> |
| 326518 | *Calliope calliope* | Alex Thomas | <https://www.xeno-canto.org/326518> |
| 139593 | *Oenanthe oenanthe* | Andrew Spencer | <https://www.xeno-canto.org/139593> |
| 354835 | *Oenanthe oenanthe* | Frank Lambert | <https://www.xeno-canto.org/354835> |
| 203610 | *Oenanthe oenanthe* | Andrew Spencer | <https://www.xeno-canto.org/203610> |
| 203610 | *Oenanthe oenanthe* | Andrew Spencer | <https://www.xeno-canto.org/203610> |
| 294071 | *Bombycilla cedrorum* | Lance A. M. Benner | <https://www.xeno-canto.org/294071> |
| 375862 | *Bombycilla cedrorum* | Jeremy Welch | <https://www.xeno-canto.org/375862> |
| 319175 | *Bombycilla garrulus* | Martin St-Michel | <https://www.xeno-canto.org/319175> |
| 18968 | *Bombycilla garrulus* | Andrew Spencer | <https://www.xeno-canto.org/18968> |
| 340719 | *Regulus calendula* | Bruce Lagerquist | <https://www.xeno-canto.org/340719> |
| 371123 | *Regulus calendula* | Jeff Dyck | <https://www.xeno-canto.org/371123> |
| 65850 | *Regulus calendula* | Ian Cruickshank | <https://www.xeno-canto.org/65850> |
| 192500 | *Regulus calendula* | Richard E. Webster | <https://www.xeno-canto.org/192500> |
| 189416 | *Regulus satrapa* | Andrew Spencer | <https://www.xeno-canto.org/189416> |
| 269231 | *Regulus satrapa* | Frank Lambert | <https://www.xeno-canto.org/269231> |
| 252878 | *Regulus satrapa* | Iain | <https://www.xeno-canto.org/252878> |
| 104683 | *Regulus satrapa* | Andrew Spencer | <https://www.xeno-canto.org/104683> |
| 302647 | *Haemorhous purpureus* | James Bradley | <https://www.xeno-canto.org/302647> |
| 195203 | *Haemorhous purpureus* | Richard E. Webster | <https://www.xeno-canto.org/195203> |
| 76419 | *Haemorhous purpureus* | Andrew Spencer | <https://www.xeno-canto.org/76419> |
| 376618 | *Haemorhous purpureus* | Bruce Lagerquist | <https://www.xeno-canto.org/376618> |
| 302193 | *Haemorhous cassinii* | James Bradley | <https://www.xeno-canto.org/302193> |
| 365853 | *Haemorhous cassinii* | Bruce Lagerquist | <https://www.xeno-canto.org/365853> |
| 340366 | *Haemorhous cassinii* | Bruce Lagerquist | <https://www.xeno-canto.org/340366> |
| 365853 | *Haemorhous cassinii* | Bruce Lagerquist | <https://www.xeno-canto.org/365853> |
| 161313 | *Spinus lawrencei* | Paul Marvin | <https://www.xeno-canto.org/161313> |
| 328992 | *Spinus lawrencei* | Lance A. M. Benner | <https://www.xeno-canto.org/328992> |
| 161308 | *Spinus lawrencei* | Paul Marvin | <https://www.xeno-canto.org/161308> |
| 71938 | *Spinus lawrencei* | Richard E Webster | <https://www.xeno-canto.org/71938> |
| 144129 | *Spinus psaltria* | Lauren Harter | <https://www.xeno-canto.org/144129> |
| 374080 | *Spinus psaltria* | Matt Wistrand | <https://www.xeno-canto.org/374080> |
| 71954 | *Spinus psaltria* | Richard E Webster | <https://www.xeno-canto.org/71954> |
| 324033 | *Spinus psaltria* | Paul Marvin | <https://www.xeno-canto.org/324033> |
| 140679 | *Acanthis hornemanni* | Andrew Spencer | <https://www.xeno-canto.org/140679> |
| 142535 | *Acanthis hornemanni* | Andrew Spencer | <https://www.xeno-canto.org/142535> |
| 142565 | *Acanthis hornemanni* | Andrew Spencer | <https://www.xeno-canto.org/142565> |
| 140678 | *Acanthis hornemanni* | Andrew Spencer | <https://www.xeno-canto.org/140678> |
| 141708 | *Acanthis flammea* | Andrew Spencer | <https://www.xeno-canto.org/141708> |
| 216985 | *Acanthis flammea* | Janne Bruun | <https://www.xeno-canto.org/216985> |
| 141708 | *Acanthis flammea* | Andrew Spencer | <https://www.xeno-canto.org/141708> |
| 132609 | *Acanthis flammea* | Jarek Matusiak | <https://www.xeno-canto.org/132609> |
| 192152 | *Loxia leucoptera* | Martin St-Michel | <https://www.xeno-canto.org/192152> |
| 15165 | *Loxia leucoptera* | Andrew Spencer | <https://www.xeno-canto.org/15165> |
| 323021 | *Loxia leucoptera* | Peter Boesman | <https://www.xeno-canto.org/323021> |
| 165652 | *Loxia leucoptera* | Davyd Betchkal | <https://www.xeno-canto.org/165652> |
| 327483 | *Loxia curvirostra* | Bruce Lagerquist | <https://www.xeno-canto.org/327483> |
| 76432 | *Loxia curvirostra* | Andrew Spencer | <https://www.xeno-canto.org/76432> |
| 143212 | *Loxia curvirostra* | Andrew Spencer | <https://www.xeno-canto.org/143212> |
| 143214 | *Loxia curvirostra* | Andrew Spencer | <https://www.xeno-canto.org/143214> |
| 293744 | *Calcarius ornatus* | Paul Marvin | <https://www.xeno-canto.org/293744> |
| 161063 | *Calcarius ornatus* | Paul Marvin | <https://www.xeno-canto.org/161063> |
| 70076 | *Calcarius pictus* | Paul Driver | <https://www.xeno-canto.org/70076> |
| 141001 | *Calcarius pictus* | Andrew Spencer | <https://www.xeno-canto.org/141001> |
| 331841 | *Plectrophenax hyperboreus* | John Puschock | <https://www.xeno-canto.org/331841> |
| 331842 | *Plectrophenax hyperboreus* | John Puschock | <https://www.xeno-canto.org/331842> |
| 352930 | *Plectrophenax nivalis* | Frank Lambert | <https://www.xeno-canto.org/352930> |
| 104581 | *Plectrophenax nivalis* | Ryan P. O’Donnell | <https://www.xeno-canto.org/104581> |
| 375266 | *Parkesia noveboracensis* | Susan Hochgraf | <https://www.xeno-canto.org/375266> |
| 30791 | *Parkesia noveboracensis* | Andrew Spencer | <https://www.xeno-canto.org/30791> |
| 55023 | *Parkesia noveboracensis* | Andrew Spencer | <https://www.xeno-canto.org/55023> |
| 176656 | *Parkesia motacilla* | David Swain | <https://www.xeno-canto.org/176656> |
| 175483 | *Parkesia motacilla* | Peter Wilton | <https://www.xeno-canto.org/175483> |
| 253571 | *Geothlypis philadelphia* | Martin St-Michel | <https://www.xeno-canto.org/253571> |
| 142746 | *Geothlypis philadelphia* | Martin St-Michel | <https://www.xeno-canto.org/142746> |
| 79659 | *Geothlypis tolmiei* | Ian Cruickshank | <https://www.xeno-canto.org/79659> |
| 379004 | *Geothlypis tolmiei* | James Bradley | <https://www.xeno-canto.org/379004> |
| 78891 | *Vermivora cyanoptera* | Jonathon Jongsma | <https://www.xeno-canto.org/78891> |
| 320488 | *Vermivora cyanoptera* | Jonathon Jongsma | <https://www.xeno-canto.org/320488> |
| 103849 | *Vermivora chrysoptera* | Todd Wilson | <https://www.xeno-canto.org/103849> |
| 103587 | *Vermivora chrysoptera* | Andrew Spencer | <https://www.xeno-canto.org/103587> |
| 323617 | *Leiothlypis luciae* | Richard E. Webster | <https://www.xeno-canto.org/323617> |
| 323622 | *Leiothlypis luciae* | Richard E. Webster | <https://www.xeno-canto.org/323622> |
| 175370 | *Leiothlypis virginiae* | Micah Riegner | <https://www.xeno-canto.org/175370> |
| 133180 | *Leiothlypis virginiae* | Micah Riegner | <https://www.xeno-canto.org/133180> |
| 79502 | *Setophaga striata* | Andrew Spencer | <https://www.xeno-canto.org/79502> |
| 330578 | *Setophaga striata* | Martin St-Michel | <https://www.xeno-canto.org/330578> |
| 324946 | *Setophaga striata* | Martin St-Michel | <https://www.xeno-canto.org/324946> |
| 219836 | *Bucco capensis* | Peter Boesman | <https://www.xeno-canto.org/219836> |
| 78900 | *Setophaga castanea* | Andrew Spencer | <https://www.xeno-canto.org/78900> |
| 137479 | *Setophaga castanea* | Martin St-Michel | <https://www.xeno-canto.org/137479> |
| 133258 | *Setophaga castanea* | Fernand Deroussen | <https://www.xeno-canto.org/133258> |
| 253753 | *Setophaga castanea* | Martin St-Michel | <https://www.xeno-canto.org/253753> |
| 50445 | *Setophaga aestiva* | Ian Davies | <https://www.xeno-canto.org/50445> |
| 316039 | *Setophaga aestiva* | John Baur | <https://www.xeno-canto.org/316039> |
| 243422 | *Setophaga aestiva* | Maxime Aubert | <https://www.xeno-canto.org/243422> |
| 292262 | *Setophaga aestiva* | Martin St-Michel | <https://www.xeno-canto.org/292262> |
| 330193 | *Setophaga pensylvanica* | Martin St-Michel | <https://www.xeno-canto.org/330193> |
| 78815 | *Setophaga pensylvanica* | Andrew Spencer | <https://www.xeno-canto.org/78815> |
| 83530 | *Setophaga pensylvanica* | Jelmer Poelstra | <https://www.xeno-canto.org/83530> |
| 52446 | *Setophaga pensylvanica* | Andrew Spencer | <https://www.xeno-canto.org/52446> |
| 13586 | *Setophaga americana* | Chris Parrish | <https://www.xeno-canto.org/13586> |
| 187645 | *Setophaga americana* | Paul Driver | <https://www.xeno-canto.org/187645> |
| 134971 | *Setophaga americana* | L G Price | <https://www.xeno-canto.org/134971> |
| 171645 | *Setophaga americana* | Dan Lane | <https://www.xeno-canto.org/171645> |
| 29780 | *Setophaga pitiayumi* | Daniel Lane | <https://www.xeno-canto.org/29780> |
| 147503 | *Setophaga pitiayumi* | Paul Marvin | <https://www.xeno-canto.org/147503> |
| 100178 | *Setophaga pitiayumi* | Chris Benesh | <https://www.xeno-canto.org/100178> |
| 29781 | *Setophaga pitiayumi* | Daniel Lane | <https://www.xeno-canto.org/29781> |
| 30785 | *Setophaga virens* | Andrew Spencer | <https://www.xeno-canto.org/30785> |
| 59121 | *Setophaga virens* | Tayler Brooks | <https://www.xeno-canto.org/59121> |
| 138633 | *Setophaga virens* | Jonathon Jongsma | <https://www.xeno-canto.org/138633> |
| 237743 | *Setophaga virens* | Dan Lane | <https://www.xeno-canto.org/237743> |
| 64462 | *Setophaga chrysoparia* | Chris Warren | <https://www.xeno-canto.org/64462> |
| 34946 | *Setophaga chrysoparia* | Andrew Spencer | <https://www.xeno-canto.org/34946> |
| 171977 | *Setophaga chrysoparia* | Ronnie Kramer | <https://www.xeno-canto.org/171977> |
| 297557 | *Setophaga chrysoparia* | Ross Gallardy | <https://www.xeno-canto.org/297557> |
| 269030 | *Setophaga townsendi* | Frank Lambert | <https://www.xeno-canto.org/269030> |
| 76413 | *Setophaga townsendi* | Andrew Spencer | <https://www.xeno-canto.org/76413> |
| 36563 | *Setophaga townsendi* | Tayler Brooks | <https://www.xeno-canto.org/36563> |
| 134887 | *Setophaga townsendi* | Tom Forwood Jr. | <https://www.xeno-canto.org/134887> |
| 143150 | *Setophaga occidentalis* | Andrew Spencer | <https://www.xeno-canto.org/143150> |
| 143037 | *Setophaga occidentalis* | Andrew Spencer | <https://www.xeno-canto.org/143037> |
| 143147 | *Setophaga occidentalis* | Andrew Spencer | <https://www.xeno-canto.org/143147> |
| 104725 | *Setophaga occidentalis* | Walter Szeliga | <https://www.xeno-canto.org/104725> |
| 34652 | *Setophaga graciae* | Andrew Spencer | <https://www.xeno-canto.org/34652> |
| 333564 | *Setophaga graciae* | Richard E. Webster | <https://www.xeno-canto.org/333564> |
| 153516 | *Setophaga graciae* | Paul Marvin | <https://www.xeno-canto.org/153516> |
| 235007 | *Setophaga graciae* | Garrett MacDonald | <https://www.xeno-canto.org/235007> |
| 56222 | *Setophaga nigrescens* | Scott Olmstead | <https://www.xeno-canto.org/56222> |
| 293949 | *Setophaga nigrescens* | Patrick Turgeon | <https://www.xeno-canto.org/293949> |
| 175373 | *Setophaga nigrescens* | Micah Riegner | <https://www.xeno-canto.org/175373> |
| 364282 | *Setophaga nigrescens* | Ken Blankenship | <https://www.xeno-canto.org/364282> |
| 79605 | *Cardellina pusilla* | Andrew Spencer | <https://www.xeno-canto.org/79605> |
| 55017 | *Cardellina pusilla* | Andrew Spencer | <https://www.xeno-canto.org/55017> |
| 102395 | *Cardellina rubrifrons* | Scott Olmstead | <https://www.xeno-canto.org/102395> |
| 34328 | *Cardellina rubrifrons* | Andrew Spencer | <https://www.xeno-canto.org/34328> |
| 299339 | *Cardellina rubrifrons* | Lauren Harter | <https://www.xeno-canto.org/299339> |
| 324538 | *Cardellina rubrifrons* | Richard E. Webster | <https://www.xeno-canto.org/324538> |
| 342584 | *Icterus bullockii* | Bruce Lagerquist | <https://www.xeno-canto.org/342584> |
| 34869 | *Icterus bullockii* | Tayler Brooks | <https://www.xeno-canto.org/34869> |
| 185570 | *Icterus galbula* | Richard E. Webster | <https://www.xeno-canto.org/185570> |
| 185570 | *Icterus galbula* | Richard E. Webster | <https://www.xeno-canto.org/185570> |
| 178046 | *Icterus galbula* | Jonathon Jongsma | <https://www.xeno-canto.org/178046> |
| 134872 | *Icterus galbula* | Jonathon Jongsma | <https://www.xeno-canto.org/134872> |
| 278197 | *Sonus naturalis* | Patrick Turgeon | <https://www.xeno-canto.org/278197> |
| 160813 | *Icterus galbula* | Ian Cruickshank | <https://www.xeno-canto.org/160813> |
| 71058 | *Icterus parisorum* | Mary Beth Stowe | <https://www.xeno-canto.org/71058> |
| 34323 | *Icterus parisorum* | Andrew Spencer | <https://www.xeno-canto.org/34323> |
| 71057 | *Icterus parisorum* | Mary Beth Stowe | <https://www.xeno-canto.org/71057> |
| 71057 | *Icterus parisorum* | Mary Beth Stowe | <https://www.xeno-canto.org/71057> |
| 224636 | *Icterus parisorum* | Peter Boesman | <https://www.xeno-canto.org/224636> |
| 224638 | *Icterus parisorum* | Peter Boesman | <https://www.xeno-canto.org/224638> |
| 358991 | *Icterus graduacauda* | Paul Marvin | <https://www.xeno-canto.org/358991> |
| 330587 | *Icterus graduacauda* | Andrew Spencer | <https://www.xeno-canto.org/330587> |
| 330605 | *Icterus graduacauda* | Andrew Spencer | <https://www.xeno-canto.org/330605> |
| 224599 | *Icterus graduacauda* | Peter Boesman | <https://www.xeno-canto.org/224599> |
| 360193 | *Icterus graduacauda* | Paul Marvin | <https://www.xeno-canto.org/360193> |
| 190409 | *Icterus graduacauda* | Andrew Spencer | <https://www.xeno-canto.org/190409> |
| 286430 | *Icterus spurius* | Paul Marvin | <https://www.xeno-canto.org/286430> |
| 173791 | *Icterus spurius* | Paul Marvin | <https://www.xeno-canto.org/173791> |
| 173791 | *Icterus spurius* | Paul Marvin | <https://www.xeno-canto.org/173791> |
| 173791 | *Icterus spurius* | Paul Marvin | <https://www.xeno-canto.org/173791> |
| 56323 | *Icterus spurius* | Chuck Davis | <https://www.xeno-canto.org/56323> |
| 56323 | *Icterus spurius* | Chuck Davis | <https://www.xeno-canto.org/56323> |
| 371807 | *Icterus cucullatus* | Manuel Grosselet | <https://www.xeno-canto.org/371807> |
| 371807 | *Icterus cucullatus* | Manuel Grosselet | <https://www.xeno-canto.org/371807> |
| 299105 | *Icterus cucullatus* | Paul Marvin | <https://www.xeno-canto.org/299105> |
| 299105 | *Icterus cucullatus* | Paul Marvin | <https://www.xeno-canto.org/299105> |
| 313460 | *Icterus cucullatus* | Manuel Grosselet | <https://www.xeno-canto.org/313460> |
| 371807 | *Icterus cucullatus* | Manuel Grosselet | <https://www.xeno-canto.org/371807> |
| 275165 | *Molothrus bonariensis* | Robert S. Ridgely | <https://www.xeno-canto.org/275165> |
| 364743 | *Molothrus bonariensis* | Paul Marvin | <https://www.xeno-canto.org/364743> |
| 315047 | *Molothrus bonariensis* | Paul Marvin | <https://www.xeno-canto.org/315047> |
| 197319 | *Molothrus bonariensis* | Paul Marvin | <https://www.xeno-canto.org/197319> |
| 131566 | *Molothrus ater* | Eric DeFonso | <https://www.xeno-canto.org/131566> |
| 160985 | *Molothrus ater* | Paul Marvin | <https://www.xeno-canto.org/160985> |
| 197272 | *Molothrus ater* | Paul Marvin | <https://www.xeno-canto.org/197272> |
| 160987 | *Molothrus ater* | Paul Marvin | <https://www.xeno-canto.org/160987> |
| 358848 | *Agelaius tricolor* | Jim Holmes | <https://www.xeno-canto.org/358848> |
| 162048 | *Agelaius tricolor* | Paul Marvin | <https://www.xeno-canto.org/162048> |
| 191985 | *Agelaius phoeniceus* | Patrick Turgeon | <https://www.xeno-canto.org/191985> |
| 167989 | *Agelaius phoeniceus* | Ian Cruickshank | <https://www.xeno-canto.org/167989> |
| 164257 | *Agelaius phoeniceus* | Ian Cruickshank | <https://www.xeno-canto.org/164257> |
| 125253 | *Agelaius phoeniceus* | Thomas G. Graves | <https://www.xeno-canto.org/125253> |
| 307865 | *Agelaius phoeniceus* | Thomas G. Graves | <https://www.xeno-canto.org/307865> |
| 363449 | *Agelaius phoeniceus* | Thomas G. Graves | <https://www.xeno-canto.org/363449> |
| 64532 | *Quiscalus major* | Mike Nelson | <https://www.xeno-canto.org/64532> |
| 317042 | *Quiscalus major* | Jim Holmes | <https://www.xeno-canto.org/317042> |
| 317165 | *Quiscalus major* | Dan Lane | <https://www.xeno-canto.org/317165> |
| 254941 | *Quiscalus major* | Nick Komar | <https://www.xeno-canto.org/254941> |
| 176148 | *Quiscalus mexicanus* | Ted Floyd | <https://www.xeno-canto.org/176148> |
| 163159 | *Quiscalus mexicanus* | Paul Marvin | <https://www.xeno-canto.org/163159> |
| 163159 | *Quiscalus mexicanus* | Paul Marvin | <https://www.xeno-canto.org/163159> |
| 163159 | *Quiscalus mexicanus* | Paul Marvin | <https://www.xeno-canto.org/163159> |
| 378600 | *Euphagus cyanocephalus* | Thomas Magarian | <https://www.xeno-canto.org/378600> |
| 352025 | *Euphagus cyanocephalus* | Bruce Lagerquist | <https://www.xeno-canto.org/352025> |
| 286407 | *Euphagus carolinus* | Bobby Wilcox | <https://www.xeno-canto.org/286407> |
| 65829 | *Euphagus carolinus* | Ian Cruickshank | <https://www.xeno-canto.org/65829> |
| 333314 | *Dolichonyx oryzivorus* | Nathan Hentze | <https://www.xeno-canto.org/333314> |
| 185815 | *Dolichonyx oryzivorus* | Richard E. Webster | <https://www.xeno-canto.org/185815> |
| 110084 | *Xanthocephalus xanthocephalus* | Andrew Spencer | <https://www.xeno-canto.org/110084> |
| 302568 | *Xanthocephalus xanthocephalus* | James Bradley | <https://www.xeno-canto.org/302568> |
| 320486 | *Sturnella magna* | Jonathon Jongsma | <https://www.xeno-canto.org/320486> |
| 320478 | *Sturnella magna* | Jonathon Jongsma | <https://www.xeno-canto.org/320478> |
| 320478 | *Sturnella magna* | Jonathon Jongsma | <https://www.xeno-canto.org/320478> |
| 373464 | *Sturnella magna* | Bobby Wilcox | <https://www.xeno-canto.org/373464> |
| 313791 | *Sturnella magna* | Matt Wistrand | <https://www.xeno-canto.org/313791> |
| 321630 | *Sturnella magna* | Tom Behnfield | <https://www.xeno-canto.org/321630> |
| 76505 | *Sturnella neglecta* | Jonathon Jongsma | <https://www.xeno-canto.org/76505> |
| 55149 | *Sturnella neglecta* | Todd Wilson | <https://www.xeno-canto.org/55149> |
| 14290 | *Sturnella neglecta* | Andrew Spencer | <https://www.xeno-canto.org/14290> |
| 104524 | *Sturnella neglecta* | Jonathon Jongsma | <https://www.xeno-canto.org/104524> |
| 179850 | *Chondestes grammacus* | Ted Floyd | <https://www.xeno-canto.org/179850> |
| 76967 | *Chondestes grammacus* | Andrew Spencer | <https://www.xeno-canto.org/76967> |
| 76967 | *Chondestes grammacus* | Andrew Spencer | <https://www.xeno-canto.org/76967> |
| 184978 | *Chondestes grammacus* | Richard E. Webster | <https://www.xeno-canto.org/184978> |
| 14299 | *Calamospiza melanocorys* | Chris Parrish | <https://www.xeno-canto.org/14299> |
| 192321 | *Calamospiza melanocorys* | Paul Marvin | <https://www.xeno-canto.org/192321> |
| 185132 | *Calamospiza melanocorys* | Richard E. Webster | <https://www.xeno-canto.org/185132> |
| 13604 | *Calamospiza melanocorys* | Andrew Spencer | <https://www.xeno-canto.org/13604> |
| 297494 | *Melozone aberti* | Ross Gallardy | <https://www.xeno-canto.org/297494> |
| 111857 | *Melozone aberti* | Richard E Webster | <https://www.xeno-canto.org/111857> |
| 297494 | *Melozone aberti* | Ross Gallardy | <https://www.xeno-canto.org/297494> |
| 79661 | *Melozone aberti* | Scott Olmstead | <https://www.xeno-canto.org/79661> |
| 376023 | *Melozone aberti* | Antonio Xeira | <https://www.xeno-canto.org/376023> |
| 247614 | *Melozone aberti* | Patrick Turgeon | <https://www.xeno-canto.org/247614> |
| 297483 | *Melozone crissalis* | Ross Gallardy | <https://www.xeno-canto.org/297483> |
| 126358 | *Melozone crissalis* | Richard E. Webster | <https://www.xeno-canto.org/126358> |
| 370508 | *Pipilo erythrophthalmus* | Eric DeFonso | <https://www.xeno-canto.org/370508> |
| 254338 | *Pipilo erythrophthalmus* | Bryan Calk | <https://www.xeno-canto.org/254338> |
| 228170 | *Pipilo erythrophthalmus* | Peter Boesman | <https://www.xeno-canto.org/228170> |
| 78874 | *Pipilo erythrophthalmus* | Jonathon Jongsma | <https://www.xeno-canto.org/78874> |
| 213439 | *Pipilo maculatus* | Oscar Ballesteors | <https://www.xeno-canto.org/213439> |
| 54425 | *Pipilo maculatus* | Mike Nelson | <https://www.xeno-canto.org/54425> |
| 104527 | *Pipilo maculatus* | Jonathon Jongsma | <https://www.xeno-canto.org/104527> |
| 373799 | *Pipilo maculatus* | Patrick Turgeon | <https://www.xeno-canto.org/373799> |
| 320479 | *Ammodramus henslowii* | Jonathon Jongsma | <https://www.xeno-canto.org/320479> |
| 302528 | *Ammodramus henslowii* | Lauren Harter | <https://www.xeno-canto.org/302528> |
| 6758 | *Ammodramus bairdii* | Darrell L. Peterson | <https://www.xeno-canto.org/6758> |
| 103658 | *Ammodramus bairdii* | Andrew Spencer | <https://www.xeno-canto.org/103658> |
| 185576 | *Ammodramus bairdii* | Richard E. Webster | <https://www.xeno-canto.org/185576> |
| 321046 | *Melospiza georgiana* | Michael Harrison | <https://www.xeno-canto.org/321046> |
| 294004 | *Melospiza georgiana* | Martin St-Michel | <https://www.xeno-canto.org/294004> |
| 256458 | *Melospiza lincolnii* | Ian Cruickshank | <https://www.xeno-canto.org/256458> |
| 302413 | *Melospiza lincolnii* | James Bradley | <https://www.xeno-canto.org/302413> |
| 55559 | *Ammodramus caudacutus* | Andrew Spencer | <https://www.xeno-canto.org/55559> |
| 59191 | *Ammodramus caudacutus* | Tayler Brooks | <https://www.xeno-canto.org/59191> |
| 55204 | *Ammodramus nelsoni* | Andrew Spencer | <https://www.xeno-canto.org/55204> |
| 110145 | *Ammodramus nelsoni* | Andrew Spencer | <https://www.xeno-canto.org/110145> |
| 373914 | *Junco phaeonotus* | Scott Olmstead | <https://www.xeno-canto.org/373914> |
| 253912 | *Junco phaeonotus* | Bobby Wilcox | <https://www.xeno-canto.org/253912> |
| 241177 | *Junco phaeonotus* | Jarrod Swackhamer | <https://www.xeno-canto.org/241177> |
| 253909 | *Junco phaeonotus* | Bobby Wilcox | <https://www.xeno-canto.org/253909> |
| 321752 | *Junco phaeonotus* | Richard E. Webster | <https://www.xeno-canto.org/321752> |
| 321764 | *Junco phaeonotus* | Richard E. Webster | <https://www.xeno-canto.org/321764> |
| 76622 | *Junco hyemalis* | Andrew Spencer | <https://www.xeno-canto.org/76622> |
| 314047 | *Junco hyemalis* | Ken Blankenship | <https://www.xeno-canto.org/314047> |
| 111224 | *Junco hyemalis* | Richard E Webster | <https://www.xeno-canto.org/111224> |
| 111225 | *Junco hyemalis* | Richard E Webster | <https://www.xeno-canto.org/111225> |
| 111225 | *Junco hyemalis* | Richard E Webster | <https://www.xeno-canto.org/111225> |
| 169518 | *Junco hyemalis* | Micah Riegner | <https://www.xeno-canto.org/169518> |
| 354817 | *Zonotrichia atricapilla* | Frank Lambert | <https://www.xeno-canto.org/354817> |
| 322859 | *Zonotrichia atricapilla* | Peter Boesman | <https://www.xeno-canto.org/322859> |
| 322889 | *Zonotrichia leucophrys* | Peter Boesman | <https://www.xeno-canto.org/322889> |
| 353498 | *Zonotrichia leucophrys* | Frank Lambert | <https://www.xeno-canto.org/353498> |
| 322623 | *Amphispiza bilineata* | Richard E. Webster | <https://www.xeno-canto.org/322623> |
| 111338 | *Amphispiza bilineata* | Richard E Webster | <https://www.xeno-canto.org/111338> |
| 297533 | *Amphispiza bilineata* | Ross Gallardy | <https://www.xeno-canto.org/297533> |
| 322635 | *Amphispiza bilineata* | Richard E. Webster | <https://www.xeno-canto.org/322635> |
| 34419 | *Amphispiza quinquestriata* | Andrew Spencer | <https://www.xeno-canto.org/34419> |
| 87005 | *Amphispiza quinquestriata* | Daniel Lane | <https://www.xeno-canto.org/87005> |
| 41239 | *Amphispiza quinquestriata* | Scott Olmstead | <https://www.xeno-canto.org/41239> |
| 234106 | *Amphispiza quinquestriata* | James and David Bradley | <https://www.xeno-canto.org/234106> |
| 186399 | *Spizella breweri* | Richard E. Webster | <https://www.xeno-canto.org/186399> |
| 341518 | *Spizella breweri* | Thomas Magarian | <https://www.xeno-canto.org/341518> |
| 142631 | *Spizella pusilla* | Jonathon Jongsma | <https://www.xeno-canto.org/142631> |
| 1379 | *Spizella pusilla* | Robin Carter | <https://www.xeno-canto.org/1379> |
| 160801 | *Spizella pallida* | Ian Cruickshank | <https://www.xeno-canto.org/160801> |
| 186813 | *Spizella pallida* | Richard E. Webster | <https://www.xeno-canto.org/186813> |
| 14415 | *Spizella passerina* | Chris Parrish | <https://www.xeno-canto.org/14415> |
| 188094 | *Spizella passerina* | Richard E. Webster | <https://www.xeno-canto.org/188094> |
| 13646 | *Spizella passerina* | Chris Parrish | <https://www.xeno-canto.org/13646> |
| 189062 | *Spizella passerina* | Richard E. Webster | <https://www.xeno-canto.org/189062> |
| 195836 | *Spizella passerina* | Richard E. Webster | <https://www.xeno-canto.org/195836> |
| 138703 | *Spizella passerina* | Garrett MacDonald | <https://www.xeno-canto.org/138703> |
| 83720 | *Ammodramus savannarum* | James Morgan | <https://www.xeno-canto.org/83720> |
| 246800 | *Ammodramus savannarum* | Eric DeFonso | <https://www.xeno-canto.org/246800> |
| 176151 | *Arremonops rufivirgatus* | Ted Floyd | <https://www.xeno-canto.org/176151> |
| 164249 | *Arremonops rufivirgatus* | Paul Marvin | <https://www.xeno-canto.org/164249> |
| 33516 | *Peucaea aestivalis* | Andrew Spencer | <https://www.xeno-canto.org/33516> |
| 33781 | *Peucaea aestivalis* | Andrew Spencer | <https://www.xeno-canto.org/33781> |
| 152127 | *Peucaea aestivalis* | Dan Lane | <https://www.xeno-canto.org/152127> |
| 135133 | *Peucaea aestivalis* | Dan Lane | <https://www.xeno-canto.org/135133> |
| 111468 | *Peucaea cassinii* | Richard E Webster | <https://www.xeno-canto.org/111468> |
| 360791 | *Peucaea cassinii* | Antonio Xeira | <https://www.xeno-canto.org/360791> |
| 111468 | *Peucaea cassinii* | Richard E Webster | <https://www.xeno-canto.org/111468> |
| 123897 | *Peucaea cassinii* | Chris Harrison | <https://www.xeno-canto.org/123897> |
| 13594 | *Peucaea cassinii* | Andrew Spencer | <https://www.xeno-canto.org/13594> |
| 172649 | *Peucaea cassinii* | Eric DeFonso | <https://www.xeno-canto.org/172649> |
| 14960 | *Passerina caerulea* | Andrew Spencer | <https://www.xeno-canto.org/14960> |
| 191550 | *Passerina caerulea* | Phoenix Birder | <https://www.xeno-canto.org/191550> |
| 375861 | *Passerina amoena* | Jeremy Welch | <https://www.xeno-canto.org/375861> |
| 206004 | *Passerina amoena* | Eric DeFonso | <https://www.xeno-canto.org/206004> |
| 277012 | *Passerina ciris* | Paul Marvin | <https://www.xeno-canto.org/277012> |
| 371810 | *Passerina ciris* | Manuel Grosselet | <https://www.xeno-canto.org/371810> |
| 266317 | *Passerina versicolor* | Richard E. Webster | <https://www.xeno-canto.org/266317> |
| 34436 | *Passerina versicolor* | Andrew Spencer | <https://www.xeno-canto.org/34436> |
| 382102 | *Passerina versicolor* | Manuel Grosselet | <https://www.xeno-canto.org/382102> |
| 6658 | *Passerina versicolor* | Manuel Grosselet | <https://www.xeno-canto.org/6658> |
| 369969 | *Pheucticus melanocephalus* | Eric DeFonso | <https://www.xeno-canto.org/369969> |
| 172821 | *Pheucticus melanocephalus* | Eric DeFonso | <https://www.xeno-canto.org/172821> |
| 109305 | *Pheucticus melanocephalus* | Andrew Spencer | <https://www.xeno-canto.org/109305> |
| 254456 | *Pheucticus melanocephalus* | Jonathon Jongsma | <https://www.xeno-canto.org/254456> |
| 315288 | *Pheucticus ludovicianus* | Antonio Xeira | <https://www.xeno-canto.org/315288> |
| 134880 | *Pheucticus ludovicianus* | Jonathon Jongsma | <https://www.xeno-canto.org/134880> |
| 373511 | *Pheucticus ludovicianus* | Bobby Wilcox | <https://www.xeno-canto.org/373511> |
| 181480 | *Piranga ludoviciana* | Randy Dzenkiw | <https://www.xeno-canto.org/181480> |
| 190053 | *Piranga ludoviciana* | Richard E. Webster | <https://www.xeno-canto.org/190053> |
| 192752 | *Piranga ludoviciana* | Paul Marvin | <https://www.xeno-canto.org/192752> |
| 302200 | *Piranga ludoviciana* | James Bradley | <https://www.xeno-canto.org/302200> |
| 144033 | *Piranga olivacea* | Laura Gooch | <https://www.xeno-canto.org/144033> |
| 179639 | *Piranga olivacea* | Martin St-Michel | <https://www.xeno-canto.org/179639> |
| 291932 | *Piranga olivacea* | Martin St-Michel | <https://www.xeno-canto.org/291932> |
| 17153 | *Piranga olivacea* | Allen T. Chartier | <https://www.xeno-canto.org/17153> |
| 244067 | *Piranga olivacea* | Maxime Aubert | <https://www.xeno-canto.org/244067> |
| 17152 | *Piranga olivacea* | Allen T. Chartier | <https://www.xeno-canto.org/17152> |
| 368457 | *Cardinalis sinuatus* | Dan Lane | <https://www.xeno-canto.org/368457> |
| 322450 | *Cardinalis sinuatus* | Richard E. Webster | <https://www.xeno-canto.org/322450> |
| 382035 | *Cardinalis sinuatus* | Manuel Grosselet | <https://www.xeno-canto.org/382035> |
| 322440 | *Cardinalis sinuatus* | Richard E. Webster | <https://www.xeno-canto.org/322440> |
| 235432 | *Cardinalis cardinalis* | Danny Zapata-Henao | <https://www.xeno-canto.org/235432> |
| 291402 | *Cardinalis cardinalis* | Eric Hough | <https://www.xeno-canto.org/291402> |
| 297495 | *Cardinalis cardinalis* | Ross Gallardy | <https://www.xeno-canto.org/297495> |
| 285316 | *Cardinalis cardinalis* | Paul Marvin | <https://www.xeno-canto.org/285316> |
| 26684 | *Anthus cervinus* | Patrik Åberg | <https://www.xeno-canto.org/26684> |
| 105855 | *Cinnyris solaris* | Colin Trainor | <https://www.xeno-canto.org/105855> |
| 26684 | *Anthus cervinus* | Patrik Åberg | <https://www.xeno-canto.org/26684> |
| 382130 | *Anthus cervinus* | Jens Kirkeby | <https://www.xeno-canto.org/382130> |
| 203607 | *Anthus rubescens* | Andrew Spencer | <https://www.xeno-canto.org/203607> |
| 203608 | *Anthus rubescens* | Andrew Spencer | <https://www.xeno-canto.org/203608> |
| 62832 | *Motacilla flava* | Holger Schielzeth | <https://www.xeno-canto.org/62832> |
| 362774 | *Motacilla flava* | Dimitri | <https://www.xeno-canto.org/362774> |
| 345893 | *Motacilla alba* | Tero Linjama | <https://www.xeno-canto.org/345893> |
| 374719 | *Motacilla alba* | Frank A. Roos | <https://www.xeno-canto.org/374719> |
| 215457 | *Aphelocoma woodhouseii* | Kristie Nelson | <https://www.xeno-canto.org/215457> |
| 290279 | *Aphelocoma woodhouseii* | Manuel Grosselet | <https://www.xeno-canto.org/290279> |
| 215457 | *Aphelocoma woodhouseii* | Kristie Nelson | <https://www.xeno-canto.org/215457> |
| 360967 | *Aphelocoma californica* | Paul Marvin | <https://www.xeno-canto.org/360967> |
| 297455 | *Aphelocoma californica* | Ross Gallardy | <https://www.xeno-canto.org/297455> |
| 349731 | *Aphelocoma californica* | Paul Marvin | <https://www.xeno-canto.org/349731> |
| 352935 | *Troglodytes pacificus* | Frank Lambert | <https://www.xeno-canto.org/352935> |
| 195801 | *Troglodytes pacificus* | Richard E. Webster | <https://www.xeno-canto.org/195801> |
| 193225 | *Troglodytes pacificus* | Richard E. Webster | <https://www.xeno-canto.org/193225> |
| 195054 | *Troglodytes pacificus* | Richard E. Webster | <https://www.xeno-canto.org/195054> |
| 193993 | *Troglodytes pacificus* | Richard E. Webster | <https://www.xeno-canto.org/193993> |
| 195801 | *Troglodytes pacificus* | Richard E. Webster | <https://www.xeno-canto.org/195801> |
| 351174 | *Troglodytes hiemalis* | Richard E. Webster | <https://www.xeno-canto.org/351174> |
| 103884 | *Troglodytes hiemalis* | Todd Wilson | <https://www.xeno-canto.org/103884> |
| 103884 | *Troglodytes hiemalis* | Todd Wilson | <https://www.xeno-canto.org/103884> |
| 188933 | *Troglodytes hiemalis* | Richard E. Webster | <https://www.xeno-canto.org/188933> |
| 377372 | *Troglodytes hiemalis* | Michael Harrison | <https://www.xeno-canto.org/377372> |
| 188931 | *Troglodytes hiemalis* | Richard E. Webster | <https://www.xeno-canto.org/188931> |
| 262814 | *Artemisiospiza nevadensis* | Lance A. M. Benner | <https://www.xeno-canto.org/262814> |
| 364570 | *Artemisiospiza nevadensis* | Jim Holmes | <https://www.xeno-canto.org/364570> |
| 368296 | *Artemisiospiza belli* | Jim Holmes | <https://www.xeno-canto.org/368296> |
| 309039 | *Artemisiospiza belli* | Shannon Skalos | <https://www.xeno-canto.org/309039> |
| 303354 | *Sitta pygmaea* | Ted Floyd | <https://www.xeno-canto.org/303354> |
| 343422 | *Sitta pygmaea* | Bruce Lagerquist | <https://www.xeno-canto.org/343422> |
| 161393 | *Sitta pygmaea* | Paul Marvin | <https://www.xeno-canto.org/161393> |
| 363167 | *Sitta pygmaea* | Frank Lambert | <https://www.xeno-canto.org/363167> |
| 362017 | *Sitta canadensis* | Frank Lambert | <https://www.xeno-canto.org/362017> |
| 269235 | *Sitta canadensis* | Frank Lambert | <https://www.xeno-canto.org/269235> |
| 269235 | *Sitta canadensis* | Frank Lambert | <https://www.xeno-canto.org/269235> |
| 172414 | *Sitta canadensis* | Eric DeFonso | <https://www.xeno-canto.org/172414> |
| 255797 | *Toxostoma curvirostre* | Richard E. Webster | <https://www.xeno-canto.org/255797> |
| 309812 | *Toxostoma curvirostre* | Manuel Grosselet | <https://www.xeno-canto.org/309812> |
| 368460 | *Toxostoma curvirostre* | Dan Lane | <https://www.xeno-canto.org/368460> |
| 371093 | *Toxostoma curvirostre* | Manuel Grosselet | <https://www.xeno-canto.org/371093> |
| 21656 | *Toxostoma longirostre* | Chris Parrish | <https://www.xeno-canto.org/21656> |
| 5077 | *Toxostoma longirostre* | Nathan Pieplow | <https://www.xeno-canto.org/5077> |
| 368455 | *Toxostoma longirostre* | Dan Lane | <https://www.xeno-canto.org/368455> |
| 176252 | *Toxostoma longirostre* | Ted Floyd | <https://www.xeno-canto.org/176252> |
| 253571 | *Geothlypis philadelphia* | Martin St-Michel | <https://www.xeno-canto.org/253571> |
| 142746 | *Geothlypis philadelphia* | Martin St-Michel | <https://www.xeno-canto.org/142746> |
| 252951 | *Oporornis agilis* | Nancy Hetrick | <https://www.xeno-canto.org/252951> |
| 181481 | *Oporornis agilis* | Randy Dzenkiw | <https://www.xeno-canto.org/181481> |
| 244475 | *Molothrus aeneus* | Bobby Wilcox | <https://www.xeno-canto.org/244475> |
| 246087 | *Molothrus aeneus* | Scott Olmstead | <https://www.xeno-canto.org/246087> |
| 333844 | *Molothrus aeneus* | Richard E. Webster | <https://www.xeno-canto.org/333844> |
| 246087 | *Molothrus aeneus* | Scott Olmstead | <https://www.xeno-canto.org/246087> |
| 131566 | *Molothrus ater* | Eric DeFonso | <https://www.xeno-canto.org/131566> |
| 160985 | *Molothrus ater* | Paul Marvin | <https://www.xeno-canto.org/160985> |
| 197272 | *Molothrus ater* | Paul Marvin | <https://www.xeno-canto.org/197272> |
| 160987 | *Molothrus ater* | Paul Marvin | <https://www.xeno-canto.org/160987> |
| 297494 | *Melozone aberti* | Ross Gallardy | <https://www.xeno-canto.org/297494> |
| 111857 | *Melozone aberti* | Richard E Webster | <https://www.xeno-canto.org/111857> |
| 297494 | *Melozone aberti* | Ross Gallardy | <https://www.xeno-canto.org/297494> |
| 79661 | *Melozone aberti* | Scott Olmstead | <https://www.xeno-canto.org/79661> |
| 376023 | *Melozone aberti* | Antonio Xeira | <https://www.xeno-canto.org/376023> |
| 247614 | *Melozone aberti* | Patrick Turgeon | <https://www.xeno-canto.org/247614> |
| 323281 | *Melozone fusca* | Richard E. Webster | <https://www.xeno-canto.org/323281> |
| 257200 | *Melozone fusca* | Scott Olmstead | <https://www.xeno-canto.org/257200> |
| 323281 | *Melozone fusca* | Richard E. Webster | <https://www.xeno-canto.org/323281> |
| 18127 | *Melozone fusca* | Andrew Spencer | <https://www.xeno-canto.org/18127> |
| 297491 | *Peucaea botterii* | Ross Gallardy | <https://www.xeno-canto.org/297491> |
| 196552 | *Peucaea botterii* | Dan Lane | <https://www.xeno-canto.org/196552> |
| 111468 | *Peucaea cassinii* | Richard E Webster | <https://www.xeno-canto.org/111468> |
| 360791 | *Peucaea cassinii* | Antonio Xeira | <https://www.xeno-canto.org/360791> |
| 268679 | *Myiarchus tyrannulus* | Richard E. Webster | <https://www.xeno-canto.org/268679> |
| 35375 | *Myiarchus tyrannulus* | Andrew Spencer | <https://www.xeno-canto.org/35375> |
| 122918 | *Myiarchus cinerascens* | Richard E. Webster | <https://www.xeno-canto.org/122918> |
| 318523 | *Myiarchus cinerascens* | Jarrod Swackhamer | <https://www.xeno-canto.org/318523> |
| 184465 | *Catharus fuscescens* | Martin St-Michel | <https://www.xeno-canto.org/184465> |
| 318447 | *Catharus fuscescens* | Jim Berry | <https://www.xeno-canto.org/318447> |
| 30771 | *Catharus fuscescens* | Andrew Spencer | <https://www.xeno-canto.org/30771> |
| 30771 | *Catharus fuscescens* | Andrew Spencer | <https://www.xeno-canto.org/30771> |
| 293438 | *Catharus fuscescens* | Martin St-Michel | <https://www.xeno-canto.org/293438> |
| 142745 | *Catharus fuscescens* | Martin St-Michel | <https://www.xeno-canto.org/142745> |
| 54830 | *Catharus bicknelli* | Andrew Spencer | <https://www.xeno-canto.org/54830> |
| 160919 | *Catharus bicknelli* | Paul Marvin | <https://www.xeno-canto.org/160919> |
| 79605 | *Cardellina pusilla* | Andrew Spencer | <https://www.xeno-canto.org/79605> |
| 55017 | *Cardellina pusilla* | Andrew Spencer | <https://www.xeno-canto.org/55017> |
| 79614 | *Cardellina canadensis* | Andrew Spencer | <https://www.xeno-canto.org/79614> |
| 186311 | *Cardellina canadensis* | Martin St-Michel | <https://www.xeno-canto.org/186311> |
| 296170 | *Cardellina canadensis* | Paul Marvin | <https://www.xeno-canto.org/296170> |
| 318455 | *Cardellina canadensis* | Jim Berry | <https://www.xeno-canto.org/318455> |
| 55559 | *Ammodramus caudacutus* | Andrew Spencer | <https://www.xeno-canto.org/55559> |
| 59191 | *Ammodramus caudacutus* | Tayler Brooks | <https://www.xeno-canto.org/59191> |
| 59157 | *Ammodramus maritimus* | Tayler Brooks | <https://www.xeno-canto.org/59157> |
| 359417 | *Ammodramus maritimus* | Linda Stehlik | <https://www.xeno-canto.org/359417> |
| 319003 | *Piranga ludoviciana* | Ted Floyd | <https://www.xeno-canto.org/319003> |
| 323204 | *Piranga ludoviciana* | Richard E. Webster | <https://www.xeno-canto.org/323204> |
| 1252 | *Piranga ludoviciana* | Don Jones | <https://www.xeno-canto.org/1252> |
| 14092 | *Piranga ludoviciana* | Andrew Spencer | <https://www.xeno-canto.org/14092> |
| 293628 | *Piranga hepatica* | Paul Marvin | <https://www.xeno-canto.org/293628> |
| 35383 | *Piranga hepatica* | Andrew Spencer | <https://www.xeno-canto.org/35383> |
| 35383 | *Piranga hepatica* | Andrew Spencer | <https://www.xeno-canto.org/35383> |
| 21764 | *Piranga hepatica* | Chris Parrish | <https://www.xeno-canto.org/21764> |
| 65375 | *Pooecetes gramineus* | Eric DeFonso | <https://www.xeno-canto.org/65375> |
| 366656 | *Pooecetes gramineus* | Frank Lambert | <https://www.xeno-canto.org/366656> |
| 125749 | *Pooecetes gramineus* | Richard E. Webster | <https://www.xeno-canto.org/125749> |
| 377801 | *Pooecetes gramineus* | Bobby Wilcox | <https://www.xeno-canto.org/377801> |
| 368296 | *Artemisiospiza belli* | Jim Holmes | <https://www.xeno-canto.org/368296> |
| 309039 | *Artemisiospiza belli* | Shannon Skalos | <https://www.xeno-canto.org/309039> |
| 356271 | *Spizelloides arborea* | Frank Lambert | <https://www.xeno-canto.org/356271> |
| 203711 | *Spizelloides arborea* | Andrew Spencer | <https://www.xeno-canto.org/203711> |
| 353480 | *Passerella iliaca* | Frank Lambert | <https://www.xeno-canto.org/353480> |
| 322857 | *Passerella iliaca* | Peter Boesman | <https://www.xeno-canto.org/322857> |
| 209847 | *Zonotrichia querula* | Terry Davis | <https://www.xeno-canto.org/209847> |
| 142845 | *Zonotrichia querula* | Ian Cruickshank | <https://www.xeno-canto.org/142845> |
| 329881 | *Zonotrichia albicollis* | John Patterson | <https://www.xeno-canto.org/329881> |
| 329835 | *Zonotrichia albicollis* | John Patterson | <https://www.xeno-canto.org/329835> |
| 143941 | *Leucosticte tephrocotis* | Andrew Spencer | <https://www.xeno-canto.org/143941> |
| 143944 | *Leucosticte tephrocotis* | Andrew Spencer | <https://www.xeno-canto.org/143944> |
| 107919 | *Leucosticte tephrocotis* | Jelmer Poelstra | <https://www.xeno-canto.org/107919> |
| 344692 | *Leucosticte tephrocotis* | Thomas Magarian | <https://www.xeno-canto.org/344692> |
| 284845 | *Leucosticte atrata* | Paul Marvin | <https://www.xeno-canto.org/284845> |
| 143900 | *Leucosticte atrata* | Andrew Spencer | <https://www.xeno-canto.org/143900> |
| 143749 | *Leucosticte atrata* | Andrew Spencer | <https://www.xeno-canto.org/143749> |
| 284240 | *Leucosticte atrata* | Paul Marvin | <https://www.xeno-canto.org/284240> |
| 284544 | *Leucosticte atrata* | Paul Marvin | <https://www.xeno-canto.org/284544> |
| 143901 | *Leucosticte atrata* | Andrew Spencer | <https://www.xeno-canto.org/143901> |
| 322979 | *Poecile hudsonicus* | Peter Boesman | <https://www.xeno-canto.org/322979> |
| 46489 | *Poecile hudsonicus* | Andrew Spencer | <https://www.xeno-canto.org/46489> |
| 269229 | *Poecile rufescens* | Frank Lambert | <https://www.xeno-canto.org/269229> |
| 92190 | *Poecile rufescens* | Nathan Pieplow | <https://www.xeno-canto.org/92190> |
| 360934 | *Poecile carolinensis* | Matt Brady | <https://www.xeno-canto.org/360934> |
| 309927 | *Poecile carolinensis* | J.R. Rigby | <https://www.xeno-canto.org/309927> |
| 169225 | *Poecile sclateri* | Paul Marvin | <https://www.xeno-canto.org/169225> |
| 255743 | *Poecile sclateri* | Frank Lambert | <https://www.xeno-canto.org/255743> |
| 34325 | *Poecile sclateri* | Andrew Spencer | <https://www.xeno-canto.org/34325> |
| 317973 | *Poecile sclateri* | Jim Holmes | <https://www.xeno-canto.org/317973> |
| 103759 | *Limnothlypis swainsonii* | Mike Nelson | <https://www.xeno-canto.org/103759> |
| 59134 | *Limnothlypis swainsonii* | Tayler Brooks | <https://www.xeno-canto.org/59134> |
| 33466 | *Protonotaria citrea* | Andrew Spencer | <https://www.xeno-canto.org/33466> |
| 316226 | *Protonotaria citrea* | Bates Estabrooks | <https://www.xeno-canto.org/316226> |
| 59145 | *Geothlypis formosa* | Tayler Brooks | <https://www.xeno-canto.org/59145> |
| 20433 | *Geothlypis formosa* | Chris Parrish | <https://www.xeno-canto.org/20433> |
| 316231 | *Geothlypis trichas* | Antonio Xeira | <https://www.xeno-canto.org/316231> |
| 362478 | *Geothlypis trichas* | Scott Gravette | <https://www.xeno-canto.org/362478> |
| 78803 | *Setophaga ruticilla* | Andrew Spencer | <https://www.xeno-canto.org/78803> |
| 293435 | *Setophaga ruticilla* | Martin St-Michel | <https://www.xeno-canto.org/293435> |
| 179305 | *Setophaga ruticilla* | Martin St-Michel | <https://www.xeno-canto.org/179305> |
| 247312 | *Setophaga ruticilla* | Iain | <https://www.xeno-canto.org/247312> |
| 59119 | *Setophaga caerulescens* | Tayler Brooks | <https://www.xeno-canto.org/59119> |
| 49874 | *Setophaga caerulescens* | Andrew Spencer | <https://www.xeno-canto.org/49874> |
| 133250 | *Setophaga caerulescens* | Fernand DEROUSSEN | <https://www.xeno-canto.org/133250> |
| 186365 | *Setophaga caerulescens* | Martin St-Michel | <https://www.xeno-canto.org/186365> |
| 13586 | *Setophaga americana* | Chris Parrish | <https://www.xeno-canto.org/13586> |
| 187645 | *Setophaga americana* | Paul Driver | <https://www.xeno-canto.org/187645> |
| 134971 | *Setophaga americana* | L G Price | <https://www.xeno-canto.org/134971> |
| 171645 | *Setophaga americana* | Dan Lane | <https://www.xeno-canto.org/171645> |
| 83553 | *Setophaga tigrina* | Jelmer Poelstra | <https://www.xeno-canto.org/83553> |
| 134506 | *Setophaga tigrina* | Martin St-Michel | <https://www.xeno-canto.org/134506> |
| 365856 | *Setophaga dominica* | Matt Wistrand | <https://www.xeno-canto.org/365856> |
| 372014 | *Setophaga dominica* | Eric DeFonso | <https://www.xeno-canto.org/372014> |
| 316232 | *Setophaga dominica* | Antonio Xeira | <https://www.xeno-canto.org/316232> |
| 312741 | *Setophaga dominica* | Aidan Place | <https://www.xeno-canto.org/312741> |
| 81084 | *Setophaga pinus* | Mike Nelson | <https://www.xeno-canto.org/81084> |
| 87973 | *Setophaga pinus* | Mike Nelson | <https://www.xeno-canto.org/87973> |
| 316727 | *Setophaga pinus* | Matt Wistrand | <https://www.xeno-canto.org/316727> |
| 356277 | *Setophaga pinus* | Scott Gravette | <https://www.xeno-canto.org/356277> |
| 299197 | *Turdus obscurus* | Albert Lastukhin & Vadim Ivushkin | <https://www.xeno-canto.org/299197> |
| 299196 | *Turdus obscurus* | Albert Lastukhin & Vadim Ivushkin | <https://www.xeno-canto.org/299196> |
| 113696 | *Turdus obscurus* | Adria Sole | <https://www.xeno-canto.org/113696> |
| 299196 | *Turdus obscurus* | Albert Lastukhin & Vadim Ivushkin | <https://www.xeno-canto.org/299196> |
| 351863 | *Turdus migratorius* | Richard E. Webster | <https://www.xeno-canto.org/351863> |
| 322822 | *Turdus migratorius* | Peter Boesman | <https://www.xeno-canto.org/322822> |
| 322797 | *Turdus migratorius* | Peter Boesman | <https://www.xeno-canto.org/322797> |
| 353500 | *Turdus migratorius* | Frank Lambert | <https://www.xeno-canto.org/353500> |
| 327514 | *Turdus rufopalliatus* | Manuel Grosselet | <https://www.xeno-canto.org/327514> |
| 314225 | *Turdus rufopalliatus* | Manuel Grosselet | <https://www.xeno-canto.org/314225> |
| 317533 | *Turdus rufopalliatus* | Manuel Grosselet | <https://www.xeno-canto.org/317533> |
| 6243 | *Turdus rufopalliatus* | Manuel Grosselet | <https://www.xeno-canto.org/6243> |
| 277327 | *Turdus grayi* | Paul Marvin | <https://www.xeno-canto.org/277327> |
| 277034 | *Turdus grayi* | Paul Marvin | <https://www.xeno-canto.org/277034> |
| 330611 | *Turdus grayi* | Andrew Spencer | <https://www.xeno-canto.org/330611> |
| 147307 | *Turdus grayi* | Paul Marvin | <https://www.xeno-canto.org/147307> |
| 103087 | *Empidonax wrightii* | Eric DeFonso | <https://www.xeno-canto.org/103087> |
| 14088 | *Empidonax wrightii* | Chris Parrish | <https://www.xeno-canto.org/14088> |
| 35909 | *Empidonax minimus* | Tayler Brooks | <https://www.xeno-canto.org/35909> |
| 248003 | *Empidonax minimus* | Ryan P. O’Donnell | <https://www.xeno-canto.org/248003> |
| 359298 | *Tyrannus melancholicus* | Kent Livezey | <https://www.xeno-canto.org/359298> |
| 308701 | *Tyrannus melancholicus* | Mario Trejo | <https://www.xeno-canto.org/308701> |
| 133304 | *Tyrannus couchii* | Andrew Spencer | <https://www.xeno-canto.org/133304> |
| 305522 | *Tyrannus couchii* | Paul Marvin | <https://www.xeno-canto.org/305522> |
| 306945 | *Vireo huttoni* | Scott Olmstead | <https://www.xeno-canto.org/306945> |
| 323411 | *Vireo huttoni* | Richard E. Webster | <https://www.xeno-canto.org/323411> |
| 235061 | *Vireo huttoni* | Garrett MacDonald | <https://www.xeno-canto.org/235061> |
| 235062 | *Vireo huttoni* | Garrett MacDonald | <https://www.xeno-canto.org/235062> |
| 279009 | *Vireo huttoni* | Nick Komar | <https://www.xeno-canto.org/279009> |
| 297530 | *Vireo huttoni* | Ross Gallardy | <https://www.xeno-canto.org/297530> |
| 80612 | *Vireo vicinior* | Scott Olmstead | <https://www.xeno-canto.org/80612> |
| 179357 | *Vireo vicinior* | Tim Marquardt | <https://www.xeno-canto.org/179357> |
| 282205 | *Vireo vicinior* | Paul Marvin | <https://www.xeno-canto.org/282205> |
| 282192 | *Vireo vicinior* | Paul Marvin | <https://www.xeno-canto.org/282192> |
| 282184 | *Vireo vicinior* | Paul Marvin | <https://www.xeno-canto.org/282184> |
| 334409 | *Vireo vicinior* | Scott Gravette | <https://www.xeno-canto.org/334409> |
| 34881 | *Vireo bellii* | Andrew Spencer | <https://www.xeno-canto.org/34881> |
| 109424 | *Vireo bellii* | Paul Marvin | <https://www.xeno-canto.org/109424> |
| 34881 | *Vireo bellii* | Andrew Spencer | <https://www.xeno-canto.org/34881> |
| 34315 | *Vireo bellii* | Andrew Spencer | <https://www.xeno-canto.org/34315> |
| 160976 | *Vireo atricapilla* | Paul Marvin | <https://www.xeno-canto.org/160976> |
| 152882 | *Vireo atricapilla* | Andrew Spencer | <https://www.xeno-canto.org/152882> |
| 34844 | *Vireo atricapilla* | Andrew Spencer | <https://www.xeno-canto.org/34844> |
| 21738 | *Vireo atricapilla* | Chris Parrish | <https://www.xeno-canto.org/21738> |
| 133059 | *Vireo atricapilla* | Andrew Spencer | <https://www.xeno-canto.org/133059> |
| 105703 | *Vireo atricapilla* | Nathan Pieplow | <https://www.xeno-canto.org/105703> |
| 145771 | *Vireo altiloquus* | Paul Marvin | <https://www.xeno-canto.org/145771> |
| 173666 | *Vireo altiloquus* | Eric DeFonso | <https://www.xeno-canto.org/173666> |
| 104635 | *Vireo altiloquus* | Andrew Spencer | <https://www.xeno-canto.org/104635> |
| 308673 | *Vireo altiloquus* | Ross Gallardy | <https://www.xeno-canto.org/308673> |
| 196029 | *Vireo altiloquus* | Paul Marvin | <https://www.xeno-canto.org/196029> |
| 213218 | *Vireo altiloquus* | Albert Lastukhin & Max Lastukhin | <https://www.xeno-canto.org/213218> |
| 252426 | *Vireo flavoviridis* | Dan Lane | <https://www.xeno-canto.org/252426> |
| 334255 | *Vireo flavoviridis* | Jeff Norris | <https://www.xeno-canto.org/334255> |
| 28547 | *Vireo flavoviridis* | Daniel Lane | <https://www.xeno-canto.org/28547> |
| 252424 | *Vireo flavoviridis* | Dan Lane | <https://www.xeno-canto.org/252424> |
| 359505 | *Aphelocoma insularis* | Paul Marvin | <https://www.xeno-canto.org/359505> |
| 358426 | *Aphelocoma insularis* | Paul Marvin | <https://www.xeno-canto.org/358426> |
| 360967 | *Aphelocoma californica* | Paul Marvin | <https://www.xeno-canto.org/360967> |
| 297455 | *Aphelocoma californica* | Ross Gallardy | <https://www.xeno-canto.org/297455> |
| 349731 | *Aphelocoma californica* | Paul Marvin | <https://www.xeno-canto.org/349731> |

### Table S3. Acoustic parameters extracted using the R package warbleR or calculated with R. Most variables were transformed prior to phylogenetic principal component analysis to be approximately normally distributed.

| Acoustic Parameters | Transformations |
| --- | --- |
| Duration | log(x + 0.01) |
| Mean frequency | log(x + 0.01) |
| Standard deviation of frequency | log(x + 0.01) |
| Median frequency | log(x + 0.01) |
| First quartile frequency | log(x + 0.01) |
| Third quartile frequency | log(x + 0.01) |
| Frequency interquartile range | log(x + 0.01) |
| Median time (time at which the signal is divided in two time intervals of equal energy) | log(x + 0.01) |
| First quartile time | log(x + 0.01) |
| Third quartile time | log(x + 0.01) |
| Time interquartile range | log(x + 0.01) |
| Spectral skewness | reciprocal |
| Spectral kurtosis (peakedness) | log(x + 0.01) |
| Spectral entropy (energy distribution of the frequency spectrum) |  |
| Time entropy (energy distribution on the time envelope) |  |
| Product of time and spectral entropy |  |
| Spectral flatness | log(x + 0.01) |
| Average of fundamental frequency | log(x + 0.01) |
| Minimum fundamental frequency | log(x + 0.01) |
| Average dominant frequency | log(x + 0.01) |
| Maximum of dominant frequency | sqrt |
| Minimum dominant frequency | sqrt |
| Range of dominant frequency measured across the acoustic signal | sqrt |
| Modulation index | log(x + 0.01) |
| Dominant frequency measurement at the start of the signal | sqrt |
| Dominant frequency measurement at the end of the signal | sqrt |
| Slope of the change in dominant frequency through time |  |
| Number of notes | log(x + 0.01) |
| Note rate | log(x + 0.01) |
| Length of longest note | log(x + 0.01) |
| Percentage of the song consisting of notes |  |
| Average note length | log(x + 0.01) |
| Average pause length | log(x + 0.01) |
| Length of longest pause | log(x + 0.01) |

### Table S4. Generalized linear model of ecological predictors of interspecific territoriality.

| Variable | Estimate | SE | *z* | *P* |
| --- | --- | --- | --- | --- |
| (Intercept) | 0.34 | 1.45 | 0.23 | 0.816 |
| Syntopy | 0.078 | 0.33 | 0.24 | 0.811 |
| Hybridization | 0.80 | 0.76 | 1.06 | 0.288 |
| Plumage dissimilarity | 0.12 | 0.34 | 0.34 | 0.734 |
| Song dissimilarity (pPCA) | -0.37 | 0.39 | -0.96 | 0.336 |
| Song similarity (SPCC) | 0.14 | 0.44 | 0.32 | 0.749 |
| Mass difference | -0.23 | 0.36 | -0.62 | 0.534 |
| Bill length difference | 0.014 | 0.40 | 0.036 | 0.917 |
| Guild overlap | -0.37 | 0.54 | -0.69 | 0.490 |

### Table S5. Generalized linear model of ecological predictors of interspecific territoriality, controlling for patristic distance.

| Variable | Estimate | SE | *z* | *P* |
| --- | --- | --- | --- | --- |
| (Intercept) | -0.07 | 1.54 | -0.04 | 0.965 |
| Syntopy | 0.02 | 0.34 | 0.06 | 0.954 |
| Hybridization | 0.55 | 0.81 | 0.68 | 0.499 |
| Patristic distance | -0.76 | 0.46 | -1.65 | 0.098 |
| Plumage dissimilarity | 0.29 | 0.36 | 0.82 | 0.413 |
| Song dissimilarity (pPCA) | -0.08 | 0.43 | -0.20 | 0.842 |
| Song similarity (SPCC) | 0.16 | 0.45 | 0.36 | 0.721 |
| Mass difference | -0.40 | 0.39 | -1.03 | 0.305 |
| Bill length difference | 0.29 | 0.44 | 0.65 | 0.513 |
| Guild overlap | -0.15 | 0.58 | -0.26 | 0.791 |

### Table S6. Generalized linear model predicting interspecific territoriality, with an interaction between hybridization and syntopy, and controlling for patristic distance.

| Variable | Estimate | SE | *z* | *P* |
| --- | --- | --- | --- | --- |
| (Intercept) | -0.84 | 4.00 | -0.21 | 0.833 |
| Syntopy | 5.71 | 2.87 | 1.99 | 0.047* |
| Hybridization | 3.52 | 2.06 | 1.71 | 0.088 |
| Patristic distance | -0.99 | 0.64 | -1.55 | 0.121 |
| Plumage dissimilarity | 0.62 | 0.55 | 1.13 | 0.26 |
| Song dissimilarity (pPCA) | -0.02 | 0.75 | -0.02 | 0.983 |
| Song similarity (SPCC) | -0.81 | 0.69 | -1.17 | 0.242 |
| Mass difference | -0.37 | 0.45 | -0.82 | 0.412 |
| Bill length difference | 0.19 | 0.62 | 0.31 | 0.754 |
| Guild overlap | -0.89 | 1.18 | -0.75 | 0.455 |
| Syntopy x hybridization | -6.63 | 2.96 | -2.24 | 0.025* |

Table S7. Generalized linear model predicting interspecific territoriality with interaction between syntopy and guild overlap.

| Variable | Estimate | SE | *z* | *P* |
| --- | --- | --- | --- | --- |
| (Intercept) | 0.02 | 1.59 | 0.01 | 0.989 |
| Syntopy | 2.08 | 1.59 | 1.31 | 0.190 |
| Guild overlap | -0.23 | 0.60 | -0.39 | 0.698 |
| Hybridization | 0.61 | 0.81 | 0.76 | 0.449 |
| Plumage dissimilarity | 0.06 | 0.35 | 0.16 | 0.870 |
| Song dissimilarity (pPCA) | -0.71 | 0.47 | -1.51 | 0.132 |
| Song similarity (SPCC) | 0.06 | 0.45 | 0.13 | 0.899 |
| Mass difference | -0.18 | 0.38 | -0.46 | 0.649 |
| Bill length difference | 0.002 | 0.40 | 0.005 | 0.996 |
| Syntopy x guild overlap | -0.79 | 0.58 | -1.37 | 0.172 |

### Table S8. Generalized linear model predicting interspecific territoriality, with an interaction between foraging guild and syntopy, and controlling for patristic distance.

| Variable | Estimate | SE | *z* | *P* |
| --- | --- | --- | --- | --- |
| (Intercept) | -0.26 | 1.70 | -0.15 | 0.878 |
| Syntopy | 2.03 | 1.70 | 1.19 | 0.234 |
| Guild overlap | -0.04 | 0.64 | -0.07 | 0.947 |
| Hybridization | 0.30 | 0.88 | 0.35 | 0.730 |
| Patristic distance | -0.73 | 0.47 | -1.56 | 0.119 |
| Plumage dissimilarity | 0.22 | 0.37 | 0.59 | 0.559 |
| Song dissimilarity (pPCA) | -0.40 | 0.51 | -0.80 | 0.425 |
| Song similarity (SPCC) | 0.12 | 0.46 | 0.25 | 0.802 |
| Mass difference | -0.31 | 0.41 | -0.76 | 0.447 |
| Bill length difference | 0.26 | 0.45 | 0.57 | 0.568 |
| Syntopy x guild overlap | -0.79 | 0.62 | -1.28 | 0.201 |

### Table S9. Generalized linear model predicting interspecific territoriality, with an interaction between the size principal component and syntopy, and controlling for patristic distance.

| Variable | Estimate | SE | *z* | *P* |
| --- | --- | --- | --- | --- |
| (Intercept) | 0.34 | 1.73 | 0.20 | 0.844 |
| Syntopy | -0.18 | 0.41 | -0.44 | 0.663 |
| Mass difference | -0.17 | 0.58 | -0.29 | 0.770 |
| Guild overlap | -0.34 | 0.65 | -0.52 | 0.600 |
| Hybridization | 0.41 | 0.94 | 0.44 | 0.662 |
| Patristic distance | -0.72 | 0.52 | -1.39 | 0.163 |
| Plumage dissimilarity | 0.57 | 0.44 | 1.31 | 0.190 |
| Song dissimilarity (pPCA) | -0.22 | 0.47 | -0.48 | 0.630 |
| Song similarity (SPCC) | 0.22 | 0.50 | 0.44 | 0.663 |
| Bill length difference | 0.31 | 0.56 | 0.55 | 0.579 |
| Syntopy x mass difference | 1.81 | 0.81 | 2.25 | 0.025* |

### Table S10. AICc comparison of generalized linear models predicting interspecific territoriality, without and with controlling for sympatry. Models with and without patristic distance are compared.

|  | AICc value | |
| --- | --- | --- |
| Model | Without Sympatry | With Sympatry |
| No Interactions, with Patristic Distance | 74.84 | 72.63 |
| No Interactions, without Patristic Distance | 73.90 | 72.02 |
| Hybrid x Syntopy, with Patristic Distance | 59.32 | 60.92 |
| Hybrid x Syntopy, without Patristic Distance | 58.60 | 60.06 |
| Guild x Syntopy, with Patristic Distance | 73.96 | 72.52 |
| Guild x Syntopy, without Patristic Distance | 73.33 | 72.00 |
| Size PC x Syntopy, with Patristic Distance | 67.16 | 65.70 |
| Size PC x Syntopy, without Patristic Distance | 67.12 | 66.21 |

### Table S11. Generalized linear models predicting continuous breeding range overlap.

| Model | Variable | Estimate | SE | *t* | *P* |
| --- | --- | --- | --- | --- | --- |
| 1. AICc = -17.78 | Patristic distance | 0.026 | 0.19 | 0.14 | 0.89 |
| 2. AICc = -15.74 | Patristic distance | 0.068 | 0.20 | 0.34 | 0.74 |
|  | I.T. | 0.25 | 0.40 | 0.61 | 0.54 |
| 3. AICc = -16.23 | Patristic distance | -0.053 | 0.21 | -0.25 | 0.80 |
|  | Hybridization | -0.39 | 0.42 | -0.93 | 0.35 |
| 4. AICc = -14.12 | Patristic distance | -0.011 | 0.22 | -0.052 | 0.96 |
|  | I.T. | 0.26 | 0.40 | 0.65 | 0.52 |
|  | Hybridization | -0.40 | 0.42 | -0.95 | 0.34 |
| 5. AICc = -16.66 | I.T. | 0.27 | 0.39 | 0.69 | 0.49 |
|  | Hybridization | -0.39 | 0.39 | -1.00 | 0.32 |
| 6. AICc = -14.76 | Patristic distance | -0.013 | 0.21 | -0.063 | 0.95 |
|  | I.T. | 1.12 | 0.62 | 1.82 | 0.069 |
|  | Hybridization | 0.22 | 0.53 | 0.42 | 0.68 |
|  | I.T. x Hybridization | -1.43 | 0.78 | -2.2 | 0.068 |
| 7. AICc = -17.43 | I.T. | 1.13 | 0.61 | 1.85 | 0.064 |
|  | Hybridization | 0.23 | 0.51 | 0.45 | 0.65 |
|  | I.T. x Hybridization | -1.43 | 0.78 | -1.82 | 0.068 |

### Table S12. Model Comparison where parapatry-sympatry cutoff is 20%. For each of three models (see Supplement 6), model parameters are calculated without (Null) the covariate or with a covariate (I.T., Hybridization, or I.T.*Hybridization).

|  | Model 1 | | | |  | Model 2 | | | |  | Model 3 | | | |
| --- | --- | --- | --- | --- | --- | --- | --- | --- | --- | --- | --- | --- | --- | --- |
|  | Null | IT | Hybr. | IT*H |  | Null | IT | Hybr. | IT*H |  | Null | IT | Hybr. | IT*H |
| ln.L | -25.0 | -23.0 | -24.5 | -22.1 |  | -25.0 | -23.0 | -24.5 | -22.1 |  | -25.0 | -23.0 | -23.9 | -22.1 |
| AICc | 52.15 | 50.28 | 53.32 | **48.57** |  | 54.34 | 54.94 | 58.04 | 53.29 |  | 56.64 | 60.10 | 61.95 | 58.50 |
| n_0_ | 45.00 | 24.00 | 18.00 | 30.00 |  | 45.00 | 24.00 | 18.00 | 30.00 |  | 45.00 | 24.00 | 18.00 | 30.00 |
| n_1_ | NA | 21.00 | 27.00 | 15.00 |  | NA | 21.00 | 27.00 | 15.00 |  | NA | 21.00 | 27.00 | 15.00 |
| θ_0_ | 0.76 | 0.87 | 0.83 | 0.87 |  | NA | NA | NA | NA |  | NA | NA | NA | NA |
| θ_0_ | NA | 0.62 | 0.70 | 0.53 |  | NA | NA | NA | NA |  | NA | NA | NA | NA |
| a_0_ | NA | NA | NA | NA |  | 0.76 | 0.87 | 0.83 | 0.87 |  | 0.76 | 0.88 | 1.00 | 0.87 |
| a_1_ | NA | NA | NA | NA |  | NA | 0.63 | 0.70 | 0.53 |  | NA | 0.63 | 0.70 | 0.54 |
| k_0_ | NA | NA | NA | NA |  | 33.32 | 46.09 | 12.27 | 35.46 |  | 59.92 | 39.02 | 0.31 | 60.90 |
| k_1_ | NA | NA | NA | NA |  | NA | 4.70 | 43.22 | 38.05 |  | NA | 10.90 | 48.08 | 11.01 |
| w_0_ | NA | NA | NA | NA |  | NA | NA | NA | NA |  | 0.08 | 0.00 | 0.00 | 0.02 |
| w_1_ | NA | NA | NA | NA |  | NA | NA | NA | NA |  | NA | 0.36 | 0.00 | 0.21 |
| conv | 0.00 | 0.00 | 0.00 | 0.00 |  | 0.00 | 0.00 | 0.00 | 0.00 |  | 0.00 | 0.00 | 0.00 | 0.00 |
| χ | NA | 4.06 | 1.02 | 5.77 |  | 0.00 | 4.12 | 1.02 | 5.77 |  | 0.00 | 4.16 | 2.32 | 5.77 |
| P | NA | 0.00 | 0.00 | 0.00 |  | 1.00 | 0.04 | 0.31 | 0.02 |  | 1.00 | 0.12 | 0.31 | 0.06 |

### Table S13. Model Comparison where parapatry-sympatry cutoff is 25%. For each of three models (see Supplement 6), model parameters are calculated without (Null) the covariate or with a covariate (I.T., Hybridization, or I.T.*Hybridization).

|  | Model 1 | | | |  | Model 2 | | | |  | Model 3 | | | |
| --- | --- | --- | --- | --- | --- | --- | --- | --- | --- | --- | --- | --- | --- | --- |
|  | Null | IT | Hybr. | IT*H |  | Null | IT | Hybr. | IT*H |  | Null | IT | Hybr. | IT*H |
| ln.L | -27.1 | -25.2 | -26.7 | -23.9 |  | -27.1 | -25.1 | -26.7 | -23.9 |  | -27.1 | -25.1 | -26.7 | -23.9 |
| AICc | 56.20 | 54.59 | 57.73 | **52.05** |  | 58.39 | 59.20 | 62.44 | 56.76 |  | 60.69 | 64.40 | 67.65 | 61.97 |
| n_0_ | 45.00 | 24.00 | 18.00 | 30.00 |  | 45.00 | 24.00 | 18.00 | 30.00 |  | 45.00 | 24.00 | 18.00 | 30.00 |
| n_1_ | NA | 21.00 | 27.00 | 15.00 |  | NA | 21.00 | 27.00 | 15.00 |  | NA | 21.00 | 27.00 | 15.00 |
| θ_0_ | 0.71 | 0.83 | 0.78 | 0.83 |  | NA | NA | NA | NA |  | NA | NA | NA | NA |
| θ_0_ | NA | 0.57 | 0.67 | 0.47 |  | NA | NA | NA | NA |  | NA | NA | NA | NA |
| a_0_ | NA | NA | NA | NA |  | 0.71 | 0.83 | 0.78 | 0.83 |  | 0.71 | 0.83 | 0.78 | 0.83 |
| a_1_ | NA | NA | NA | NA |  | NA | 0.59 | 0.67 | 0.47 |  | NA | 0.59 | 0.67 | 0.47 |
| k_0_ | NA | NA | NA | NA |  | 42.27 | 38.22 | 19.72 | 51.67 |  | 300.70 | 55.36 | 12.75 | 43.11 |
| k_1_ | NA | NA | NA | NA |  | NA | 3.04 | 112.8 | 33.58 |  | NA | 3.01 | 41.06 | 36.06 |
| w_0_ | NA | NA | NA | NA |  | NA | NA | NA | NA |  | 0.00 | 0.01 | 0.04 | 0.01 |
| w_1_ | NA | NA | NA | NA |  | NA | NA | NA | NA |  | NA | 0.08 | 0.00 | 0.00 |
| conv | 0.00 | 0.00 | 0.00 | 0.00 |  | 0.00 | 0.00 | 0.00 | 0.00 |  | 0.00 | 0.00 | 0.00 | 0.00 |
| χ | NA | 3.79 | 0.66 | 6.34 |  | 0.00 | 3.90 | 0.66 | 6.34 |  | 0.00 | 3.91 | 0.66 | 6.34 |
| P | NA | 0.00 | 0.00 | 0.00 |  | 1.00 | 0.05 | 0.42 | 0.01 |  | 1.00 | 0.14 | 0.72 | 0.04 |

### Table S14. Model Comparison where parapatry-sympatry cutoff is 30%. For each of three models (see Supplement 6), model parameters are calculated without (Null) the covariate or with a covariate (I.T., Hybridization, or I.T.*Hybridization).

|  | Model 1 | | | |  | Model 2 | | | |  | Model 3 | | | |
| --- | --- | --- | --- | --- | --- | --- | --- | --- | --- | --- | --- | --- | --- | --- |
|  | Null | IT | Hybr. | IT*H |  | Null | IT | Hybr. | IT*H |  | Null | IT | Hybr. | IT*H |
| ln.L | -29.3 | -26.8 | -29.3 | -26.4 |  | -29.2 | -26.2 | -29.2 | -26.0 |  | -29.2 | -24.7 | -29.1 | -24.6 |
| AICc | 60.67 | 57.91 | 62.79 | **57.07** |  | 62.64 | 61.46 | 67.32 | 61.06 |  | 64.91 | 63.50 | 72.49 | 63.44 |
| n_0_ | 45.00 | 24.00 | 18.00 | 30.00 |  | 45.00 | 24.00 | 18.00 | 30.00 |  | 45.00 | 24.00 | 18.00 | 30.00 |
| n_1_ | NA | 21.00 | 27.00 | 15.00 |  | NA | 21.00 | 27.00 | 15.00 |  | NA | 21.00 | 27.00 | 15.00 |
| θ_0_ | 0.64 | 0.79 | 0.67 | 0.77 |  | NA | NA | NA | NA |  | NA | NA | NA | NA |
| θ_0_ | NA | 0.48 | 0.63 | 0.40 |  | NA | NA | NA | NA |  | NA | NA | NA | NA |
| a_0_ | NA | NA | NA | NA |  | 0.66 | 0.79 | 0.67 | 0.77 |  | 0.66 | 0.79 | 0.67 | 0.77 |
| a_1_ | NA | NA | NA | NA |  | NA | 0.53 | 0.66 | 0.45 |  | NA | 0.56 | 0.66 | 0.50 |
| k_0_ | NA | NA | NA | NA |  | 3.45 | 45.22 | 10.93 | 41.48 |  | 3.58 | 46.68 | 11.87 | 72.37 |
| k_1_ | NA | NA | NA | NA |  | NA | 1.54 | 3.30 | 1.77 |  | NA | 1432.59 | 2.87 | 711.97 |
| w_0_ | NA | NA | NA | NA |  | NA | NA | NA | NA |  | 0.09 | 0.01 | 0.06 | 0.07 |
| w_1_ | NA | NA | NA | NA |  | NA | NA | NA | NA |  | NA | 0.98 | 0.00 | 0.96 |
| conv | 0.00 | 0.00 | 0.00 | 0.00 |  | 0.00 | 0.00 | 0.00 | 0.00 |  | 0.00 | 0.00 | 0.00 | 0.00 |
| χ | NA | 4.95 | 0.06 | 5.79 |  | 0.22 | 6.11 | 0.25 | 6.52 |  | 0.25 | 9.28 | 0.29 | 9.34 |
| P | NA | 0.00 | 0.00 | 0.00 |  | 0.64 | 0.01 | 0.62 | 0.01 |  | 0.88 | 0.01 | 0.87 | 0.01 |

### Table S15. Model Comparison where parapatry-sympatry cutoff is 35%, 40%, or 45% (results are identical for all three cutoffs). For each of three models (see Supplement 6), model parameters are calculated without (Null) the covariate or with a covariate (I.T., Hybridization, or I.T.*Hybridization).

|  | Model 1 | | | |  | Model 2 | | | |  | Model 3 | | | |
| --- | --- | --- | --- | --- | --- | --- | --- | --- | --- | --- | --- | --- | --- | --- |
|  | Null | IT | Hybr. | IT*H |  | Null | IT | Hybr. | IT*H |  | Null | IT | Hybr. | IT*H |
| ln.L | -29.8 | -28.0 | -29.8 | -27.5 |  | -29.8 | -27.4 | -29.7 | -27.1 |  | -29.8 | -25.9 | -29.7 | -25.7 |
| AICc | 61.76 | 60.34 | 63.94 | **59.27** |  | 63.84 | 63.89 | 68.47 | 63.26 |  | 66.12 | 65.93 | 73.64 | 65.64 |
| n_0_ | 45.00 | 24.00 | 18.00 | 30.00 |  | 45.00 | 24.00 | 18.00 | 30.00 |  | 45.00 | 24.00 | 18.00 | 30.00 |
| n_1_ | NA | 21.00 | 27.00 | 15.00 |  | NA | 21.00 | 27.00 | 15.00 |  | NA | 21.00 | 27.00 | 15.00 |
| θ_0_ | 0.62 | 0.75 | 0.61 | 0.73 |  | NA | NA | NA | NA |  | NA | NA | NA | NA |
| θ_0_ | NA | 0.48 | 0.63 | 0.40 |  | NA | NA | NA | NA |  | NA | NA | NA | NA |
| a_0_ | NA | NA | NA | NA |  | 0.63 | 0.75 | 0.61 | 0.73 |  | 0.63 | 0.75 | 0.61 | 0.73 |
| a_1_ | NA | NA | NA | NA |  | NA | 0.53 | 0.66 | 0.45 |  | NA | 0.56 | 0.66 | 0.50 |
| k_0_ | NA | NA | NA | NA |  | 3.93 | 54.48 | 17.41 | 52.20 |  | 3.78 | 49.35 | 11.20 | 58.31 |
| k_1_ | NA | NA | NA | NA |  | NA | 1.54 | 3.30 | 1.77 |  | NA | 1432.59 | 2.87 | 711.97 |
| w_0_ | NA | NA | NA | NA |  | NA | NA | NA | NA |  | 0.03 | 0.06 | 0.01 | 0.04 |
| w_1_ | NA | NA | NA | NA |  | NA | NA | NA | NA |  | NA | 0.98 | 0.00 | 0.96 |
| conv | 0.00 | 0.00 | 0.00 | 0.00 |  | 0.00 | 0.00 | 0.00 | 0.00 |  | 0.00 | 0.00 | 0.00 | 0.00 |
| χ | NA | 3.61 | 0.02 | 4.68 |  | 0.11 | 4.78 | 0.20 | 5.41 |  | 0.13 | 7.94 | 0.24 | 8.24 |
| P | NA | 0.00 | 0.00 | 0.00 |  | 0.74 | 0.03 | 0.65 | 0.02 |  | 0.93 | 0.02 | 0.89 | 0.02 |

### Table S16. Model Comparison where parapatry-sympatry cutoff is 50% or 55% (results are identical for both cutoffs). For each of three models (see Supplement 6), model parameters are calculated without (Null) the covariate or with a covariate (I.T., Hybridization, or I.T.*Hybridization).

|  | Model 1 | | | |  | Model 2 | | | |  | Model 3 | | | |
| --- | --- | --- | --- | --- | --- | --- | --- | --- | --- | --- | --- | --- | --- | --- |
|  | Null | IT | Hybr. | IT*H |  | Null | IT | Hybr. | IT*H |  | Null | IT | Hybr. | IT*H |
| ln.L | -30.6 | -29.8 | -30.6 | -29.2 |  | -30.4 | -29.2 | -30.2 | -28.8 |  | -30.3 | -27.6 | -29.3 | -27.4 |
| AICc | 63.38 | 63.90 | 65.51 | **62.67** |  | 64.98 | 67.45 | 69.34 | 66.65 |  | 67.16 | 69.49 | 72.76 | 69.04 |
| n_0_ | 45.00 | 24.00 | 18.00 | 30.00 |  | 45.00 | 24.00 | 18.00 | 30.00 |  | 45.00 | 24.00 | 18.00 | 30.00 |
| n_1_ | NA | 21.00 | 27.00 | 15.00 |  | NA | 21.00 | 27.00 | 15.00 |  | NA | 21.00 | 27.00 | 15.00 |
| θ_0_ | 0.58 | 0.67 | 0.56 | 0.67 |  | NA | NA | NA | NA |  | NA | NA | NA | NA |
| θ_0_ | NA | 0.48 | 0.59 | 0.40 |  | NA | NA | NA | NA |  | NA | NA | NA | NA |
| a_0_ | NA | NA | NA | NA |  | 0.61 | 0.67 | 0.56 | 0.67 |  | 0.61 | 0.67 | 0.59 | 0.67 |
| a_1_ | NA | NA | NA | NA |  | NA | 0.53 | 0.66 | 0.45 |  | NA | 0.56 | 0.66 | 0.50 |
| k_0_ | NA | NA | NA | NA |  | 2.14 | 31.91 | 1.90 | 37.86 |  | 2.35 | 57.45 | 104.6 | 329.86 |
| k_1_ | NA | NA | NA | NA |  | NA | 1.54 | 1.85 | 1.77 |  | NA | 1432.59 | 2.34 | 711.97 |
| w_0_ | NA | NA | NA | NA |  | NA | NA | NA | NA |  | 0.23 | 0.01 | 1.56 | 0.00 |
| w_1_ | NA | NA | NA | NA |  | NA | NA | NA | NA |  | NA | 0.98 | 0.33 | 0.96 |
| conv | 0.00 | 0.00 | 0.00 | 0.00 |  | 0.00 | 0.00 | 0.00 | 0.00 |  | 0.00 | 0.00 | 0.00 | 0.00 |
| χ | NA | 1.67 | 0.06 | 2.91 |  | 0.60 | 2.84 | 0.95 | 3.64 |  | 0.71 | 6.01 | 2.74 | 6.46 |
| P | NA | 0.00 | 0.00 | 0.00 |  | 0.44 | 0.09 | 0.33 | 0.06 |  | 0.70 | 0.05 | 0.25 | 0.04 |

### Table S17. Model Comparison where Covariate = IT and parapatry-sympatry cutoff is 60%. For each of three models (see Supplement 6), model parameters are calculated with (Cov) and without (Null) the covariate.

|  | Model 1 | | | |  | Model 2 | | | |  | Model 3 | | | |
| --- | --- | --- | --- | --- | --- | --- | --- | --- | --- | --- | --- | --- | --- | --- |
|  | Null | IT | Hybr. | IT*H |  | Null | IT | Hybr. | IT*H |  | Null | IT | Hybr. | IT*H |
| ln.L | -31.1 | -28.4 | -30.6 | -25.6 |  | -31.0 | -28.0 | -30.6 | -25.6 |  | -29.3 | -27.3 | -28.9 | -25.1 |
| AICc | 64.28 | 61.17 | 65.51 | **55.50** |  | 66.33 | 64.97 | 70.20 | 60.12 |  | 65.11 | 68.88 | 71.97 | 64.45 |
| n_0_ | 45.00 | 24.00 | 18.00 | 30.00 |  | 45.00 | 24.00 | 18.00 | 30.00 |  | 45.00 | 24.00 | 18.00 | 30.00 |
| n_1_ | NA | 21.00 | 27.00 | 15.00 |  | NA | 21.00 | 27.00 | 15.00 |  | NA | 21.00 | 27.00 | 15.00 |
| θ_0_ | 0.47 | 0.63 | 0.56 | 0.63 |  | NA | NA | NA | NA |  | NA | NA | NA | NA |
| θ_0_ | NA | 0.29 | 0.41 | 0.13 |  | NA | NA | NA | NA |  | NA | NA | NA | NA |
| a_0_ | NA | NA | NA | NA |  | 0.50 | 0.62 | 0.56 | 0.63 |  | 1.00 | 0.63 | 0.59 | 0.63 |
| a_1_ | NA | NA | NA | NA |  | NA | 0.38 | 0.41 | 0.14 |  | NA | 0.33 | 1.00 | 0.17 |
| k_0_ | NA | NA | NA | NA |  | 1.77 | 50.03 | 1.90 | 36.09 |  | 0.14 | 51.71 | 104.60 | 53.75 |
| k_1_ | NA | NA | NA | NA |  | NA | 0.74 | 48.37 | 2.54 |  | NA | 882.40 | 0.12 | 1.8e7 |
| w_0_ | NA | NA | NA | NA |  | NA | NA | NA | NA |  | 5.29 | 0.00 | 1.56 | 0.08 |
| w_1_ | NA | NA | NA | NA |  | NA | NA | NA | NA |  | NA | 0.97 | 6.70 | 0.97 |
| conv | 0.00 | 0.00 | 0.00 | 0.00 |  | 0.00 | 0.00 | 0.00 | 0.00 |  | 0.00 | 0.00 | 0.00 | 0.00 |
| χ | NA | 5.30 | 0.95 | 10.97 |  | 0.14 | 6.21 | 0.98 | 11.06 |  | 3.66 | 7.51 | 4.42 | 11.94 |
| P | NA | 0.00 | 0.00 | 0.00 |  | 0.71 | 0.01 | 0.32 | 0.00 |  | 0.16 | 0.02 | 0.11 | 0.00 |

### Table S18. Model Comparison where parapatry-sympatry cutoff is 65%. For each of three models (see Supplement 6), model parameters are calculated without (Null) the covariate or with a covariate (I.T., Hybridization, or I.T.*Hybridization).

|  | Model 1 | | | |  | Model 2 | | | |  | Model 3 | | | |
| --- | --- | --- | --- | --- | --- | --- | --- | --- | --- | --- | --- | --- | --- | --- |
|  | Null | IT | Hybr. | IT*H |  | Null | IT | Hybr. | IT*H |  | Null | IT | Hybr. | IT*H |
| ln.L | -29.8 | -29.1 | -29.6 | -26.7 |  | -29.8 | -28.7 | -29.6 | -26.6 |  | -28.4 | -27.4 | -27.8 | -26.2 |
| AICc | 61.76 | 62.52 | 63.39 | **57.65** |  | 63.95 | 66.32 | 68.10 | 62.28 |  | 63.46 | 68.98 | 69.81 | 66.61 |
| n_0_ | 45.00 | 24.00 | 18.00 | 30.00 |  | 45.00 | 24.00 | 18.00 | 30.00 |  | 45.00 | 24.00 | 18.00 | 30.00 |
| n_1_ | NA | 21.00 | 27.00 | 15.00 |  | NA | 21.00 | 27.00 | 15.00 |  | NA | 21.00 | 27.00 | 15.00 |
| θ_0_ | 0.38 | 0.46 | 0.44 | 0.50 |  | NA | NA | NA | NA |  | NA | NA | NA | NA |
| θ_0_ | NA | 0.29 | 0.33 | 0.13 |  | NA | NA | NA | NA |  | NA | NA | NA | NA |
| a_0_ | NA | NA | NA | NA |  | 0.38 | 0.46 | 0.44 | 0.50 |  | 1.00 | 1.00 | 0.47 | 0.50 |
| a_1_ | NA | NA | NA | NA |  | NA | 0.38 | 0.33 | 0.14 |  | NA | 0.33 | 1.00 | 0.17 |
| k_0_ | NA | NA | NA | NA |  | 41.39 | 43.35 | 3.43 | 139.5 |  | 0.11 | 0.08 | 403.4 | 51.57 |
| k_1_ | NA | NA | NA | NA |  | NA | 0.74 | 44.15 | 2.54 |  | NA | 882.40 | 0.13 | 1.8e7 |
| w_0_ | NA | NA | NA | NA |  | NA | NA | NA | NA |  | 9.21 | 8.49 | 1.48 | 0.00 |
| w_1_ | NA | NA | NA | NA |  | NA | NA | NA | NA |  | NA | 0.97 | 8.81 | 0.97 |
| conv | 0.00 | 0.00 | 0.00 | 0.00 |  | 0.00 | 0.00 | 0.00 | 0.00 |  | 0.00 | 0.00 | 0.00 | 0.00 |
| χ | NA | 1.44 | 0.56 | 6.30 |  | 0.00 | 2.34 | 0.57 | 6.39 |  | 2.79 | 4.90 | 4.07 | 7.26 |
| P | NA | 0.00 | 0.00 | 0.00 |  | 1.00 | 0.13 | 0.45 | 0.01 |  | 0.25 | 0.09 | 0.13 | 0.03 |

Table S19. ΔAICc values for Markov models of the probability of transitioning from parapatry to sympatry, with different covariates. This comparison was done for different cutoff values of percent breeding range overlap for designating parapatric or sympatry species pairs. Cutoffs 35%-45% and 50%-55% are combined because no species pairs had percent breeding range overlap values within those intervals.

|  | Parapatry-sympatry cutoff (% breeding range overlap) | | | | | | |
| --- | --- | --- | --- | --- | --- | --- | --- |
|  | 20% | 25% | 30% | 35% - 45% | 50% - 55% | 60% | 65% |
| No Covariate | **0.00** | **0.00** | 0.07 | 1.57 | 1.68 | 0.79 | **0.00** |
| I.T.* | 2.08 | 2.18 | 2.08 | 3.76 | 3.75 | 2.37 | 2.11 |
| Hybrid | 2.15 | 1.39 | **0.00** | **0.00** | **0.00** | 2.73 | 1.72 |
| I.T. x Hybrid | 1.63 | 1.90 | 2.11 | 3.75 | 3.85 | **0.00** | 1.16 |

*I.T. = interspecific territoriality
